# An evaluation of the efficacy, reliability, and sensitivity of motion correction strategies for resting-state functional MRI

**DOI:** 10.1101/156380

**Authors:** Linden Parkes, Ben Fulcher, Murat Yücel, Alex Fornito

## Abstract

Estimates of functional connectivity derived from resting-state functional magnetic resonance imaging (rs-fMRI) are sensitive to artefacts caused by in-scanner head motion. This susceptibility has motivated the development of numerous denoising methods designed to mitigate motion-related artefacts. Here, we compare popular retrospective rs-fMRI denoising methods, such as regression of head motion parameters and mean white matter (WM) and cerebrospinal fluid (CSF) (with and without expansion terms), aCompCor, volume censoring (e.g., scrubbing and spike regression), global signal regression and ICA-AROMA, combined into 19 different pipelines. These pipelines were evaluated across five different quality control benchmarks in four independent datasets associated with varying levels of motion. Pipelines were benchmarked by examining the residual relationship between in-scanner movement and functional connectivity after denoising; the effect of distance on this residual relationship; whole-brain differences in functional connectivity between high- and low-motion healthy controls (HC); the temporal degrees of freedom lost during denoising; and the test-retest reliability of functional connectivity estimates. We also compared the sensitivity of each pipeline to clinical differences in functional connectivity in independent samples of schizophrenia and obsessive-compulsive disorder. Our results indicate that (1) simple linear regression of regional fMRI time series against head motion parameters and WM/CSF signals (with or without expansion terms) is not sufficient to remove head motion artefacts; (2) aCompCor pipelines may only be viable in low-motion data; (3) volume censoring performs well at minimising motion-related artefact but a major benefit of this approach derives from the exclusion of high-motion individuals; (4) while not as effective as volume censoring, ICA-AROMA performed well across our benchmarks for relatively low cost in terms of data loss; and (5) group comparisons in functional connectivity between healthy controls and schizophrenia patients are highly dependent on preprocessing strategy. We offer some recommendations for best practice and outline some simple analyses to facilitate transparent reporting of the degree to which a given set of findings may be affected by motion-related artefact.

## 1. Introduction

Fluctuations of the blood-oxygenation-level-dependent (BOLD) signal recorded with functional magnetic resonance imaging during task-free “resting state” experiments (rs-fMRI) are highly organized, being correlated across anatomically distributed networks (Fox and Raichle, 2007) that correspond to those typically co-activated during task performance (Smith et al., 2009). These spontaneous dynamics predict task-evoked activation and behaviour (Cole et al., 2014; Fox and Raichle, 2007; Fox et al., 2007), can be used to accurately identify individuals across repeated scans (Finn et al., 2015), and are under significant genetic control (Fornito et al., 2011; Glahn et al., 2010). These findings suggest that resting-state fMRI can be used to probe a functionally important aspect of intrinsic brain dynamics which, together with the relative ease of data acquisition, has made the technique an attractive phenotyping tool for studies of brain disease and at-risk populations (Dandash et al., 2014; Fornito and Bullmore, 2010; Fornito et al., 2013).

A major obstacle in the analysis of fMRI data, particularly those acquired during unconstrained resting-state conditions, is contamination of the BOLD signal by head motion and fluctuations in non-neuronal physiological processes. Head motion is a particularly pernicious problem. Even small movements of the head between volumes acquired during a scan will cause erroneous intensity changes in BOLD data that are not accounted for by volume realignment. For example, due to the fact that volumes are acquired over multiple slices, movement of the head inside the scanner’s reference frame can cause excitation of different slices at subsequent time points relative to previous ones. These so-called ‘spin history’ effects (Friston et al., 1996) lead to motion-related changes in signal intensity in the BOLD data that obfuscate measurement of localized haemodynamics. In turn, this contamination can influence estimates of functional connectivity – i.e., statistical estimates of pairwise time series covariation – such that increased motion inflates coupling between nearby brain regions (type 1 effects), inflates long-range coupling if the signal disruption is similar and widespread across the brain (type 2 effects), or can reduce coupling between regions if the disruption varies across regions (type 3 effects) (Power et al., 2015). The problem is especially pronounced in group comparisons (e.g., between patients and controls), where differences in head motion can introduce systematic bias to connectivity estimates.

The most commonly-used method for removing motion-related noise from BOLD signals is linear regression (Fox and Raichle, 2007). With this approach, voxel-wise BOLD time series are regressed against head motion time series estimated along six dimensions (i.e., translational displacements along the *X*-, *Y*-, and *Z*-axes, and rotational displacements of pitch, roll, and yaw; hereafter referred to as head motion parameters, or HMP). To control for fluctuations in non-neuronal physiology, it is also common to regress voxel-averaged time series extracted from tissue compartments thought to contain nuisance signals, such as white matter (WM) and cerebrospinal fluid (CSF) (Fox and Raichle, 2007). The residuals of this confound regression are then used for further analysis. Expansion terms can also be added to the model to account for residual variance not removed by first-order effects. For example, Friston et al. (1996) recommended an autoregressive model that also included the motion estimates from the previous time point, as well as square terms (Friston-24 model; *M*_*t*_, *M*_*t-1*_, 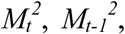 where *M* is the motion time series for a given dimension and *t* is time). Other researchers have used expanded models that incorporate temporal derivatives, calculated as backwards differences (Van Dijk et al., 2012), or have included both temporal derivatives and square terms (Satterthwaite et al., 2013).

Another common yet controversial method for correcting for physiological noise and head motion is global signal regression (GSR). GSR corrects for covariance between voxel-wise BOLD signals and the mean BOLD signal averaged across all voxels. GSR has been shown to reduce non-neuronal sources of physiological variance in the BOLD signal, such as those linked to respiration (Birn, 2012; Power et al., 2017b), and to mitigate the effects of in-scanner movement (Power et al., 2014; Yan et al., 2013a). However, GSR may also remove BOLD signal fluctuations of neuronal origin (Chen et al., 2012), spuriously weakening some correlations. The method also changes the distribution of functional connectivity estimates in the brain so that it is approximately centred on zero, which causes the emergence of negative correlations which may be artefactual (Fox et al., 2009; Murphy et al., 2009). There is also evidence that GSR can drive artefactual group differences in functional connectivity (Gotts et al., 2013; Saad et al., 2012). Critically, the extent to which GSR removes noise or signal from BOLD data may be contingent on the amount of global noise present (Chen et al., 2012), which further complicates group comparisons since group differences in head motion (and potentially, non-neuronal physiology) will cause differences in noise levels and thus lead to GSR exerting a differential effect on BOLD time series.

Using the various covariates described above in a regression model can reduce noise in BOLD data, but the overall effects of subtle in-scanner movements from volume-to-volume, called framewise displacements (FDs), are not fully removed by this approach (Power et al., 2012; Satterthwaite et al., 2012; Van Dijk et al., 2012). One solution, proposed by Power and colleagues (Power et al., 2015; 2014; 2012; 2013), is called “scrubbing”, and involves removing data points (volumes) for which the FD exceeds some threshold. A related strategy is called spike regression (Lemieux et al., 2007; Satterthwaite et al., 2013), which involves modelling the influence of contaminated time points using separate delta functions, one for each contaminated time point, and removing these effects via linear regression. Several groups have shown that these methods can effectively mitigate the impact of FDs on functional connectivity estimates (Power et al., 2012; Satterthwaite et al., 2013; Yan et al., 2013a), yet they come at the cost of potentially large amounts of lost data.

To get around the limitations of volume censoring, several alternative, data-driven methods have emerged that can reduce noise without data censoring (Behzadi et al., 2007; Muschelli et al., 2014; Pruim et al., 2015b). One popular method is CompCor (Behzadi et al., 2007), in which BOLD time series from voxels presumed to sample non-neuronal physiology are summarised as temporal principal components and entered as nuisance parameters in a linear regression model. A popular application of this technique is known as anatomical CompCor (aCompCor; Muschelli et al., 2014), which uses noise-related principal components estimated from WM and CSF voxels time courses. Other recent approaches use automated selection of noise-related components from a spatial independent component analysis (ICA) of the data. For example, ICA-FIX selects noise components based on matches to a manually curated training set of noise components (Griffanti et al., 2014; Salimi-Khorshidi et al., 2014). More recently, a simpler method called ICA-AROMA, which does not rely on the establishment of a study-specific training dataset, has been developed for the automatic detection of motion-related components according to a specific set of *a priori* criteria (Pruim et al., 2015b; 2015a). Alternative, methods for noise correction that rely on wavelet decomposition of the BOLD signal itself have also been proposed (Patel et al., 2014).

Several recent evaluation studies have been performed to compare the relative performance of many of these BOLD denoising methods on one or more of several quality-control benchmarks. These benchmarks, and the acronyms used to identify them, are summarised in Table 1. Evaluations of different pipelines using various combinations of these benchmarks have generally failed to converge on a single approach that performs best across all outcome metrics. In general, it appears that, compared to simple first-order linear regression models (that include the 6 HMP as well as mean WM/CSF signals), models that include expansion terms perform better on HLM contrasts and motion-BOLD contrasts, but show reduced TRT (Satterthwaite et al., 2012; Yan et al., 2013a). Adding volume censoring to these expansion models yields further improvements to QC-FC correlations, QC-FC distance-dependence, HLM contrasts, and motion-BOLD contrasts (Ciric et al., 2017; Power et al., 2012; Satterthwaite et al., 2013; Yan et al., 2013a). Moreover, incorporating GSR leads to substantial improvements in motion-BOLD contrasts and QC-FC correlations (Ciric et al., 2017; Yan et al., 2013a), but exaggerates QC-FC distance-dependence (Ciric et al., 2017). One study found that expanded motion regression with aCompCor was effective at reducing FD-DVARS without GSR or scrubbing (Muschelli et al., 2014). Another study found that ICA-AROMA performed equivalently to expanded motion regression with scrubbing and outperformed expanded motion regression with aCompCor on HLM contrasts, while yielding reduced tDOF-loss (Pruim et al., 2015a). A subsequent comparison found that expanded regression models plus scrubbing outperformed both ICA-AROMA and aCompCor on QC-FC correlations, but that ICA-AROMA was the only method to show virtually no QC-FC distance-dependence (Ciric et al., 2017).

**Table 1.**
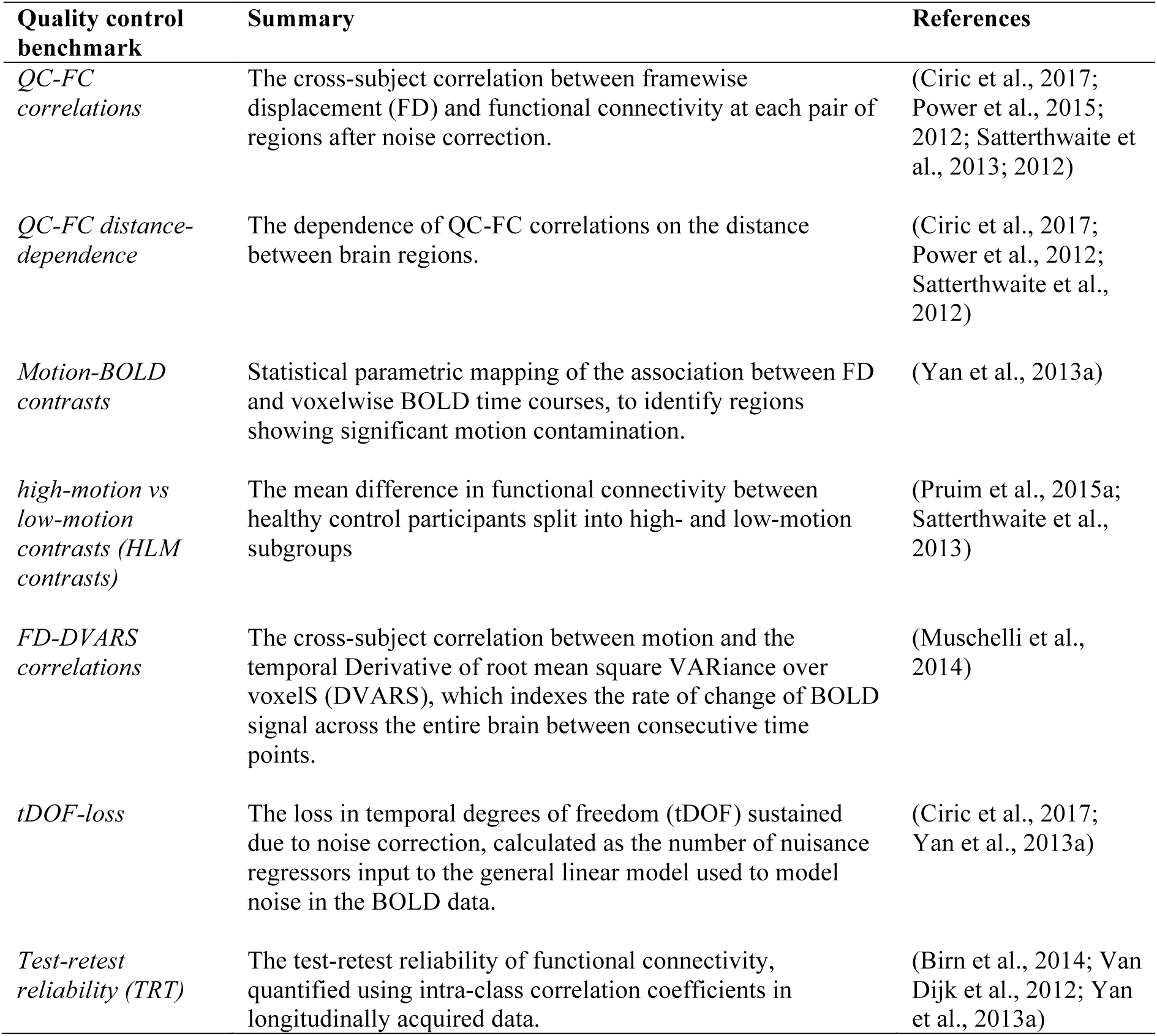
Summary of quality control metrics

| **Quality control benchmark** | **Summary** | **References** |
| --- | --- | --- |
| *QC-FC correlations* | The cross-subject correlation between framewise displacement (FD) and functional connectivity at each pair of regions after noise correction. | (Ciric et al., 2017; Power et al., 2015; 2012; Satterthwaite et al., 2013; 2012) |
| *QC-FC distance-dependence* | The dependence of QC-FC correlations on the distance between brain regions. | (Ciric et al., 2017; Power et al., 2012; Satterthwaite et al., 2012) |
| *Motion-BOLD contrasts* | Statistical parametric mapping of the association between FD and voxelwise BOLD time courses, to identify regions showing significant motion contamination. | (Yan et al., 2013a) |
| *high-motion vs low-motion contrasts (HLM contrasts)* | The mean difference in functional connectivity between healthy control participants split into high- and low-motion subgroups | (Pruim et al., 2015a; Satterthwaite et al., 2013) |
| *FD-DVARS correlations* | The cross-subject correlation between motion and the temporal Derivative of root mean square VARIance over voxels (DVARS), which indexes the rate of change of BOLD signal across the entire brain between consecutive time points. | (Muschelli et al., 2014) |
| *tDOF-loss* | The loss in temporal degrees of freedom (tDOF) sustained due to noise correction, calculated as the number of nuisance regressors input to the general linear model used to model noise in the BOLD data. | (Ciric et al., 2017; Yan et al., 2013a) |
| *Test-retest reliability (TRT)* | The test-retest reliability of functional connectivity, quantified using intra-class correlation coefficients in longitudinally acquired data. | (Birn et al., 2014; Van Dijk et al., 2012; Yan et al., 2013a) |

Together, these results suggest that it is difficult to find a single denoising method that performs well across all quality control measures, and that there is a general trade-off between adequately modelling the contributions of noise to the data and limiting the number of noise regressors to avoid over-fitting and/or severe tDOF-loss. However, variability across studies in terms of the benchmark measures that have been used makes it difficult to perform fair comparisons. Furthermore, none of the benchmarking studies reported thus far have examined how these noise correction strategies impact the analysis of functional connectivity differences between groups, which is a key application of rs-fMRI.

In this study, we evaluated 9 popular rs-fMRI denoising strategies, combined into 19 different pipelines, applied to four independent datasets with respect to five benchmarks: QC-FC correlations, QC-FC distance-dependence, HLM contrasts, tDOF-loss, and TRT benchmarks. Our first dataset contained a large sample of healthy control participants with low levels of motion and served as a well-powered dataset for the aforementioned benchmarks. We also examined the relative sensitivity of each method in uncovering clinical group differences in two separate case-control samples. One was characterized by relatively high levels of motion and comprised healthy controls and patients with schizophrenia. The other contained low levels of motion and comprised healthy controls and patients with obsessive-compulsive disorder (OCD). The fourth dataset contained longitudinal data on healthy controls only and was used to examine TRT.

Our comprehensive assessment revealed that no single pipeline is completely effective in mitigating the contaminating effects of motion on functional connectivity, regardless of the level of motion. The top performing pipelines were those that included ICA-AROMA or some form of volume censoring. The latter slightly outperformed the former, but ICA-AROMA incurred a lower cost in terms of data loss. We also show that group differences vary considerably depending on the specific processing pipeline used. Our work highlights the need for the comprehensive reporting of motion and its impact on functional connectivity in case-control rs-fMRI studies.

## 2. Materials and methods

### 2.1 Participants and data

The rs-fMRI data used in this study were drawn from four sources: (1) the Beijing Zang dataset (*Beijing*: 192 HCs. http://fcon_1000.projects.nitrc.org/fcpClassic/FcpTable.html); (2) the Brain & Mental Health laboratory dataset (*BMH*: 39 HCs and 34 OCD patients); (3) the Consortium for Neuropsychiatric Phenomics dataset (CNP: 121 HCs and 50 schizophrenia patients Poldrack et al., 2016); and (4) the New York University dataset (*NYU:* 29 HCs). The Beijing, BMH, CNP, and NYU datasets were used to compare the relative performance of the different denoising strategies for removing the effects of in-scanner movement. The BMH and CNP datasets were used to examine which denoising strategy is most sensitive to clinical group differences (relative to HCs) in two disorders: obsessive-compulsive disorder (OCD; provided by BMH) and schizophrenia (SCZ; provided by CNP). The (NYU) dataset (http://fcon_1000.projects.nitrc.org/indi/CoRR/html/), available through the Consortium for Reliability and Reproducibility (CoRR; Zuo et al., 2014), was used to examine within- and between-session test-retest reliability of functional connectivity estimates obtained after the application of each denoising approach.

The Beijing dataset was acquired on a Siemens TRIO 3T scanner. A T1-weighted MP-RAGE structural was obtained (TE = 3.39 ms, TR = 2.53 s, flip angle = 7°, 128 slices with 1.33x1x1 mm voxels). Resting state data was obtained using BOLD contrast sensitive gradient echoplanar imaging (EPI) (TE = 30 ms, TR = 2 s, flip angle = 90°, 225 volumes, 33 slices). The BMH dataset was acquired on a Siemens MAGNETOM Skyra 3T scanner. Details of the T1 scan are TE = 2.55 ms, TR = 1.52 s, flip angle = 9°, 208 slices with 1 mm isotropic voxels. Details of the EPI scan are TE = 30 ms, TR = 2.5 s, flip angle = 90°, 189 volumes, 44 slices. The CNP dataset was acquired on one of two Siemens Trio 3T scanners. Details of the T1 scan are TE = 24 ms, TR = 5 s, flip angle = 90°, 176 slices with 1 mm isotropic voxels). Details of the EPI scan are TE = 30 ms, TR = 2 s, flip angle = 90°, 152 volumes, 34 slices. The NYU dataset was acquired using a Siemens MAGNETOM Allegra 3T scanner. Details of the T1 scan are TE = 3.25 ms, TR = 2.53 s, flip angle = 7°, 128 slices with 1.3 x 1.0 x 1.3 mm voxels. Details of the EPI scan are TE = 15 ms, TR = 2 s, flip angle = 90°, 180 volumes, 33 slices.

### 2.2 Image processing

EPI and T1-weighted scans were processed using code developed in Matlab, which is freely available through GitHub (https://github.com/lindenmp/rs-fMRI).

#### 2.2.1 Structural image processing

Each participant’s T1-weighted high-resolution structural image was processed using the following steps: (1) removing the neck using FSL’s *robustfov*; (2) segmentation of the native T1-weighted image into WM, CSF, and grey matter (GM) probability maps using SPM8’s *New Segment* routine to allow for identification of WM/CSF voxels for use with some pipelines; (3) nonlinear spatial transform of the T1-weighted image to MNI space using Advanced Normalization Tools (ANTs; Avants et al., 2008) with default settings (using the *antsRegistrationSyN.sh* script); and (4) application of the nonlinear transforms derived from the previous step to the WM, CSF, and GM masks.

Recent work by Power et al. (2017) demonstrated that, without extensive erosion of WM and CSF masks, these signals can be correlated with the GM signal. In turn, this correlation can inadvertently lead to the WM/CSF signals behaving like GSR when used as nuisance regressors. To minimise this effect, we applied five erosion cycles to the WM mask and two erosion cycles to the CSF mask following extraction of the ventricles (i.e., masking out of the CSF surrounding the sulci/gyri of the cortex; see Figure S1) before spatial normalisation to MNI space (see Table S1). If the WM/CSF masks for a given participant contained fewer than 5 voxels following a given erosion cycle, we selected the previous cycle.

### 2.2.2 Core image processing

The functional data underwent a core, common set of processing steps both before and after each denoising method was applied. The core processing pipeline used before denoising included the following steps: (1) removal of the first four volumes of each acquisition; (2) slice-time correction implemented in SPM8; (3) two-pass realignment of all volumes to the first volume (first pass) and then to the mean volume (second pass) using SPM8; (4) co-registration of EPI data to the native, cropped, high-resolution structural image via rigid-body registration using ANTs; (5) application of the nonlinear transform derived from the T1-weighted image processing pipeline to the co-registered EPI data using ANTs; (6) linear detrending of the spatially normalized BOLD time series; and (7) intensity normalisation of the EPI data to mode 1000 units. Specific denoising pipelines were then applied at this stage. These are described below.

After the application of each specific denoising pipeline, additional core processing steps included bandpass filtering between 0.008 and 0.08 Hz using the fast Fourier transform, and spatial smoothing with an 6mm FWHM kernel (although an exception was made for ICA-AROMA, which requires smoothing prior to noise correction, as discussed below).

Recent work by Power et al. (2017a) suggests that performing realignment after processing steps that involve temporal interpolation of the BOLD data (e.g., slice-time correction) leads to underestimation of the motion parameters and less stringent motion correction. Thus, we also performed realignment on the BOLD data before it underwent slice-time correction but after the first four volumes were discarded (i.e., between steps 1 and 2 above) in order to acquire ‘raw’ motion parameters. These parameters were used in all subsequent motion correction and quality control analyses (see 2.4 Outcome Measures).

### 2.3 Denoising pipelines

Our primary aim was to comprehensively evaluate the performance of currently popular methods for correcting BOLD time series for the effects of head motion. To this end, we investigated several confound regression strategies and volume censoring methods. Some denoising pipelines involve a combination of different noise correction strategies. Table 2 outlines the 19 different pipelines analysed here, each containing a particular combination of noise correction methods. Subsequent sections describe each strategy separately.

**Table 2.** Characteristics of 19 denoising pipelines analysed here.

| Pipeline | Noise corrections methods | No. of regressors |
| --- | --- | --- |
| 1 | 6HMP | 6 |
| 2 | 6HMP+2Phys | 8 |
| 3 | 6HMP+2Phys+GSR | 9 |
| 4 | 24HMP | 24 |
| 5 | 24HMP+8Phys | 32 |
| 6 | 24HMP+8Phys+4GSR | 36 |
| 7 | 24HMP+aCompCor | 34 |
| 8 | 24HMP+aCompCor50 | 24+k |
| 9 | 24HMP+aCompCor+4GSR | 38 |
| 10 | 24HMP+aCompCor50+4GSR | 28+k |
| 11 | 12HMP+aCompCor (Muschelli et al., 2014) | 22 |
| 12 | 12HMP+aCompCor50 (Muschelli et al., 2014) | 12+k |
| 13 | ICA-AROMA+2Phys (Pruim et al., 2015b) | 2+k |
| 14 | ICA-AROMA+2P+GSR | 3+k |
| 15 | ICA-AROMA+8Phys | 8+k |
| 16 | ICA-AROMA+8P+4GSR | 12+k |
| 17 | 24HMP+8Phys+4GSR+SpikeReg | 36+k |
| 18 | 24HMP+8Phys+4GSR+JP12Scrub | 36+k |
| 19 | 24HMP+4Phys+2GSR+JP14Scrub | 30+k |

*Notes.* HMP, head motion parameters. Phys, average white matter (WM) and cerebrospinal fluid (CSF) signals. GSR, global signal regression. aCompCor, anatomical component correction using top the 5 components in each of WM and CSF compartments. aCompCor50, anatomical component correction using enough components to explain 50% of the variance in each of WM and CSF compartments. SpikeReg, spike regression. JP12Scrub, basic
scrubbing. JP14Scrub, optimized scrubbing. $k$ denotes an arbitrary number of additional regressors estimated automatically by the denoising method and which can vary from person to person.

#### 2.3.1 Regression of head motion parameters

For each participant, the two-pass realignment of the BOLD data prior to slice-time correction yielded six time-series describing in-scanner movement along six dimensions – three translational axes of *X*, *Y*, and *Z*, and three rotational axes of pitch, roll, and yaw. BOLD time series were regressed against these head motion parameters and various expansion terms. In the 6HMP model, only the original six head motion parameters were employed as covariates. The 12HMP model employed these 6 parameters, as well as expansion terms derived by computing the temporal derivatives of each parameter (calculated as first-order backwards differences in the head motion time series data). The 24HMP model employs these 12 parameters, as well as the squares of both the original and derivative time series (Satterthwaite et al., 2013).

#### 2.3.2 Regression of mean white matter and cerebrospinal fluid signals

A popular method for controlling for additional head motion effects beyond the 6HMP, 12HMP, and 24HMP models, in addition to capturing physiological fluctuations of non-neuronal origin, is to generate a representative time series from tissue compartments that do not include grey matter. This involves extracting an averaged time series from all WM voxels, and separately for all CSF voxels. To this end, we generated eroded WM and CSF tissue masks that resulted in conservative estimates of the WM and CSF volumes, ensuring minimal contamination with GM voxels (see *2.2.1 Structural image processing*). As above, the resulting averaged WM and CSF signals were then incorporated into denoising procedures either in their original form (2Phys), or along with their temporal derivatives (4Phys), or derivatives, squares, and squares of derivatives (8Phys).

#### 2.3.3 Global signal regression

A controversial step in fMRI denoising is global signal regression (GSR), which involves regressing voxel-wise fMRI time series against an averaged signal computed across the entire brain (Fox et al., 2007; Murphy et al., 2009). We calculated the global signal by averaging across all voxels in the BOLD data using participant-specific masks that covered the entire brain. These masks were generated by taking the union of two whole brain masks created using FSL’s *bet* function applied to the spatially normalised EPI and T1-weighted images created during pre-processing. As above, GSR was performed by either removing, via linear regression, just the global signal (GSR) or by also removing the temporal derivative (2GSR), or temporal derivative, square term, and square of the derivative (4GSR).

#### 2.3.4 aCompCor

A popular method for modelling noise in BOLD data is to apply temporal principal component analysis (PCA) to putative nuisance signals (Behzadi et al., 2007; Muschelli et al., 2014). This approach, most commonly embodied by the aCompCor procedure, involves extracting orthogonal components of temporal variance from voxel-wise time series for the WM and CSF tissue compartments. Compared to averaging across voxels, PCA offers the advantage of identifying multiple orthogonal sources of variance in the data, which may better characterise the noise signals present in the WM and CSF tissue compartments. Behzadi et al.’s implementation involves extracting components of temporal variance from a single mask that combines together voxels in the WM and CSF, whereas Muschelli et al.’s implementation conducts PCA on the WM and CSF masks separately. We adopt the latter implementation.

We defined WM and CSF masks as outlined above. PCA was performed in the time domain on the voxel time series separately for the WM and CSF masks. As per previous work by Muschelli et al. (2014), aCompCor was run using two models that differed in the number of principal components (PCs) extracted. In the aCompCor model, we extracted the leading 5 PCs for each tissue type, yielding 10 confound regressors. In the aCompCor50 models, we extracted the number of PCs that cumulatively explained at least 50% of the variance for each tissue type, yielding a variable number of confound regressors for each participant and each tissue type. The analysis of Muschelli et al. (2014) suggests that aCompCor50 outperforms aCompCor in terms of mitigating motion-related artefacts and increases the specificity of functional connectome estimations. However, aCompCor50 has the potential to cost many more temporal degrees of freedom (see *2.4.3 Loss of temporal degrees of freedom*).

#### 2.3.5 ICA-AROMA

Recently, an independent component analysis-based method for motion denoising called ICA-AROMA was introduced by Pruim et al. (2015b; 2015a). This method attempts to automatically identify and remove motion-related artefacts from BOLD data by using FSL’s MELODIC (Beckmann et al., 2005) to first decompose the BOLD data into spatially independent components (IC) before applying a predetermined, theoretically motivated classifier to identify ICs as noise or signal. Specifically, ICs were classified as motion-related if any of the following criteria were true: (1) more than 10% of IC voxels were located within CSF; (2) IC time series contained more than 35% high-frequency content, where high-frequency is defined as a fraction of the Nyquist frequency at which higher frequencies explain 50% of the total power between 0.01 Hz and the Nyquist frequency; and (3) ICs exceed a decision boundary from a two-dimensional linear discriminant analysis (LDA) classifier. For the classifier, a two-dimensional feature space was defined using (1) the proportion of IC voxels that overlapped the edges of the brain; and (2) the maximum absolute correlation between the IC time series and parameters in an expanded head motion model, which included 72 motion realignment parameters comprising the same as those in our 24HMP model plus 24 parameters from a single time-point back and forward. Pruim et al. then defined a hyperplane using an LDA classifier trained on manually labelled ICs (labelled as either ‘Motion’, ‘Resting State Network’, or ‘Other’) from 30 rs-fMRI datasets.

Like all the other methods mentioned thus far, ICA-AROMA is applied to each participant’s BOLD data separately. Due to the use of a person-specific classifier to set thresholds for detecting noise components, the number of confound regressors removed from the BOLD data can vary across participants. Unlike the other methods, ICA-AROMA requires data to be spatially smoothed before noise correction. As such, for all pipelines that included ICA-AROMA (see Table 2), spatial smoothing was performed immediately before noise correction rather than after bandpass filtering.

An alternative ICA-based approach is FMRIB’s ICA-based X-noiseifier (ICA-FIX) (Griffanti et al., 2014; Salimi-Khorshidi et al., 2014). ICA-FIX’s implementation involves a much more extensive set of features that requires re-training and manual re-labelling of noise components on each new dataset. Here we focus only on methods that can be applied automatically to rs-fMRI data with minimal user input.

#### 2.3.6 Volume censoring

Apart from the confound regression strategies outlined above, an additional method for addressing head motion confounds in BOLD data involves censoring contaminated data points (i.e., entire volumes). The two techniques investigated here are spike regression (Satterthwaite et al., 2013) and scrubbing (Power et al., 2014; 2012). We note that spike regression and scrubbing are functionally equivalent procedures (i.e., if the same volumes were removed performance would be equivalent) and that their differences lie only in how volumes are marked as contaminated and how contaminated volumes are censored. Previous research has found that both spike regression and scrubbing improve data denoising, as quantified by reduced QC-FC distance-dependence and HLM contrasts (Power et al., 2012; Satterthwaite et al., 2013).

For spike regression and scrubbing, volumes were marked as contaminated by thresholding a given participant’s FD. There are several different ways of calculating FD (see Yan et al., 2013a for a summary and definition of each). They are all highly inter-correlated with each other (e.g., *r* > 0.90) (Power et al., 2015). However, the scales of the measures differ; thus, the choice of an FD threshold depends on the specific method used to calculate FD. Here, we sought to maintain consistency with past work. We thus used different FD measures for spike regression and scrubbing, following the methodology of Satterthwaite et al. (2013) for the former and Power et al. (2013) for the latter. Specifically, for spike regression, we calculated FD using the root mean squared volume-to-volume displacement of all brain voxels measured from the six head motion parameters, a method sometimes called FD_Jenk_ (Jenkinson et al., 2002; Satterthwaite et al., 2013; Yan et al., 2013a). Based on previous work by Satterthwaite et al. (2013), volumes were marked as contaminated if FD_Jenk_ >0.25mm. Satterthwaite et al. (2013) demonstrated that a single spike regressor placed at the contaminated time point outperformed a box-car function (i.e., marking a contiguous set of time points as contaminated) that also covered proximal time points. Thus, for each contaminated volume, a separate nuisance regressor was generated that matched the length of the BOLD time-series data, containing a value of 1 at the location of the contaminated volume and 0 elsewhere. As with ICA-AROMA, the number of regressors generated with spike regression varies over participants.

We examined two variants of scrubbing, both of which were developed by Power and colleagues (Power et al., 2014; 2013; 2012). We refer to these two variants as ‘basic scrubbing’ and ‘optimized scrubbing’. For basic scrubbing, we calculated FD as the sum of absolute differences in volume-to-volume changes in the six head motion parameters, FD_Power_ (Power et al., 2012; Yan et al., 2013a). Volumes were marked as contaminated using a combination of FD_Power_ and BOLD data variance (DVARS). DVARS is an estimate of framewise changes in BOLD signal intensity, and is calculated as the root-mean-squared variance of the temporal derivative of all brain voxel time series (Power et al., 2012). Based on Power et al. (2013), volumes were marked as contaminated if either of the following were true: FD_Power_ >0.2mm or DVARS >0.3% (Power et al., 2013). Thus, immediately before statistical analysis, contaminated volumes were removed from the BOLD data.

Optimized scrubbing involved an iterative procedure, as per Power et al., (2014). Contaminated volumes were identified using the same FD_Power_/DVARS measures as above but the DVARS criterion was adjusted to >0.2% (FD_Power_ criteria was unchanged), and any uncontaminated segments of BOLD data with fewer than 5 contiguous volumes were also marked as contaminated. During detrending and nuisance regression, contaminated volumes were censored from the BOLD data so they did not influence the fit of the linear models. Prior to bandpass filtering, contaminated volumes were censored and replaced with surrogate data generated using the frequency characteristics of the uncontaminated BOLD data. Finally, as in basic scrubbing, contaminated volumes (now represented by surrogate data) were censored from the BOLD data immediately before statistical analysis.

Volume censoring involves a costly trade-off, such that every censored volume improves data quality at the expense of data quantity. Removing too many volumes can result in insufficient data to produce reliable estimates of functional connectivity. Consistent with previous literature, we excluded participants with <4 minutes of data after either spike regression or scrubbing (Satterthwaite et al., 2013; Van Dijk et al., 2012). The participants that were excluded as a consequence of this criterion varied across spike regression, basic scrubbing, and optimized scrubbing due to the differing ways volumes were flagged as contaminated.

#### 2.3.7 Network construction

After pre-processing, we generated functional connectivity networks using two independent parcellations, one containing 333 cortical regions (Gordon et al., 2016) and the other containing 264 cortical and subcortical regions (Power et al., 2011). For each region in each parcellation, we estimated the mean time series across all voxels comprising the region, multiplying each voxel time series by its grey matter probability first to weight the time series mean. We then estimated Pearson correlation coefficients between each pair of regional averaged time series. These correlations can be represented as a network, where edges connecting pairs of nodes (brain regions) represent correlation coefficients between resting state fMRI time series. These networks underwent Fisher’s *r*-to-*z* transformation to normalise the correlation distribution and facilitate group comparisons.

For the BMH data, seven ROIs from the Power parcellation were discarded from analyses due to poor overlap with participant EPI data (no ROIs were excluded for the Gordon parcellation). For the CNP dataset, six and eight ROIs from the Gordon and Power parcellations, respectively, were discarded. For the Beijing and NYU dataset, no ROIs were discarded from either the Gordon or Power parcellations. We found that our results were qualitatively very similar for both parcellations. We therefore only present results obtained with the Gordon parcellation.

### 2.4 Outcome measures

Ideally, noise correction should remove any statistical relationship between functional connectivity and in-scanner movement. A commonly used summary statistic for quantifying the degree of person-specific head motion is the temporal mean of their FD time series (mFD). Here we use FD_Jenk_ to calculate mFD (similar results are obtained using FD_Power_). This metric was used to evaluate pipeline efficacy by assessing: (1) how the correlation between mFD and functional connectivity (FC), computed across participants for each network edge, changes following denoising (QC-FC); and (2) how this relationship varies as a function of distance between nodes, given prior evidence that movement typically has a more pronounced effect on the QC-FC correlation for short-range connectivity (Power et al., 2015; 2012; Van Dijk et al., 2012). In addition, we examined four other performance metrics: (1) the change in temporal degrees of freedom caused by each denoising pipeline (tDOF-loss); (2) the sensitivity of each pipeline to differences in functional connectivity between high- and low-motion HCs; (3) the test-retest reliability of each pipeline (TRT); and (4) the sensitivity of each pipeline to clinical differences in functional connectivity. These benchmarks are outlined in more detail in the following.

#### 2.4.1 QC-FC correlations

We used Pearson correlations to quantify the association between subject-specific mFD and functional connectivity estimates for each edge, resulting in an edge-specific QC-FC value estimated across the entire sample. To measure the efficacy of each approach in removing motion-related variance, we compared the proportion of edges where this QC-FC correlation was statistically significant (*p*<0.05, uncorrected and FDR-corrected) as well as the median absolute QC-FC correlation after applying each denoising pipeline. Where applicable, we only retained the HC group when calculating QC-FC correlations to avoid biases introduced by including patients.

#### 2.4.2 QC-FC distance-dependence

In-scanner movement commonly inflates short-range functional connectivity relative to medium- and long-range connectivity (Power et al., 2015; 2012; Van Dijk et al., 2012). This spatial variation arises because each voxel is often more strongly contaminated by proximal (vs distal) voxels when the head moves in the scanner, thus spuriously increasing synchrony between nearby brain regions (type 1 effects). Successful noise correction should thus result in no distance-dependence of QC-FC correlations. It has previously been shown that ICA-AROMA is effective at mitigating this relationship while simple linear regression methods are not and GSR exacerbates it (Ciric et al., 2017). Here, we estimated the distance between regions as the Euclidean distance between the stereotaxic coordinates of the volumetric centres of brain region pairs. For each edge, we then quantified the association between this distance measure and the QC-FC correlation for that edge using Spearman’s rank correlation coefficient, *r*, due to the non-linearity of some associations. To visualise the relationship between QC-FC and distance, we also plotted the QC-FC correlation of each edge as a function of this distance metric. Since each network contained tens of thousands of edges, the resulting scatterplots displayed a dense point cloud which masked mean-level trends. To provide a more interpretable visualisation of QC-FC distance-dependence, we divided the data into 10 equiprobable bins (using equally spaced quantiles to define bins) based on nodal distance, and plotted the mean and standard deviation of QC-FC correlations in each bin. As above, where applicable, we only retained the HC group when calculating QC-FC distance-dependence to avoid biases introduced by including patients.

#### 2.4.3 Loss of temporal degrees of freedom

The pipelines examined here vary in terms of the number of regressors used to model noise in the fMRI time series. Using more nuisance regressors can capture additional sources of noise-related variance in the data and thus improve denoising, but this comes at the expense of a loss of temporal degrees of freedom. The number of time points in a BOLD dataset represents the degrees of freedom available for statistical inference, and fewer degrees of freedom can spuriously increase functional connectivity (Yan et al., 2013a). Furthermore, noise correction strategies that use variable degrees of freedom across participants (e.g., volume censoring and ICA-AROMA) can lead to artefactual group differences in functional connectivity (Pruim et al., 2015b; 2015a; Yan et al., 2013b). Thus, the relative performance of different denoising strategies must be balanced against lost temporal degrees of freedom (tDOF-loss). For each pipeline, we calculated tDOF-loss as the number of regressors and, in the case of volume censoring, the number of contaminated volumes. As above, where applicable, we only retained the HC group when calculating tDOF-loss.

#### 2.4.4 Group differences in high-motion and low-motion healthy participants

Apart from mitigating QC-FC correlations, successful noise correction should also yield minimal group differences in functional connectivity between HC participants that differ on their amount of in-scanner movement. Thus, for the BMH and CNP datasets, we split HC participants into three equally sized groups based on mFD and mapped group-differences in functional connectivity using mass-univariate statistics. Specifically, for each edge we calculated two-tailed *t*-contrasts assessing both increases and decreases in functional connectivity in high-motion HCs compared to low-motion HCs (HLM contrasts; medium-motion group was not included). Age (demeaned) and sex were entered as covariates, and in the case of the CNP dataset, scanner site was also included as a nuisance covariate. We then thresholded the *F*-contrasts at *p*<0.05 uncorrected (two-tailed) and *p*<0.05 FDR-corrected (two-tailed) and report the proportion of significant edges for each pipeline.

#### 2.4.5 Test-retest reliability

One might expect that a good denoising strategy will yield consistent and reliable estimates of functional connectivity across repeated measurements of the same subject under the same conditions. However, previous work has shown that the TRT reliability of functional connectivity decreases with increasing variance explained by denoising models (Birn et al., 2014; But see Van Dijk et al., 2012; Yan et al., 2013a). This may be due to the presence of reproducible motion artefact present in BOLD data itself (Yan et al., 2013a). To characterize the effects of noise correction pipelines on the reliability of functional connectivity estimates, we examined TRT reliability using the intra-class correlation (ICC) coefficient (Shrout and Fleiss, 1979) calculated on the NYU dataset, defined as

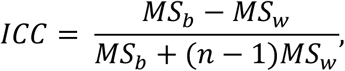

where *MS*_*b*_ is the between-subject mean square for each edge, *MS*_*w*_ is the within-subject mean square for each edge, and *n* is the number of observations per participant (here *n* is always to set to 2 since there is only ever one repeated scan). To examine both within- and between-session ICC, we retained participants from the NYU dataset that had three rs-fMRI scans obtained across two scan sessions. Scans 1 and 2 were collected during session 1 and were separated within session by 28 minutes, on average (SD = 9 days). Scan 3 was collected in session 2, which was separated from session 1 by an average of 90 days (SD = 72 days). Intra-session ICC was computed using scan 1 and scan 2 and inter-session ICC using scan 1 and scan 3.

#### 2.4.6 Sensitivity to clinical differences in functional connectivity

Resting-state fMRI is widely used to characterize case-control differences in functional connectivity. Motion can be a major confound in these analyses, particularly as some patient populations may be more prone to move in the scanner, for example due to heightened anxiety in the MRI environment, medication side-effects, or symptoms of the disease itself (e.g., tremor in Parkinson disease). An optimal denoising pipeline will remove spurious differences in functional connectivity that are driven by motion, and therefore isolate any “true” differences between groups. We assessed how different pipelines impact sensitivity to detect clinical differences by conducting case-control comparisons in: (1) the BMH dataset, comparing HCs and OCD patients; and (2) the CNP dataset, comparing HCs and SCZ patients.

To keep our analyses comparable with the wider literature examining pathophysiological mechanisms in mental health, we evaluated sensitivity to clinical group differences using the network-based statistic (NBS; Zalesky et al., 2010). The NBS improves statistical power over typical mass-univariate correction strategies such as FDR by testing the null hypothesis at the level of connected components of edges (rather than individual edges). These components are formed by applying an initial threshold to the data of *p*<0.05 uncorrected. The observed component sizes are compared to an empirical null distribution of maximal component sizes obtained by permuting group membership 5,000 times. Evaluating observed component sizes with respect to a null distribution of maximal sizes ensures that the resulting inferences on network components are corrected for multiple comparisons (Fornito et al., 2016). For each pipeline, an *F*-test was performed using the NBS to identify whether the patients and HCs differed in their functional connectivity in either direction (i.e., a two-tailed hypothesis test). For each significant sub-network of edges identified by the NBS, we first split the edges according to direction by calculating *t*-values for HCs>Patients and Patients>HCs contrasts and retained only the *t*-values for edges that were significant according to the NBS *F*-test. Then, for each set of significant *t*-values, we quantified the size as the proportion of significant edges in the subnetwork relative to the total number of connections in the parcellation. As with HLM contrasts, age (demeaned), sex, and if applicable, scanner site, were entered as nuisance covariates.

Finally, we sought to understand whether group differences identified by distinct pipelines in the same direction implicated similar sets of edges. In other words, we sought to examine whether pipelines showing differences in the same direction identified the same sets of edges as differing between groups. To address this question, we quantified the overlap between the edges comprising significant NBS components identified under different processing regimes. This overlap was quantified separately for differences in the HCs>patients and patients>HCs directions. Overlap was estimated using the Jaccard Index, which is the number of the intersecting edges in a significant component normalized by the union of the component edges across the two pipelines. A value of 0 indicates no overlap between edges and a maximal value of 1 indicates perfect overlap.

### 2.5 Participant exclusion

fMRI data are expensive to acquire and researchers are often reluctant to exclude datasets exhibiting high motion. Some of the suggested exclusion criteria for motion (e.g., Satterthwaite et al., 2013; Power et al., 2014) can lead to drastically reduced sample sizes, particularly in clinical studies where the initial pool of participants may already be small. We therefore examined denoising performance after excluding participants according to three different levels of stringency. First, under a lenient regime designed to emulate a “worst case” scenario, we only excluded participants with high levels of gross motion, defined as >0.55mm mFD (Satterthwaite et al., 2012). Second, under a stringent regime, we excluded participants if any of the following criteria were true: (i) mFD > 0.25mm; (ii) more than 20% of the FDs were above 0.2 mm; and (iii) if any FDs were greater than 5mm (adapted from Satterthwaite et al., 2013). For both lenient and stringent criteria, FD_Jenk_ was used. The third approach involved the exclusion of participants based on criteria contingent on volume censoring methods. With volume censoring, participants are excluded if a certain proportion of volumes are flagged for censoring. In the present analysis, participants were excluded if censoring resulted in less than 4 mins of uncensored data (Satterthwaite et al., 2013 (see *2.3.6 Volume censoring*)). Censoring-based exclusion, which generally resulted in the highest number of participants being excluded, was performed independently of lenient and stringent criteria. Note that this criterion is intrinsic to the volume censoring methods and is not necessary for other pipelines. However, in order to facilitate fair comparisons across methods in the same sample, we also applied censoring exclusion to non-censoring denoising techniques to enable comparison on the same subsample of individuals.

## Results

### 3.1 Sample characteristics

To characterise the level of motion in our datasets, we report mFD averaged across participants at both the whole sample-level as well as the subgroup-level when applying lenient, stringent, and censoring-based exclusionary criteria in Table 3.

**Table 3.** Motion characteristics for the Beijing, CNP, BMH, and NYU datasets. Temporal mean of framewise displacement (mFD) was averaged at the sample-level as well the subgroup-level, where subgroups were defined following lenient, stringent, and censoring-based exclusionary criteria.

| *Exclusion Criteria* | *Beijing*<br>(Mean±SD) | *CNP* | *BMH* | *NYU (scan 1 / 2 / 3)* |
| --- | --- | --- | --- | --- |
| *Lenient* | *N = 192* | *N = 170* | *N = 73* | *N = 29* |
| *Sample-level* | 0.06±0.02 | 0.12±0.08 | 0.07±0.04 | 0.07±0.05/0.07±0.04/0.08±0.05 |
| *HC* | - | 0.11±0.07 | 0.07±0.04 | - |
| *Patient* | - | 0.16±0.09 | 0.06±0.02 | - |
| *Stringent* | *N = 192* | *N = 146* | *N = 71* | *N = 26* |
| *Sample-level* | 0.06±0.02 | 0.10±0.04 | 0.07±0.03 | 0.06±0.02/0.07±0.04/0.06±0.03 |
| *HC* | - | 0.09±0.03 | 0.07±0.03 | - |
| *Patient* | - | 0.12±0.04 | 0.07±0.03 | - |
| *Censoring: SpikeReg* | *N = 192* | *N = 144* | *N = 73* | *N = 28* |
| *Sample-level* | 0.06±0.02 | 0.10±0.04 | 0.07±0.04 | 0.07±0.04/0.07±0.04/0.07±0.03 |
| *HC* | - | 0.09±0.03 | 0.07±0.04 | - |
| *Patient* | - | 0.12±0.04 | 0.08±0.04 | - |
| *Censoring: JP12Scrub* | *N = 192* | *N = 101* | *N = 65* | *N = 25* |
| *Sample-level* | 0.06±0.02 | 0.08±0.02 | 0.07±0.04 | 0.06±0.02/0.07±0.03/0.06±0.03 |
| *HC* | - | 0.08±0.02 | 0.07±0.03 | - |
| *Patient* | - | 0.08±0.02 | 0.08±0.04 | - |
| *Censoring: JP14Scrub* | *N = 186* | *N = 85* | *N = 47* | *N = 22* |
| *Sample-level* | 0.06±0.02 | 0.07±0.02 | 0.06±0.02 | 0.05±0.01/0.06±0.03/0.06±0.02 |
| *HC* | - | 0.07±0.02 | 0.06±0.02 | - |
| *Patient* | - | 0.08±0.02 | 0.06±0.02 | - |

Two-sample *t*-tests run following lenient exclusion revealed that mFD was significantly higher in the CNP dataset relative to the Beijing, BMH, and NYU datasets (Beijing, *t* = 8.89, *p*<0.001; BMH, *t* = 5.24, *p*<0.001; NYU scan 1, *t* = 4.91, *p*<0.001), indicating that participants in the CNP dataset had greater in-scanner movement (mFD did not differ between Beijing, BMH and NYU datasets). For the CNP dataset, average mFD was significantly lower in HC compared to SCZ (lenient: *t*(66) = 3.85, *p*<0.001). For the BMH data, average mFD was virtually identical for HC and OCD groups from the BMH dataset. Finally, within-session ICC was 0.63 and between-session ICC was 0.40 for the NYU dataset.

### 3.2 QC-FC correlations

Our first aim was to examine the efficacy of different noise correction pipelines in removing the relationship between participant in-scanner movement and estimates of functional connectivity, as assessed with QC-FC correlations. We examined QC-FC correlations separately for the Beijing, BMH, CNP and NYU datasets.

Figure 1 shows the percentage of 55,278 edges from the Gordon parcellation with significant QC-FC correlations (*p*<0.05, uncorrected. See supplementary materials for results using *p*<0.05, FDR-corrected) as well as the distributions of QC-FC correlations (the median of the absolute QC-FC distribution is shown alongside the distributions), for the Beijing and CNP datasets following application of each of the 19 denoising pipelines listed in Table 2. Beijing and CNP are presented here as examples of well-powered low-motion (i.e., Beijing) and high-motion (i.e., CNP) datasets (see supplementary material for results for the BMH and NYU datasets). The Beijing dataset is plotted only once in Figure 1 because no participants were excluded under both lenient and stringent criteria; results were thus identical.

**Figure 1.**
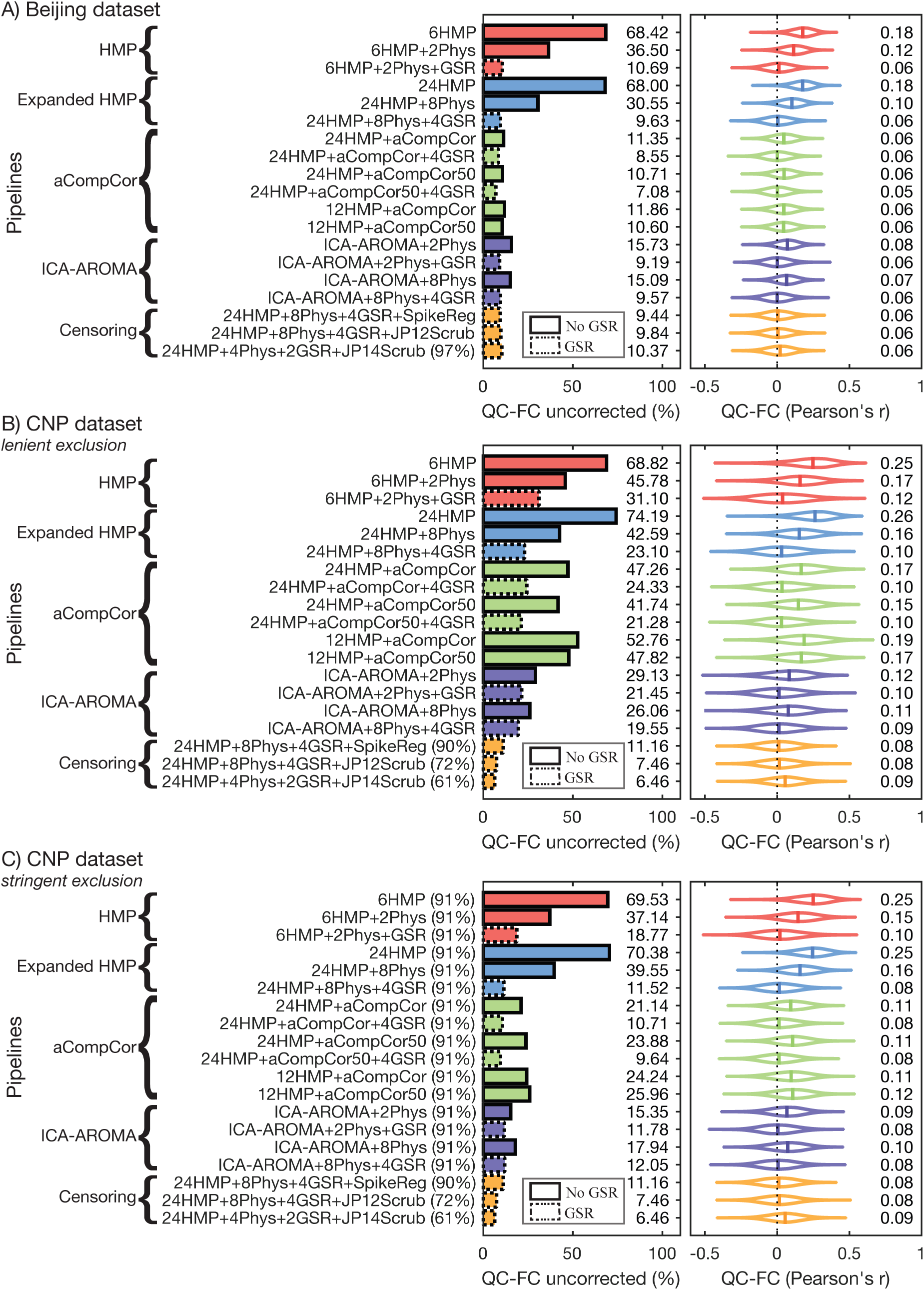
The residual effect of in-scanner motion on functional connectivity after noise correction with one of nineteen different rfMRI pre-processing pipelines. Functional connectivity at each edge was correlated with a summary metric of in-scanner movement across the entire sample (QC-FC correlations) for the Beijing and CNP datasets. The proportion of functional correlations that showed significant (*p*<0.05, uncorrected) associations with subject motion are shown in the left panels. The full distributions of QC-FC values are shown for each pipeline in the right panels with the corresponding median absolute values printed alongside. Results for the Beijing (*A*) as well as the CNP dataset under lenient (*B*) and stringent (*C*) exclusion criteria are presented here. No participants were excluded from the Beijing dataset under lenient or stringent exclusion criteria due to low motion in this sample. See supplementary material for results for the BMH and NYU datasets. Percentages in parentheses next to pipeline labels are the proportion of HCs retained in analysis. Absent parentheses indicate that 100% of HCs were retained. Dotted lines around the horizontal bars denote pipelines that incorporated global signal regression (GSR).

No pipelines successfully reduced QC-FC significant correlations to 0% or the median absolute QC-FC correlation to 0 in either the low-motion Beijing dataset, or the high-motion CNP dataset, regardless of which exclusion criteria were applied. Simple strategies relying on motion parameters alone (i.e., 6HMP and 24HMP) fared the worst, with significant QC-FC correlations as high as 68% (median absolute correlation of 0.18) in the Beijing dataset and 74% (median absolute correlation of 0.26) in the CNP dataset. This result suggests that most measures of functional connectivity in analyses using HMP correction alone are contaminated by motion, as similarly reported by Ciric et al. (2017). Adding tissue-averaged physiological regressors (e.g., 2Phys or 8Phys) substantially improved the performance of the HMP models, with the further addition of GSR having a very large impact (e.g., a reduction of 68% to 11% significant QC-FC correlations in the Beijing data and a ~66% reduction in median absolute correlation).

Relative to the HMP+Phys models, aCompCor showed improved performance for the low-motion Beijing dataset but not for the high-motion CNP dataset. For example, significant QC-FC correlations changed from ~30% to ~11%, and median absolute correlations dropped by ~40%, when switching from 24HMP+8Phys to 24HMP+aCompCor for the Beijing dataset, whereas QC-FC correlations remained comparable for the CNP dataset. As above, adding GSR improved the performance of aCompCor in both datasets. Using more principal components (i.e., switching from aCompCor to aCompCor50) only marginally improved performance in both datasets. These findings suggest that aCompCor is ineffective for denoising high motion datasets.

Pipelines incorporating ICA-AROMA outperformed the HMP+Phys and aCompCor pipelines for the high-motion CNP dataset. Adding GSR to the ICA-AROMA pipelines led to a slight improvement in performance for the Beijing and CNP datasets.

Censoring pipelines consistently performed very well in both datasets, yielding ~10% of significant QC-FC correlations or less and median absolute correlations ranging between 0.06 and 0.09. The best performing pipeline for the high-motion CNP dataset was 24HMP+4Phys+2GSR+JP14Scrub (i.e., optimized scrubbing), which yielded only 6.5% significant QC-FC correlations and a median absolute correlation of 0.09. However, the censoring-based pipelines resulted in the exclusion of 10-29% of HC participants (see Figure 1B).

In general, the application of stringent exclusion criteria improved the performance of all pipelines (Figure 1C). The largest improvements were seen for aCompCor and aCompCor50 pipelines (without GSR), suggesting that anatomical PCA-based correction methods are less effective at denoising high-motion participants than other methods.

### 3.4 QC-FC distance-dependence

An ideal denoising procedure should leave no residual association between QC-FC and inter-regional distance. Figure 2 shows the Spearman rank correlation coefficient between QC-FC correlations and edge-wise Euclidean distance for each pipeline. As above, the Beijing and CNP datasets are presented here and results for the BMH and NYU datasets are presented in the supplementary materials. Figure 3 visualises this relationship for a selection of pipelines across a series of equiprobable distance bins for the high-motion CNP dataset. Visualisations for the low-motion Beijing, BMH and NYU datasets appear in Figures S3, S6 and S9.

**Figure 2.**
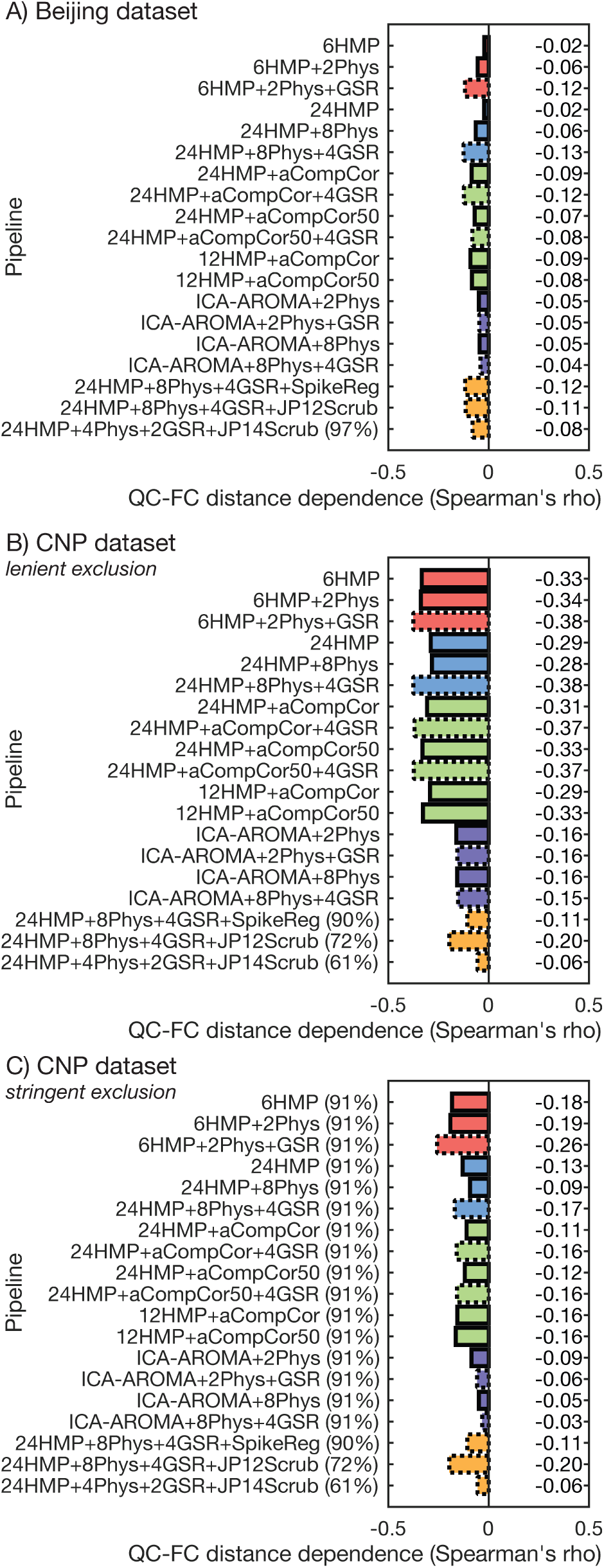
The distance-dependence of QC-FC correlations following application of each pipeline to the Beijing and CNP datasets. Distance-dependence was quantified here as a Spearman rank correlation between the QC-FC correlation of a given edge and the Euclidean distance separating the two coupled regions. Results for the Beijing (*A*) as well as the CNP dataset under lenient (*B*) and stringent (C) exclusion criteria are presented here. No participants were excluded from the Beijing dataset under lenient or stringent exclusion criteria. See supplementary material for results for the BMH and NYU datasets. Percentages in parentheses next to pipeline labels represent the proportion of HCs retained in analysis (absent parentheses indicate that 100% of HCs were retained).

**Figure 3.**
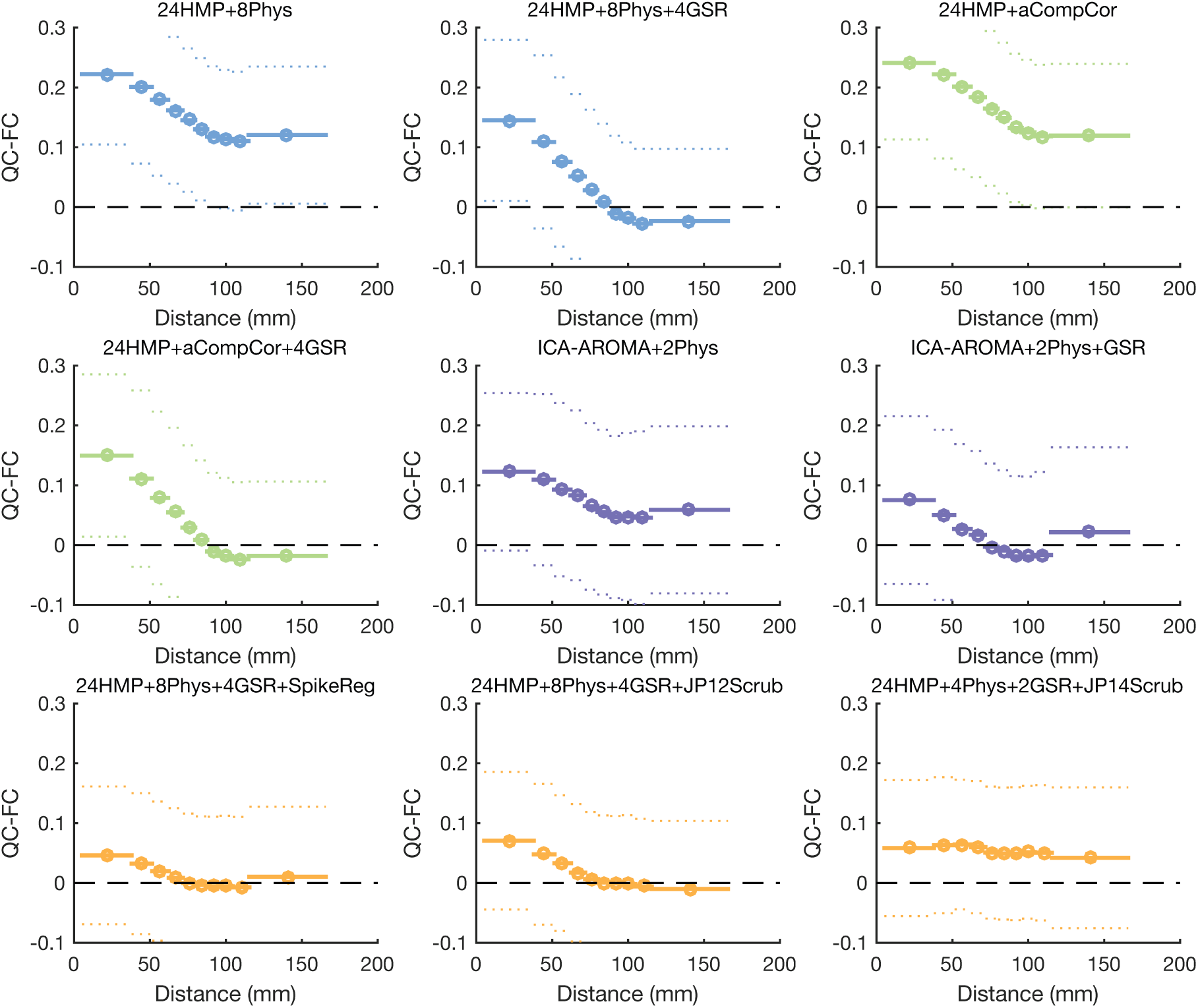
QC-FC distance-dependence for a selection of pipelines applied to the CNP dataset under lenient exclusion criteria. QC-FC correlations were split into 10 equiprobable bins (using equally spaced quantiles to define bins) based on the distance between nodes. For each bin, the mean distance between edges (circle) as well as the mean (solid line) and standard deviation (dotted line) of QC-FC correlation are shown.

Figure 2A shows that for the low-motion Beijing dataset, the HMP pipelines without any Phys or aCompCor correction showed the least linear distance-dependence, followed by the ICA-AROMA pipelines. Figure 2B shows that for the high-motion CNP dataset, the ICA-AROMA and censoring pipelines generally showed the weakest linear distance-dependence. Figure 3 shows that ICA-AROMA shows a moderate non-monotonic distance-dependence that is not adequately captured by the Spearman rank correlation in Figure 2.

For methods that were not based on ICA-AROMA, GSR generally increased distance-dependence in both Beijing and CNP. This effect occurred despite GSR having a generally positive effect on QC-FC correlations (Figure 1). This result fits with previous reports demonstrating that GSR improves QC-FC correlations at the cost of increased distance-dependence (Ciric et al., 2017).

Figure 2C shows that applying stringent exclusionary criteria to non-censoring pipelines improved linear QC-FC distance-dependence and resulted in most ICA-AROMA pipelines showing lower distance-dependence than the censoring pipelines, despite the censoring pipelines often excluding more people (see Table 3). Figure 3 shows that stringent exclusion reduced the non-monotonic distance-dependence of ICA-AROMA to a level comparable to the basic scrubbing pipeline, but optimized scrubbing produced a flatter distance-dependence profile (see Figure S4).

### 3.5 Volume censoring

Volume censoring generally performed well on the QC-FC benchmarks reported above, but these censoring pipelines generally required the exclusion of more participants than our lenient and stringent criteria due to the requirement to at least 4 minutes of uncensored data be retained (see Methods). In this sense, the comparisons above are not “fair” because the censoring methods were evaluated on a subset of participants characterized by less motion (see Table 3). To enable a fair comparison in the same set of people, we replicated the QC-FC correlations and QC-FC distance-dependence in the CNP dataset after applying the censoring-based exclusion derived from optimized scrubbing to all pipelines (Figure 4). The exclusion criteria for optimized scrubbing was chosen because it was the most stringent. This resulted in the exclusion 47 of the 121 HC participants from the CNP dataset (i.e., 61% of HC participants were retained).

**Figure 4.**
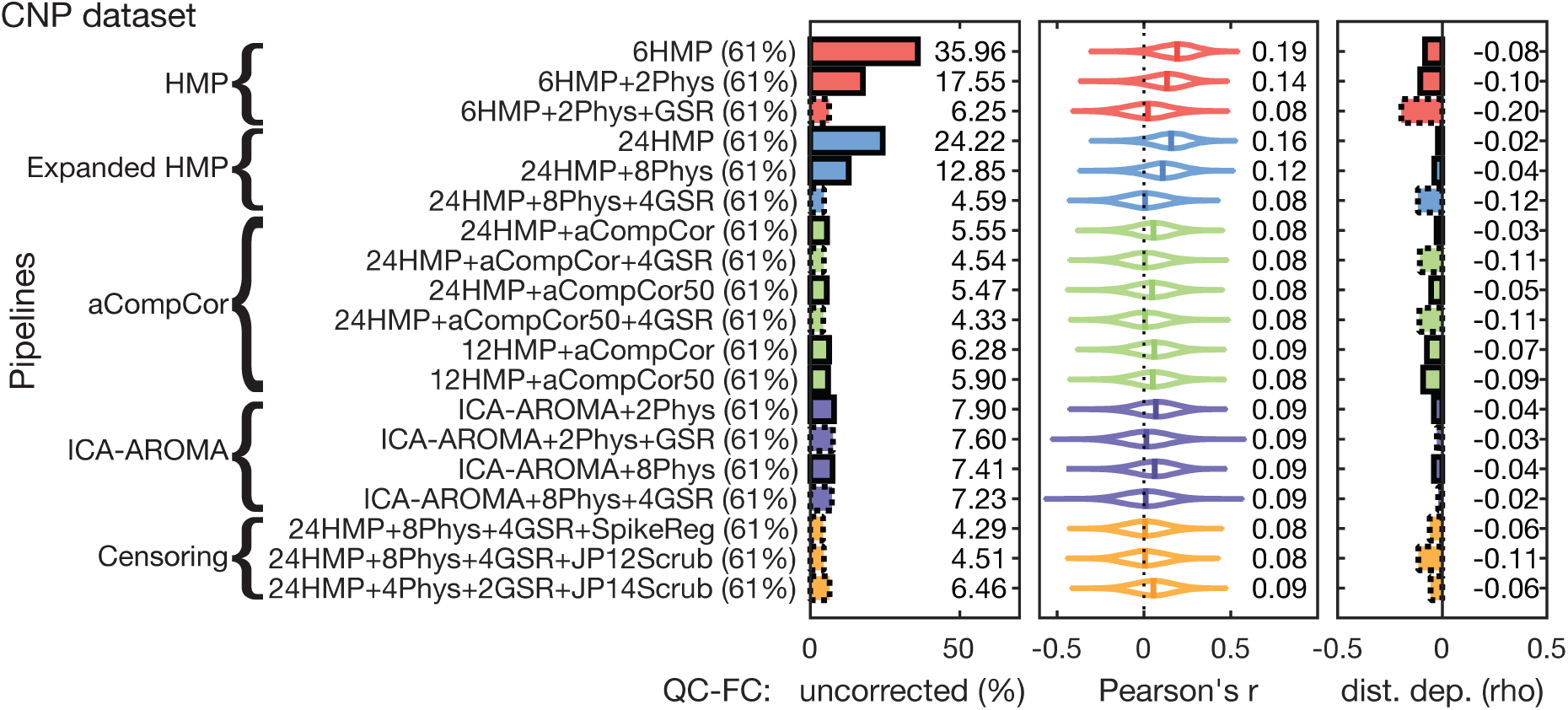
Effect of excluding participants with <4 minutes of uncensored data on mitigating motion-related artefact in the CNP dataset. We plot the proportion of edges exhibiting significant QC-FC correlations (left), the QC-FC distributions (middle), and QC-FC distance-dependence as Spearman correlation coefficients (right) all calculated following exclusion of participants according to optimized scrubbing criteria (see Methods). Percentages in parentheses next to pipeline labels represent the proportion of HCs retained in analysis.

Figure 4 shows that exclusion of HC participants with high numbers of contaminated volumes can dramatically improve QC-FC correlations and QC-FC distance-dependence across all pipelines, with the exception of the basic and expanded HMP+Phys pipelines. For the other pipelines, QC-FC correlations dropped to <10%, with some pipelines yielding <5%. QC-FC median absolute values also generally dropped for all pipelines, with all pipelines except the basic and expanded HMP models without GSR yielding <0.10. QC-FC distance-dependence was reduced as well, often by 30-50%. Figure S5 shows that when the optimized scrubbing exclusion criteria are applied to all pipelines, ICA-AROMA+2Phys and optimized scrubbing pipeline yield similarly flat profiles of distance-dependence, although applying GSR to ICA-AROMA appears to exacerbate the non-monotonic effect. These results suggest that much of the benefit of volume censoring comes from the exclusion of high-motion individuals.

### 3.6 Loss of temporal degrees of freedom

Using an increasing number of nuisance regressors reduces the degrees of freedom in the BOLD data, which can lead to spurious increases in functional connectivity (Yan et al., 2013a). Thus, we next characterised the loss of temporal degrees of freedom (tDOF-loss) for each pipeline and dataset. Results for Beijing and CNP are presented in Figure 5 (see supplementary material for BMH and NYU datasets) after applying lenient exclusion criteria. For all datasets, the aCompCor50 pipelines resulted in the highest tDOF-loss, with the loss often approximately double the tDOF-loss of the other pipelines. As expected, the relatively simple 6HMP class of models yielded the lowest tDOF-loss. The ICA-AROMA pipelines often showed lower tDOF-loss relative to all other pipelines (except for the non-expanded HMP pipelines), and were associated with lower tDOF-loss in the high-motion CNP data compared to the low-motion Beijing data. This result suggests that ICA-AROMA may be more effective at identifying noise components in high-motion compared to low-motion datasets.

**Figure 5.**
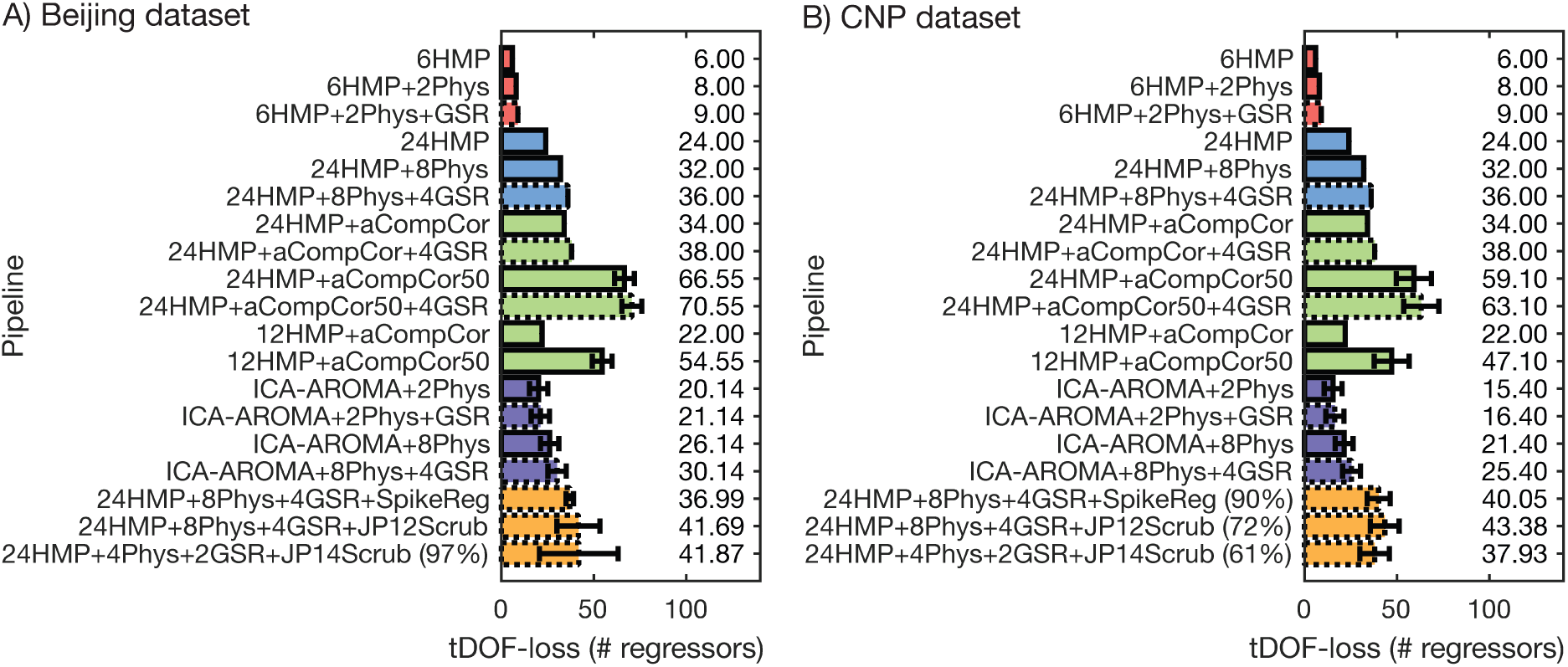
Different fMRI noise correction methods reduce tDOF by different amounts in the (A) Beijing and (B) CNP data. The ICA-AROMA and aCompCor50 methods define a variable number of noise regressors in different participants. For these pipelines, the sample mean and standard deviation (error bars) are presented. Pipelines using aCompCor50 yield the highest tDOF-loss. ICA-AROMA pipelines offer relatively modest tDOF-loss despite being highly effective at mitigating motion artefacts. Percentages in parentheses next to pipeline labels represent the proportion of HCs retained in analysis (absent parentheses indicate that 100% of HCs were retained).

With regards to aCompCor50, tDOF-loss is not split evenly across the WM and CSF compartments. For the Beijing dataset, the WM-specific tDOF-loss was 40±5 and CSF-specific tDOF-loss was 2±1. For the BMH dataset, the WM-specific tDOF-loss was 43±6 and CSF-specific tDOF-loss was 4±2. For the CNP dataset, the WM-specific tDOF-loss was 32±9 and CSF-specific tDOF-loss was 3±2. For the NYU dataset, the WM-specific tDOF-loss was 38±9 and CSF-specific tDOF-loss was 5±3.

For many of our pipelines, the number of nuisance regressors, and hence the number of degrees of freedom, is explicitly set in the model and is invariant across participants. However, for ICA-AROMA, aCompCor50, and censoring-based pipelines the temporal degrees of freedom are free to vary across participants. When comparing two or more groups, this variability in tDOF-loss may confound comparisons of functional connectivity (Yan et al., 2013b). To investigate this possibility in our data, we performed two-sample *t*-tests comparing tDOF-loss between HCs and patients in the BMH and CNP datasets for the 24HMP+aCompCor50, ICA-AROMA+2Phys, and 24HMP+4Phys+2GSR+JP14Scrub pipelines. For the BMH dataset, no significant difference was observed in tDOF-loss between HCs and OCD participants for any of the pipelines. For the CNP dataset, tDOF-loss was significantly higher in the SCZ participants (mean tDOF-loss = 19±6) compared to HCs (mean tDOF-loss = 15±5) for the ICA-AROMA+2Phys pipeline (*t(74)* = 4.18, *p* < 0.001, uncorrected). No difference was observed for the 24HMP+aCompCor50 or the 24HMP+4Phys+2GSR+JP14Scrub pipeline.

These results suggest that ICA-AROMA is effective at reducing QC-FC and QC-FC distance-dependence for only moderate tDOF-loss. However, if there are systematic group differences in the amount of head motion, tDOF-loss may also vary across groups when using this approach.

### 3.7 Group differences in high-motion and low-motion healthy participants (HLM contrasts)

Assuming similar demographic characteristics, two groups of HC participants should exhibit similar functional connectivity patterns. As such, any differences observed between groups of HCs split by motion can be attributed to ineffective denoising. We thus examined differences in functional connectivity between low-motion and high-motion subsets of HC participants from the Beijing and CNP datasets. The BMH and NYU datasets were not examined due to insufficient sample sizes. We performed HLM contrasts after applying both lenient and stringent exclusion criteria to all non-censoring pipelines (as above, censoring pipelines had their own exclusion applied).

Figure 6 shows the sizes of the significant whole-brain effects (represented as a percentage of connections comprising the significant component), identified in a contrast of high-motion HCs and low-motion HCs for each pipeline and dataset. Predictably, in both datasets, the largest effects are for high-motion HCs showing increased functional connectivity relative to low-motion HCs.

**Figure 6.**
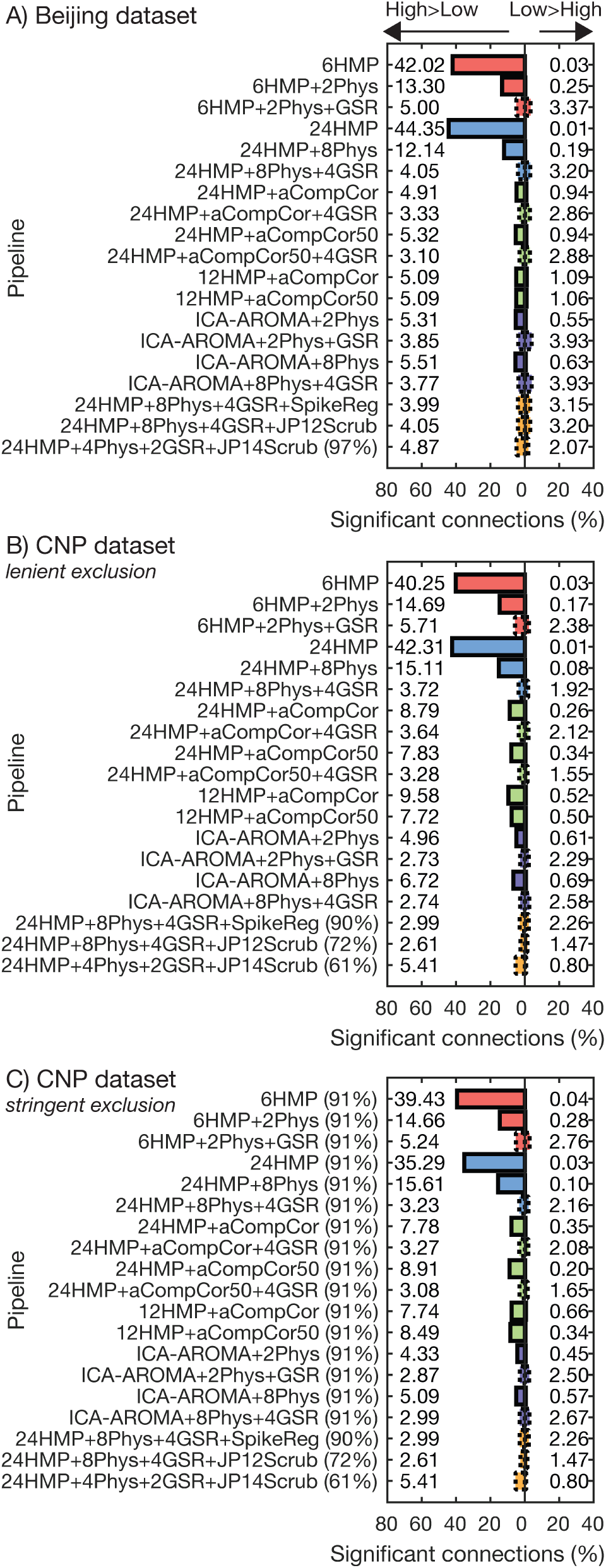
Differences in functional connectivity between groups of healthy controls (HC) that differ on their amount of in-scanner movement. Differences in functional connectivity are plotted as the percentage of edges in which functional connectivity was significantly higher for high-motion compared to low-motion HCs at *p*<0.05, uncorrected, for the Beijing (*A*) as well as the CNP dataset under lenient (*B*) and stringent (C) exclusion criteria. Contrasts are two-tailed. Percentages in parentheses next to pipeline labels represent the proportion of HCs retained in analysis (absent parentheses indicate that 100% of HCs were retained).

In both datasets, the relative performance of the different pipelines was similar to the analysis of QC-FC correlations (Figure 1); that is, pipelines that showed high levels of QC-FC correlations also yielded more connectivity differences between high- and low-motion HC subgroups. As expected, applying stringent exclusionary criteria to the CNP dataset resulted in fewer differences for all the non-censoring pipelines.

### 3.8 Test-retest reliability

Next, we examined test-retest reliability across our pipelines using the ICC calculated using the NYU dataset. High ICC is obtained when estimates of functional connectivity at each edge are consistent across repeated scans. Means and standard deviations of ICC are plotted for each pipeline separately for within- and between-session TRT in Figure 7, which shows that the HMP models typically yielded the highest short-term (within-session) and long-term (between-session) reproducibility. This is unsurprising considering that physiological and motion-related noise in the BOLD signal may be a trait-like component that is reproducible across sessions (Couvy-Duchesne et al., 2014), and that these methods are ineffective at removing such noise. Beyond the HMP models, ICC was fairly consistent across pipelines. Results are shown for lenient exclusion but we note that the results under stringent exclusion are very similar due to only 3 participants being excluded. Under censoring-based exclusion, the individuals excluded varied slightly across scan sessions, so participants were excluded from the ICC analysis if they were marked for exclusion in any of the three scan sessions. This procedure resulted in excluding 3, 6, and 14 participants from the spike regression, basic scrubbing, and optimized scrubbing, respectively.

**Figure 7.**
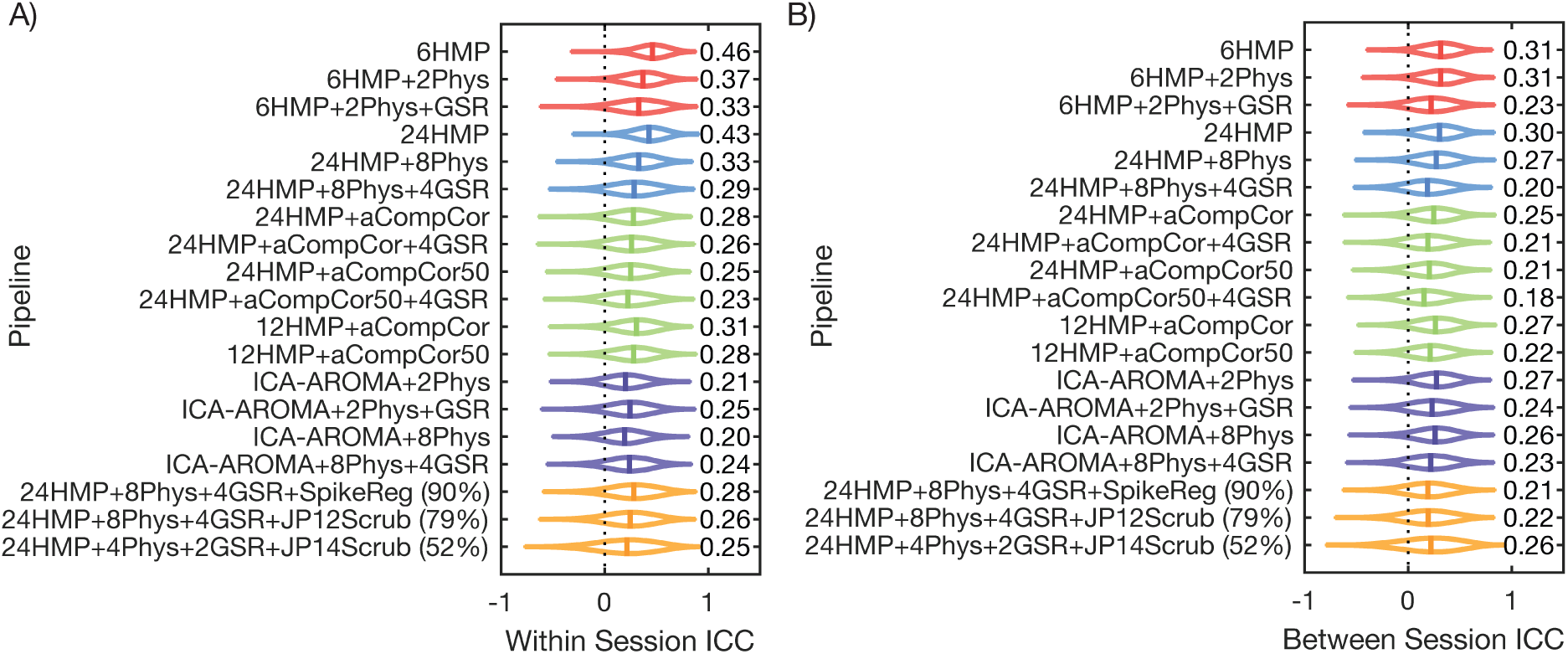
Test-retest reliability varies over fMRI noise correction pipelines. (*A*) Intra-class correlation (ICC) coefficients for within session test-retest from the NYU dataset. (*B*) ICC coefficients for between session test-retest from the NYU dataset. Percentages in parentheses next to pipeline labels represent the proportion of HCs retained in analysis (absent parentheses indicate that 100% of HCs were retained).

### 3.9 Interim summary

Taken together, the above results allow us to draw six key conclusions. First, HMP+Phys models without GSR are ineffective at mitigating motion-related artefact from rs-fMRI, regardless of the level of motion in the sample, exclusionary criteria applied, or the use of expansion terms (see Figures 1, 2, 4, and 6). The application of GSR dramatically improves the performance of these pipelines. Second, aCompCor pipelines may only be viable in low-motion datasets and perform poorly in high motion data (see Figure 1). Third, ICA-AROMA and censoring pipelines are superior to the other denoising strategies, yielding the lowest QC-FC correlations (see Figure 1 and 4), lowest QC-FC distance-dependence (see Figure 2 and 4), and minimal functional connectivity differences between high- and low-motion healthy controls (see Figure 6). Fourth, a major part of the benefit to motion-related artefact control in censoring pipelines comes from the exclusion of participants with <4 minutes of uncensored data; when this criterion is applied to all pipelines, performance differences are marginal (except for the HMP pipelines without GSR). Fifth, aCompCor and censoring pipelines yield high tDOF-loss, marking them as relatively expensive methods for controlling for motion-related artefact in rs-fMRI data. Finally, methods that were more effective in denoising were associated reduced test-retest reliability, suggesting that noise signals in BOLD data are reproducible.

### 3.10 Group differences in functional connectivity

We next examined the sensitivity of each pipeline to clinical group differences in functional connectivity in the BMH and CNP datasets using the NBS. No significant group differences were found in the comparison of OCD patients and HCs for lenient or stringent criteria. In the comparison of HC and SCZ, several different pipelines showed group differences. For each pipeline and contrast, only one significant NBS component was found. Each significant component is represented as a percentage of all possible edges in Figure 8. NBS contrasts were performed after applying both our lenient and stringent exclusion criteria to all non-censoring pipelines, with censoring pipelines having their own exclusion criteria applied.

**Figure 8.**
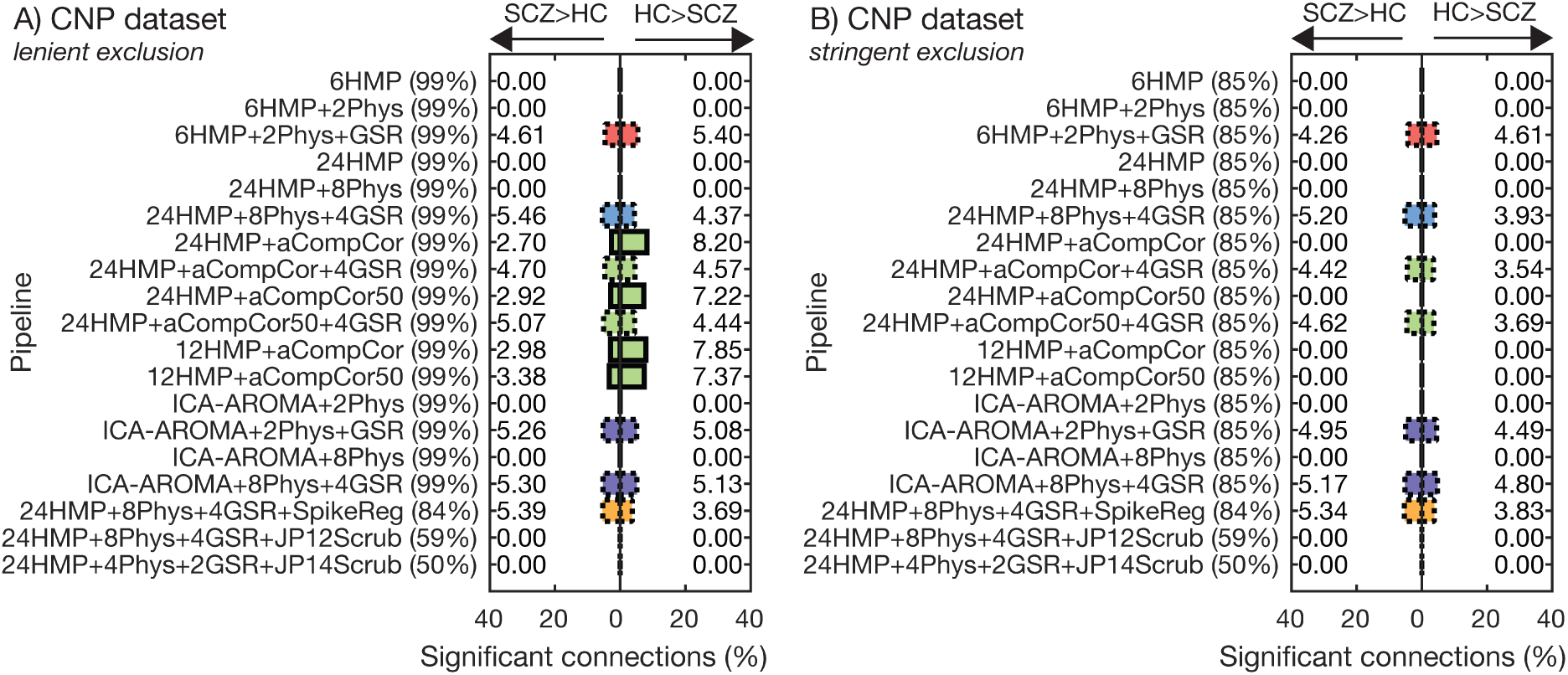
Case-control differences depend on denoising strategy for the CNP dataset. The proportion of edges contained in the NBS subnetwork showing significant differences between healthy controls and individuals with schizophrenia, *p*<0.05, component-wide corrected. Percentages in parentheses next to pipeline labels represent the proportion of participants retained in analysis.

Two general results are apparent from Figure 8. First, when stringent exclusion criteria are applied, differences are only observed for pipelines that included some form of GSR, each of which all showed bidirectional effects in the order of ~4-5% of edges in each direction. Second, the only pipelines without GSR to show any effects were the aCompCor/aCompCor50 pipelines under lenient exclusion criteria. Notably, these pipelines also performed poorly on our QC-FC benchmarks in the CNP dataset, suggesting that these effects may be driven by motion-related artefact.

Next, we used the Jaccard Index to quantify the consistency of the edges identified in each analysis as showing group differences under strict exclusion criteria. As shown in Figure 9, the most consistent effect was found with ICA-AROMA, where the overlap between ICA-AROMA+8Phys+4GSR and ICA-AROMA+2Phys+GSR was 0.7. Overlap between the other pipelines was generally quite low, ranging between 0.2 and 0.4; the only other pair to exceed 0.5 was the overlap between 24HMP+8Phys+4GSR and 24HMP+8Phys+4GSR+SpikeReg.

**Figure 9.**
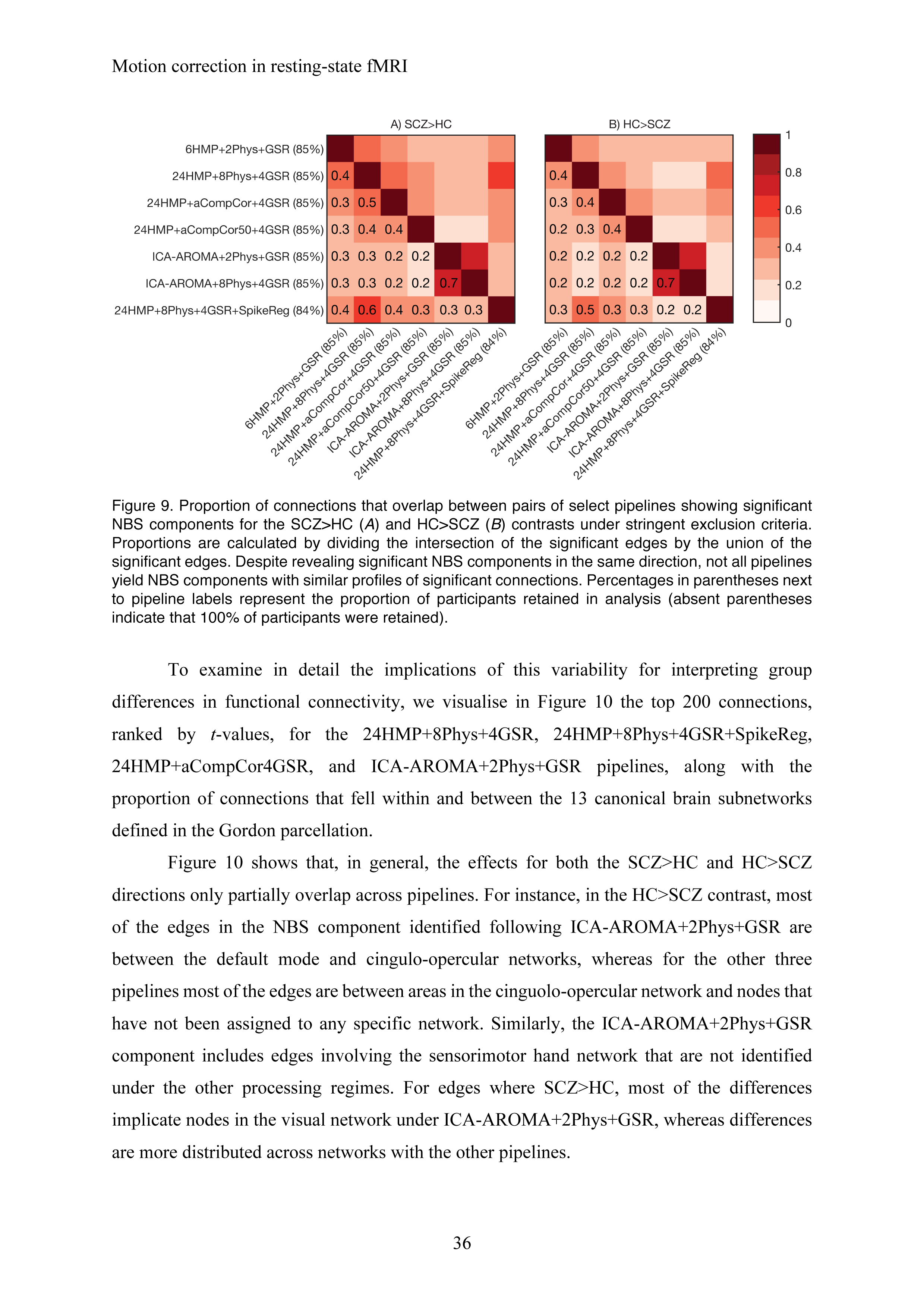
Proportion of connections that overlap between pairs of select pipelines showing significant NBS components for the SCZ>HC (*A*) and HC>SCZ (*B*) contrasts under stringent exclusion criteria. Proportions are calculated by dividing the intersection of the significant edges by the union of the significant edges. Despite revealing significant NBS components in the same direction, not all pipelines yield NBS components with similar profiles of significant connections. Percentages in parentheses next to pipeline labels represent the proportion of participants retained in analysis (absent parentheses indicate that 100% of participants were retained).

To examine in detail the implications of this variability for interpreting group differences in functional connectivity, we visualise in Figure 10 the top 200 connections, ranked by *t*-values, for the 24HMP+8Phys+4GSR, 24HMP+8Phys+4GSR+SpikeReg, 24HMP+aCompCor4GSR, and ICA-AROMA+2Phys+GSR pipelines, along with the proportion of connections that fell within and between the 13 canonical brain subnetworks defined in the Gordon parcellation.

**Figure 10.**
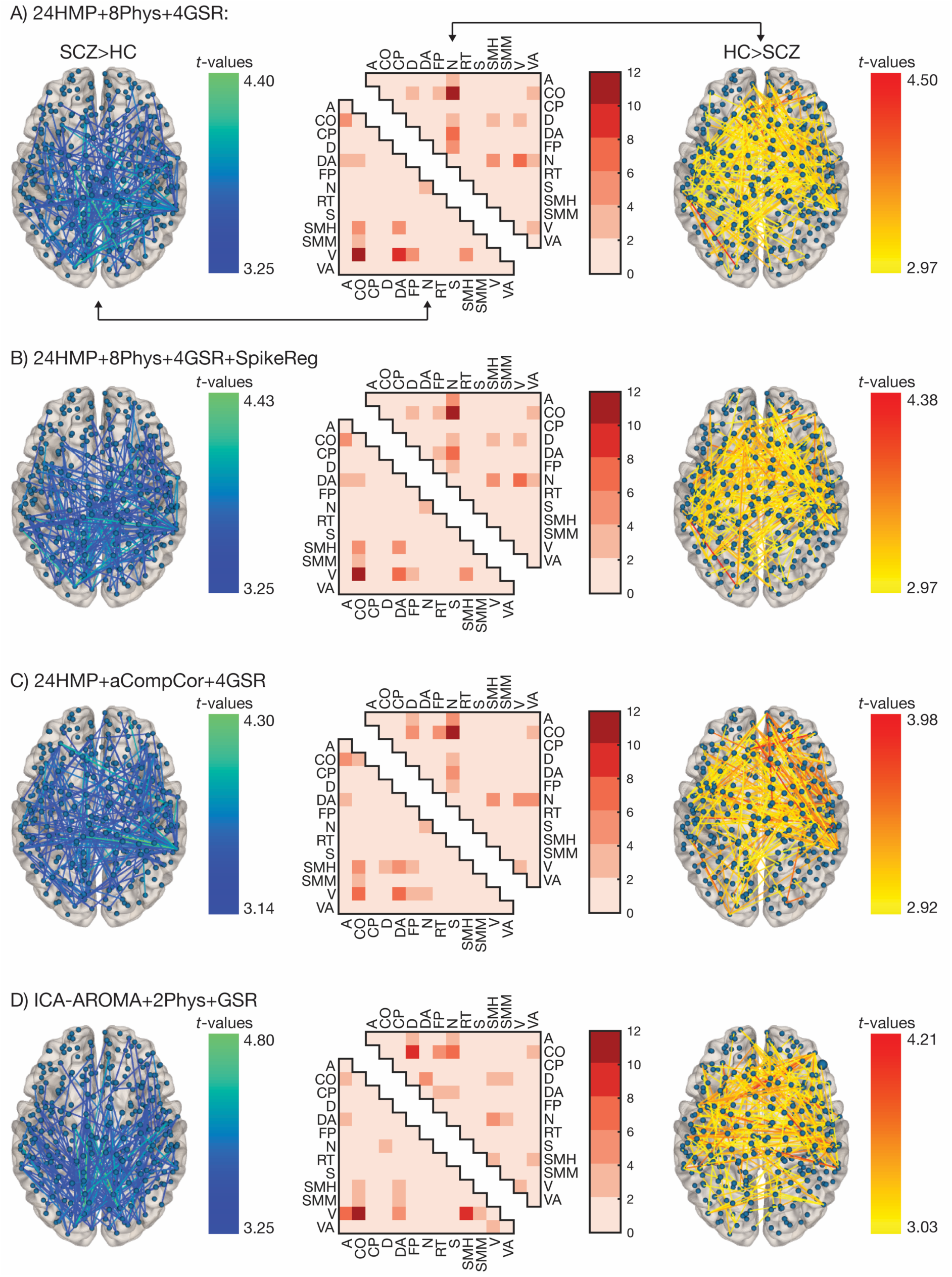
Denoising pipelines impact the anatomical distribution of group differences in functional connectivity. Here we summarize functional connectivity differences between patients with schizophrenia and healthy control participants in the CNP dataset as characterised using the (*A*) 24HMP+8Phys+4GSR, (*B*) 24HMP+8Phys+4GSR+SpikeReg, (*C*) 24HMP+aCompCor+4GSR, and (*B*) ICA-AROMA+2Phys+GSR pipelines. In each subplot, the top 200 edges, ranked by *t*-values, are represented in brain space for the SCZ>HC contrast (left) and the HC>SCZ contrast (right). The percentage of connections that fall within and between brain networks identified by the Gordon parcellation are presented in matrix form in the middle of each subplot. Matrix plots are split into lower (SCZ>HC) and upper (HC>SCZ) triangles (see arrows in subplot A). Anatomical renderings were generated using NeuroMarvl (http://immersive.erc.monash.edu.au/neuromarvl/). Subnetworks in the matrix plot are denoted as follows: A, auditory; CO, cingulo-opercular; CP, cingulo-parietal; D, default mode; DA, dorsal attention; FP, fronto-parietal; N, none; RT, retrosplenial-temporal; S, salience; SMH, sensorimotor hand; SMM, sensorimotor mouth; V, visual; VA, ventral attention.

Figure 10 shows that, in general, the effects for both the SCZ>HC and HC>SCZ directions only partially overlap across pipelines. For instance, in the HC>SCZ contrast, most of the edges in the NBS component identified following ICA-AROMA+2Phys+GSR are between the default mode and cingulo-opercular networks, whereas for the other three pipelines most of the edges are between areas in the cinguolo-opercular network and nodes that have not been assigned to any specific network. Similarly, the ICA-AROMA+2Phys+GSR component includes edges involving the sensorimotor hand network that are not identified under the other processing regimes. For edges where SCZ>HC, most of the differences implicate nodes in the visual network under ICA-AROMA+2Phys+GSR, whereas differences are more distributed across networks with the other pipelines.

Together, these results indicate that the choice of noise correction strategy can have a large effect on case-control comparisons. In the high-motion CNP dataset, the specific sets of edges identified as differing between groups showed considerable variation, leading to different conclusions about which neural systems are affected by disease.

## 4. Discussion

In this study, we compared the performance of 19 different popular denoising strategies for rs-fMRI data using five different benchmarks and four datasets, and then used those benchmarks to inform interpretation of clinical group differences in functional connectivity. Our key results can be summarized as follows:

- Across most benchmarks, particularly QC-FC correlations, QC-FC distance-dependence, and HLM contrasts, the HMP class of pipelines performed poorly in low-motion and high-motion data, indicating that simple regression of head motion parameters, even with expansion terms included, does not sufficiently correct the influence of motion on functional connectivity. Adding mean tissue physiological regressors to the HMP models improved denoising efficacy, but performance was still worse than other classes of pipelines. GSR improved the performance of HMP pipelines on QC-FC correlations but exacerbated QC-FC distance-dependence.
- aCompCor and aCompCor50 pipelines showed similar performance on QC-FC correlations and QC-FC distance-dependence. They also showed among the highest tDOF-loss of all pipelines. Their performance was worse for high-motion than low-motion datasets. GSR generally improved their efficacy.
- Censoring generally performed well across all benchmarks, but a major component of this benefit was derived from the requirement that participants with <4 minutes of uncensored data are excluded. Once this criterion was applied to all pipelines, performance differences between pipelines were reduced (except for the HMP pipelines, which still performed poorly). Censoring pipelines were also expensive in terms of tDOF-loss.
- ICA-AROMA pipelines generally performed well across nearly all benchmarks, often being marginally inferior to censoring pipelines. They also resulted in low-to-moderate tDOF-loss.
- In general, denoising pipelines incurring higher tDOF-loss were associated with lower TRT of functional connectivity estimates, such that the simplest 6HMP model showed the highest within- and between-session TRT and aCompCor50 pipelines showed the lowest. Given the relatively poor performance of the 6HMP model in denoising, it is possible that a substantial portion of reproducible BOLD signal is driven by head motion and other physiological confounds.
- Including GSR generally improved QC-FC correlations but exacerbated QC-FC distance-dependence, except when applied to ICA-AROMA. Group differences between HC and SCZ only emerged following application of GSR and in each case differences were bidirectional (Figure 6).
- In the CNP dataset, the specific set of connections identified as differing between patients and HCs was contingent on the pipeline used. ICA-AROMA pipelines showed the highest consistency of group differences under different pipeline variations but their results differed when compared to non-ICA-AROMA pipelines.

Taken together, these findings indicate that no pipeline can successfully remove all confounding effects of head motion. Censoring and ICA-AROMA are relatively successful, as assessed by nearly all our benchmarks, with the former generally outperforming the latter. However, censoring comes at increased cost both in terms of tDOF-loss and requisite participant exclusion. In the following sections, we discuss some of the key aspects of our results in more detail.

### 4.1 HMP pipelines

The most popular strategy used to correct for the confounding effects of head motion on BOLD signal variance involves simple linear regression using the canonical 6HMP, sometimes with mean tissue regressors and expansion terms included. Our evaluation indicates that this strategy is not effective even in low-motion datasets, unless GSR is also applied. These findings are consistent with past studies showing that the regression of head motion parameters is insufficient for removing the effect of motion on functional connectivity (Ciric et al., 2017; Yan et al., 2013a). Indeed, compared to aCompCor and ICA-AROMA pipelines, Ciric et al. (2017) also found that standard and expanded head motion parameter regression showed the highest QC-FC correlations. Yan et al. (2013a) also found that motion-BOLD correlations were still present across a wide range of expanded head motion models. Here, we found that HMP approaches were the most sensitive to differences between high- and low-motion CNP controls, with the 6HMP model identifying significant differences in functional connectivity across around half of all edges in the network. Thus, analyses that rely on HMP models alone are likely to be heavily contaminated by motion.

### 4.2 aCompCor/aCompCor50 pipelines

Denoising pipelines incorporating variants of aCompCor are becoming increasingly popular, and have previously been shown to be more effective than mean WM/CSF regression for removing the relationship between motion and DVARS (Muschelli et al., 2014). Our evaluation indicates that these methods only outperform HMP+Phys models in low-motion data, and that they are relatively ineffective in high-motion data. This dependence on levels of motion in the data suggests that aCompCor/aCompCor50 pipelines are not sufficiently robust to be generally applicable to a diverse range of data. Furthermore, the aCompCor50 pipelines were also associated with very high levels of tDOF-loss, incorporating ~50-70 noise regressors, on average, across all datasets.

### 4.3 ICA-AROMA pipelines

ICA-AROMA performed consistently well across all datasets. Although its performance decreased in high-motion data (e.g., CNP dataset under lenient exclusion criteria), it was often more effective HMP and aCompCor/aCompCor50 pipelines. Its performance was also consistent regardless of whether ICA-AROMA was combined with 2Phys or 8Phys, and improved only slightly with GSR/4GSR. This consistency is a desirable property of a robust noise correction approach. Consistent with Ciric et al. (2017), we found that ICA-AROMA was associated with a moderate, non-monotonic distance-dependence of QC-FC, whereas censoring pipelines typically yielded lower median absolute QC-FC correlations and flatter distance-dependence plots. When these methods were applied to the exactly same sample, ICA-AROMA showed a similar distance-dependence and median QC-FC to optimized scrubbing. A particular advantage of ICA-AROMA is that it showed the highest consistency of group differences under different pipeline variations (i.e., ICA-AROMA+8Phys+4GSR and ICA-AROMA+2Phys+GSR; Figure 9), which is another desirable property of a denoising algorithm. Note that one caveat to consider when comparing across pipelines is that ICA-AROMA pipelines involved performing spatial smoothing before denoising, as per recommendations from Pruim et al. (2015b).

One caveat of ICA-AROMA is that it results in variable tDOF-loss across participants. As a result, tDOF-loss differed significantly between the SCZ and HC groups in the CNP dataset. One option to address this variation could be to covary for tDOF-loss at the second-level. However, it can be difficult to properly control for the effects of a confound that is not balanced between groups using the general linear model (Miller and Chapman, 2001).

### 4.4 Volume censoring

Volume censoring is another popular method for removing motion-related confounds from BOLD data. Explicitly removing high motion time points has been shown to be very effective at motion correction (Power et al., 2014; 2013; 2012). However, the approach can distort the temporal properties of the data, precluding analysis of non-stationary dynamics (Hutchison et al., 2013; Zalesky et al., 2014) and other time series properties (Sethi et al., 2017). Denoising methods that do not use censoring attempt to recover true signal fluctuations, but it remains an open question whether such fluctuations can indeed be recovered in high-motion datasets.

In our analysis, the primary advantage of volume censoring came from the exclusion of participants with high FDs. As shown in Figure 4, applying stringent censoring-based exclusion of high-motion participants generally led to a dramatic reduction in QC-FC correlations, regardless of whether a HMP, aCompCor, or ICA-AROMA-based pipeline was used for denoising. After stringent exclusion, censoring-based pipelines produced a small reduction of QC-FC correlations relative to other pipelines (e.g., QC-FC significant correlations of ~4-6% for censoring pipelines compared with ~7.5% for ICA-AROMA pipelines as well as a ~10% reduction in median absolute QC-FC). This result is consistent with Ciric et al. (2017), who showed small improvements to QC-FC correlations when applying spike regression to their 36P pipeline (equivalent to our 24HMP+8Phys+4GSR pipeline). Together, our results suggest that the superior performance of censoring pipelines must be balanced with their high cost in terms of participant exclusion and tDOF-loss.

The combination of censoring with temporal filtering should be done with care (Carp, 2013; Power et al., 2013). Applying censoring after bandpass filtering, as per our basic scrubbing pipeline, can cause motion contamination from censored volumes to spread into uncensored volumes (Carp, 2013). The optimized scrubbing method was designed to minimise these effects (Power et al., 2013; Power et al., 2014). Here, we used a Fourier rather than Butterworth temporal filter as used previously. The latter can minimize the spreading effect of motion artefacts (Power et al., 2013), but can produce edge effects. Prior work has addressed this limitation by discarding the first and last 15 volumes of filtered data (Power et al., 2014), but this was not feasible for the datasets examined here. Note that in our analysis the basic scrubbing pipeline applied censoring after temporal filtering, whereas the spike regression pipeline applied censoring before filtering, which (to some extent) covers both orders. We found little difference between them, but refined analyses could focus in more detail on these order effects.

### 4.5 Global signal regression

GSR is one of the most controversial preprocessing steps in the fMRI literature. It has been shown to improve correction for motion-related artefacts and other physiological confounds (Power et al., 2017b) and it improves the spatial specificity and anatomical plausibility of seed-based functional connectivity maps (Fox et al., 2009). However, GSR mathematically forces the distribution of correlations between voxels to be centred around zero, introducing anti-correlations that may be artefactual (Fox et al., 2009).

In our analysis, GSR led to major improvements in QC-FC correlations for HMP pipelines and aCompCor pipelines, but relatively small improvements for ICA-AROMA pipelines. However, inspection of processed time series at a single subject level clearly demonstrates the superiority of GSR in removing global signal fluctuations that have previously been linked to physiological artefacts (Power et al, 2017), even in ICA-AROMA pipelines (see Supplementary material for single subject plots). As previously shown (Ciric et al., 2017), GSR exacerbated QC-FC distance-dependence in all pipelines. This effect was less pronounced when GSR was combined with ICA-AROMA, which may be attributed to ICA-AROMA removing spatially varying motion-related confounds.

Case-control differences in the CNP dataset (with stringent exclusion applied) only emerged when GSR was used. Effective denoising and high sensitivity to detecting clinical differences are desirable characteristics of a pipeline. However, without a ground truth it is difficult to determine whether these case-control results demonstrate that GSR is the only method capable of uncovering “true” group differences in the CNP dataset, or whether they reflect the transforming effect that GSR can have on between-group contrasts of functional connectivity estimates, which has been suggested to lead to spurious group differences (Saad et al., 2012).

The inherent assumption of GSR is that the global signal is mostly dominated by the noise rather than the true neural signal. The problem is that the global signal samples from regions of grey matter. These grey matter signals will contribute to the global signal to the extent that they are coupled with each other. In other words, tightly coupled sets of grey matter voxels (i.e., putative functional networks of brain), will contribute strongly to the global signal. Removal of the global signal will thus subtract the common components of these networks from other parts of the brain, potentially introducing anti-correlations between brain regions that otherwise show weak functional connectivity. In other words, the impact of GSR will depend on the initial covariance structure of grey matter (and WM/CSF) signals. Consistent with this view, Saad et al. (2012) and Gotts et al. (2013) have used both modelling and

experiments to show that the impact of GSR is greatest when two groups differ in the underlying correlation structure of the network. For instance, if the size of one network differs between patients and HCs, the global signal will contain a greater contribution from this network in one group relative the other, causing GSR to affect this system differently between the two groups, and thus potentially resulting in spurious group differences. Critically, Saad et al. (2012) derived an equation for determining the correlation structure of a network post-GSR given its architecture pre-GSR. The model of GSR transformation was later shown to explain >95% of the variance in empirical functional connectivity measures post-GSR (Gotts et al., 2013), demonstrating the explanatory power of this basic mathematical property of GSR. The low consistency in edges identified across pipelines as showing differences between HC and SCZ is consistent with the hypothesis that the effects of GSR are contingent on the initial covariance structure of the functional connectivity matrix, since different denoising pipelines will affect this structure in different ways. This dependence on initial processing may be mitigated to some extent by combining GSR with ICA-AROMA, as group differences were relatively consistent under minor variations of this combination.

While GSR itself has the potential to cause spurious group differences in functional connectivity (Saad et al., 2012), it is also true that failing to remove the global signal can conceivably lead to spurious group differences, particularly when two groups differ in the amount of global noise. For example, Power et al. (2017b) have recently shown that GSR is the only effective method for removing large-scale, global signal fluctuations in fMRI time series that are mainly attributable to respiration. The efficacy of GSR stood in stark contrast to model-based correction methods that used physiological recordings, which were only partially effective in removing these global fluctuations. If two groups differ in respiratory or other physiological processes, which is plausible in patient cohorts who may have heightened anxiety in the scanner environment, then these differences may impact variations in global noise. Furthermore, recent work investigating motion correction strategies in rs-fMRI data from the Human Connectome Project (HCP), which has superior temporal resolution to the data examined in this report, showed that mean grayordinate time series regression (a proxy for GSR focused on grey matter signals) was required alongside ICA-based correction strategies to remove motion-related global artefacts (Burgess et al., 2016). Further theoretical work on the properties of GSR and alternative correction methods will be useful in resolving the concerns associated with this processing step.

### 4.6 Test-retest reliability

In general, pipelines that incurred lower tDOF-loss were associated with higher average TRT, with the simplest noise model – the 6HMP pipeline – showing the highest reliability. Given that this method was generally not successful in removing motion effects, this result suggests that a substantial fraction of reproducible BOLD signal may be driven by head motion and/or other physiological confounds. This conclusion is consistent with prior work indicating that head motion may be associated with a specific spatial pattern of activation, which may be represent a trait-like characteristic (Yan et al., 2013b; 2013a). Twin research has shown that head motion is indeed a heritable trait (Couvy-Duchesne et al., 2014). In particular, covariance between mean translational head motion, maximum translational motion, and mean rotation was related to a single latent head motion construct for which half the variance between people was explained by additive genetic factors. This result may partly explain why successful removal of motion effects reduces TRT.

### 4.7 Sensitivity to clinical differences

The analysis of the BMH dataset revealed no differences between OCD patients and HCs for any pipeline, whereas analysis of the CNP dataset identified differences between SCZ and HCs for several denoising approaches. One interpretation of this discrepancy is that the group differences in the CNP dataset may simply reflect the higher levels of motion in this dataset. However, differences between SCZ and HC were identified even using pipelines such as ICA-AROMA+2Phys and 24HMP+8Phys+4GSR+SpikeReg, that were relatively successful in removing motion-related effects according to our benchmarks. Thus, it is possible that disease-related brain changes are more pronounced in schizophrenia than OCD, at least as measured using the current analysis strategy (i.e., brain-wide comparison of pair-wise functional connectivity in a 333-node or 264-node network). Alternatively, the lack of case-control differences in the BMH data could simply be attributed to lower statistical power relative to the CNP dataset. A troubling result is that the specific subset of connections showing differences appears to be vary considerably depending on the preprocessing pipeline used (see Figures 9 and 10). Such variability will result in a literature littered with inconsistent findings, leading to difficulties in replication and impeding the development of valid models of disease pathophysiology.

### 4.8 Controlling for motion-related artefact through denoising vs participant exclusion

The results of our analysis, as well as those from many other groups, demonstrate a strong case for the use of volume censoring methods to deal with motion-related artefacts. Indeed, we find that censoring methods are very effective at mitigating QC-FC correlations and do a very good job of minimising QC-FC distance-dependence. However, an important distinction needs to be drawn between correcting for motion-related artefact in the BOLD signal of a given individual and controlling for motion-related artefact across a given dataset. The former depends upon the sufficient ‘removal’ of nuisance-related variance from the BOLD signal; the latter can readily be achieved by simply discarding entire BOLD datasets that are contaminated by motion, which in turn reduces the number of participants available for analysis. This trade-off is not trivial because many research groups are not in the position to discard large proportions of participants to achieve adequate control over motion. Indeed, the authors of optimized scrubbing acknowledge that the method is expensive, and deliberately chose to develop it on datasets that could withstand stringent denoising and exclusion (Power et al., 2014). Indeed, in our analysis of the CNP dataset, which is relatively large compared to many schizophrenia fMRI studies in the literature, optimized scrubbing reduced the sample size from n = 171 to n = 85; i.e., a ~50% loss in participants.

In many practical circumstances, one must balance denoising efficacy with power considerations. In the next section, we discuss strategies for examining motion effects when exclusion of large numbers of participants is not possible. Moving forward, it may be that prospective strategies motion correction strategies (Herbst et al., 2013; Maclaren et al., 2012), multi-echo fMRI (Kundu et al., 2013), and/or online methods for monitoring motion to ensure that sufficient non-censored data are acquired during scanning (Dosenbach et al., 2017), may help to mitigate this severe loss of data, but these methods have not yet been extensively applied in the literature. Until such protocols receive widespread application, it may be necessary to tolerate some contamination of motion in order avoid extreme wastage of data. We discuss strategies for evaluating the potential confounding effects of motion in such cases below. Our data suggest that, while optimized scrubbing may provide strong control over motion-related artefact, techniques such as traditional scrubbing or ICA-AROMA may offer reasonable alternatives that prevent extreme loss of data.

### 4.9 Recommendations

The results of our study show that complete control of motion-related artefact is difficult to achieve. We find that censoring-based methods and GSR are very effective. The former can come at high cost in terms of data loss, and it remains unclear whether the latter distorts inter-regional correlations. Across most metrics, ICA-AROMA combined with stringent exclusion criteria showed a marginal performance decrement relative to censoring pipelines, but it did not incur the same high cost, suggesting it may be a more efficient method. It also led to more reliable group differences.

More generally, we recommend a set of steps that should be undertaken regardless of which motion-correction strategy is deployed (Figure 11). This workflow should help mitigate and characterize the effects of motion on functional connectivity and provide sufficient data to allow readers to make their own judgements regarding the success of a particular denoising procedure. The basic workflow involves the following steps. First, inspection of each participant’s processed EPI data as a time series (i.e., “carpet plot”) alongside quality control metrics such as FD and DVARS (Power, 2016). We provide code for this analysis (https://github.com/lindenmp/rs-fMRI) and show several examples in our supplementary materials, but also note that the recently available toolbox, MRIQC (https://github.com/poldracklab/mriqc), provides a user-friendly platform for this purpose.

**Figure 11.**
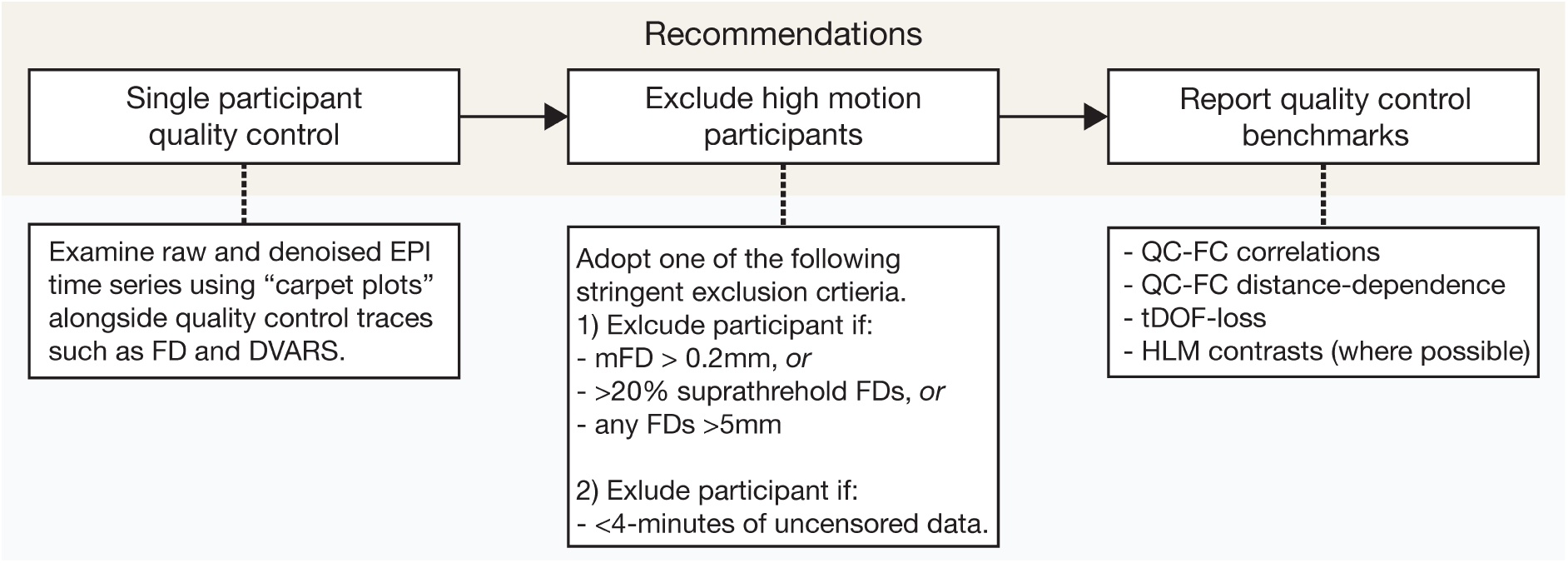
Recommended workflow for characterising and reporting motion characteristics in resting-state fMRI analyses.

The second step is to remove participants with high levels of motion. Our analysis indicates that exclusion of people who have <4-minutes of uncontaminated EPI data dramatically boosts denoising performance. For a TR of 2-2.5, we defined contaminated volumes as those with >0.25mm FD_Jenk_ for our spike regression pipeline and >0.2mm FD_Power_ for our scrubbing pipelines, but data with different TR may require different criteria. Note that these numbers (4-minutes and 0.25mm/0.2mm) are heuristic and should be validated empirically. In many studies, this level of exclusion may not be feasible. In this case, an equivalent to our stringent criteria (i.e., >0.2 mFD, >20% suprathreshold FDs, or any FDs >5mm) may be applied.

The third step of the workflow, regardless of how many people are excluded, is to report benchmark metrics to allow readers to understand the impact of motion on the data. Specifically, we recommend reporting QC-FC correlations, QC-FC distance-dependence, tDOF-loss, and, where sample size permits, group differences in functional connectivity between high- and low-motion HC participants. Even if whole-brain network analysis is not the focus of the study (e.g., seed-based studies, ICA-based canonical network studies), it is straightforward to generate whole-brain, pair-wise functional connectivity matrices using established templates such as those provided by Gordon et. al. (2016) and Power et al. (2011). We provide code to generate these benchmarks.

In a case-control design, it is essential to rule out variations in head motion as driving group differences. In addition to presenting the benchmarks discussed in the current study, a simple initial check would be to determine whether motion is correlated with functional connectivity of the same set of edges that show group differences. If overlap occurs, additional steps may be required to rule out head motion. One option would be to match patients and controls on mFD on a 1:1 basis and repeat the analysis in the matched sub-sample. However, depending on the severity of head motion in the patient group, this may result in an under-powered or unrepresentative sample. Another option would be to use a constrained permutation testing approach, in which observed group differences in functional connectivity are evaluated against a null hypothesis generated from the permutation of group labels such that mean case-control differences in mFD are preserved (van den Heuvel et al., 2017). The resulting hypothesis test thus evaluates group differences in functional connectivity that cannot be accounted for group differences in average motion. One caveat is that, if the difference in mFD between the groups is large, there may not be enough degrees of freedom to generate sufficiently randomised data.

### 4.10 Limitations

The lack of a ground truth makes benchmarking the effectiveness of any noise correction strategy difficult. We compensated for this limitation by conducting a comprehensive investigation of noise correction pipelines, looking for convergence between QC-FC benchmark analyses, loss in temporal degrees of freedom, within- and between-session reliability, and sensitivity to clinical group differences in functional connectivity. Nonetheless, we cannot distinguish true positive/negatives from false positives/negatives in case-control analyses with absolute certainty.

We focused here on simple denoising procedures that are readily available and which can be applied to any dataset. Alternative, prospective methods for motion correction that we did not consider include acquiring multi-echo data (Kundu et al., 2013) and actual monitoring of head motion in the scanner (Herbst et al., 2013; Maclaren et al., 2012). These techniques have shown great promise and provide a fruitful way of addressing the problem of head motion in future studies.

The benchmarks used in this study and others (e.g., Ciric et al., 2017) are derived from a framewise quality control measure. By definition, any framewise measure is dependent upon the sampling rate of the data. In the case of framewise displacement, the magnitude of this QC metric will vary as a function of repetition time of the BOLD data, meaning that different FD thresholds for identifying motion-related spikes may be required for data with different TRs. In our analysis, we used suggested FD exclusion/censoring criteria as initially reported by others (Power et al., 2014; 2013; Satterthwaite et al., 2013) and did not adjust for differences in TR, consistent with common practice in the literature. While the data we analyse here have similar TRs to those studied by Power and colleagues, the spike-regression-based criteria were developed using data acquired at ~3s TR, and so may not be directly applicable to our data. We note however, that despite the spike regression criteria being more lenient, spike regression-based methods often showed similar performance to those based on scrubbing, so it seems unlikely that differences in TR could account for our findings; if anything, they may have under-estimated the performance of spike regression. At present, the effect of variable TR on FD, and how it relates to data denoising and benchmarking, is not yet well characterised and will be important to address in future research.

We benchmarked the relative cost of each of our noise correction methods by examining the tDOF used as a result of nuisance regression. We estimated tDOF-loss as the number of nuisance regressors used by each pipeline, and where applicable, each participant, separately. Note that this is a simple approach following the method of Pruim et al. (2015b) as well as Yan et al. (2013a), and that it does not account for autocorrelation in the data (Patel and Bullmore, 2016).

Other methods that can be applied to any data, and which we have not considered here, are ICA-FIX (Griffanti et al., 2014; Salimi-Khorshidi et al., 2014) and wavelet despiking (Patel et al., 2014). However, recent work by Burgess et al., (2016) demonstrated that ICA-FIX on its own failed to sufficiently correct for the global artefacts present in data from the HCP, and that some form of GSR (in this case, mean grayordinate time series regression) was also required. A further limitation of ICA-FIX is that it requires curation of a training dataset with the same acquisition protocol as the analysis sample. Therefore, it requires relatively large samples so that a subset of individuals can be held out as the training set. Wavelet despiking does not suffer from this limitation, and has been shown to be more effective than volume censoring in removing the effects of head motion (Patel et al., 2014). However, the technique does not explicitly incorporate a model of signal noise and requires the setting of a free parameter that must be adapted to different kinds of data. We focused here on relatively automated methods that require little user input.

Finally, we did not examine the efficacy of these denoising pipelines for cleaning multiband fMRI data. Such data can have a distinct noise structure, which can be characterised with ICA as components with sparse and evenly spaced slices (Griffanti et al., 2016), but which are not likely to be identified as noise by the restricted feature set used in ICA-AROMA. Thus, for multiband data, techniques such as ICA-FIX, in which such noise components can be explicitly identified by users, may be more appropriate.

### 4.11 Conclusions

In summary, we comprehensively examined a wide variety of commonly adopted noise correction methods for resting state fMRI data. We found that no single method was able to completely mitigate motion-relate artefact in any of the datasets examined herein. The best performing pipelines included either volume censoring or ICA-AROMA, with censoring-based pipelines performing best. However, superior performance for censoring-based methods came at the cost of substantial data loss compared to ICA-AROMA, which was more cost-effective. Our work contributes to the growing body of literature (Ciric et al., 2017; Power et al., 2015; Satterthwaite et al., 2013; Yan et al., 2013a) highlighting the suboptimal performance of many common noise correction methods used in the literature. Crucially, our results underscore the importance of reporting the residual relationship between in-scanner movement and functional connectivity alongside case-control comparisons in clinical neuroimaging studies. Reporting QC-FC benchmarks will assist investigators and readers to understand which group differences are more likely to be true positives and which may be spurious.

## Acknowledgements

L.P. was supported by an Australian Postgraduate Award and the David Winston Turner Endowment Fund. B.F. was supported by a National Health and Medical Research Council Early Career Fellowship (ID: 1089718). M.Y. was supported by a National Health and Medical Research Council Fellowship (ID: 1117188), Monash University and the David Winston Turner Endowment Fund. A.F. was supported by an Australian Research Council Future Fellowship (ID: FT130100589) and National Health and Medical Research Council Project grants (ID: 3251213, 3251250, 3251392). We would like to acknowledge the reviewers of this manuscript for their thoughtful and detailed feedback.

## Supplementary material for *an evaluation of the efficacy, reliability, and sensitivity of motion correction strategies for resting-state functional MRI*

### White matter and cerebrospinal fluid mask erosion

As mentioned in the methods section of the main text, signals extracted from un-eroded white matter (WM) and cerebrospinal fluid (CSF) masks can be highly correlated to the mean grey matter (GM) signal. In turn, these un-eroded signals have the potential to have a similar impact on the BOLD data as global signal regression. Following recommendations from Power et al., (2017), we performed several iterations of erosions on the WM and CSF masks (Figure S1) and examined how their mean signal correlated to the mean cortical GM signal in the CNP dataset. We extracted the mean GM signal from the cortical ribbon generated using Freesurfer 6.0.

For the majority of pipelines, mean WM/CSF/GM signals were extracted from spatially unsmoothed BOLD data. However, for all of our ICA-AROMA pipelines, mean WM/CSF/GM extraction was performed on smoothed data. Thus, we examined the correlation between WM/CSF and GM for both smoothed and unsmoothed BOLD data. Table S1 shows the correlation between the WM and CSF signals and grey matter signals as a function of increasing erosion (up to 5 erosions for WM and up to 2 erosions for CSF) averaged over the healthy control participants in the CNP datasets.

**Table S1.** Correlation between mean white matter/cerebrospinal fluid and mean cortical grey matter averaged across healthy control participants from the CNP datasets. eN denotes the number of erosion iterations performed.

|  | WM-e1 | WM-e2 | WM-e3 | WM-e4 | WM-e5 | CSF-e1 | CSF-e2 |
| --- | --- | --- | --- | --- | --- | --- | --- |
|  | Mean±SD | Mean±SD | Mean±SD | Mean±SD | Mean±SD | Mean±SD | Mean±SD |
| Unsmoothed | 0.87±0.07 | 0.69±0.14 | 0.59±0.18 | 0.55±0.19 | 0.53±0.19 | 0.21±0.29 | 0.16±0.27 |
| Smoothed | 0.83±0.14 | 0.71±0.16 | 0.60±0.19 | 0.53±0.20 | 0.51±0.21 | 0.18±0.30 | 0.15±0.30 |

**Figure S1.**
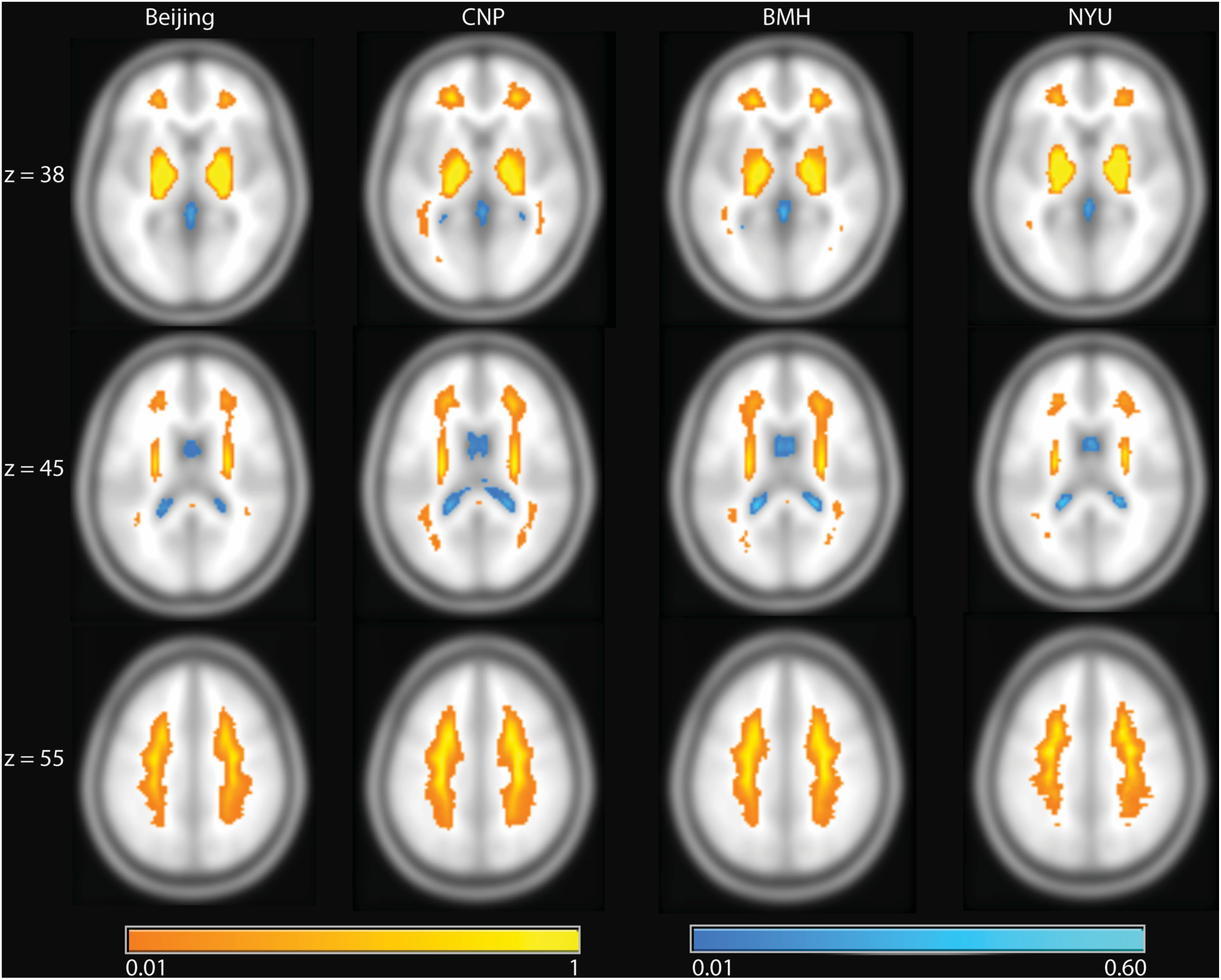
Probabilistic tissue mask used for extraction of physiological nuisance regressors. White matter (orange) and cerebrospinal fluid (blue) masks are depicted here after undergoing five and two rounds of erosions, respectively. Probabilities are calculated across whole samples in each of four different datasets: (from left to right): Beijing, CNP, BMH, and NYU.

**Figure S2.**
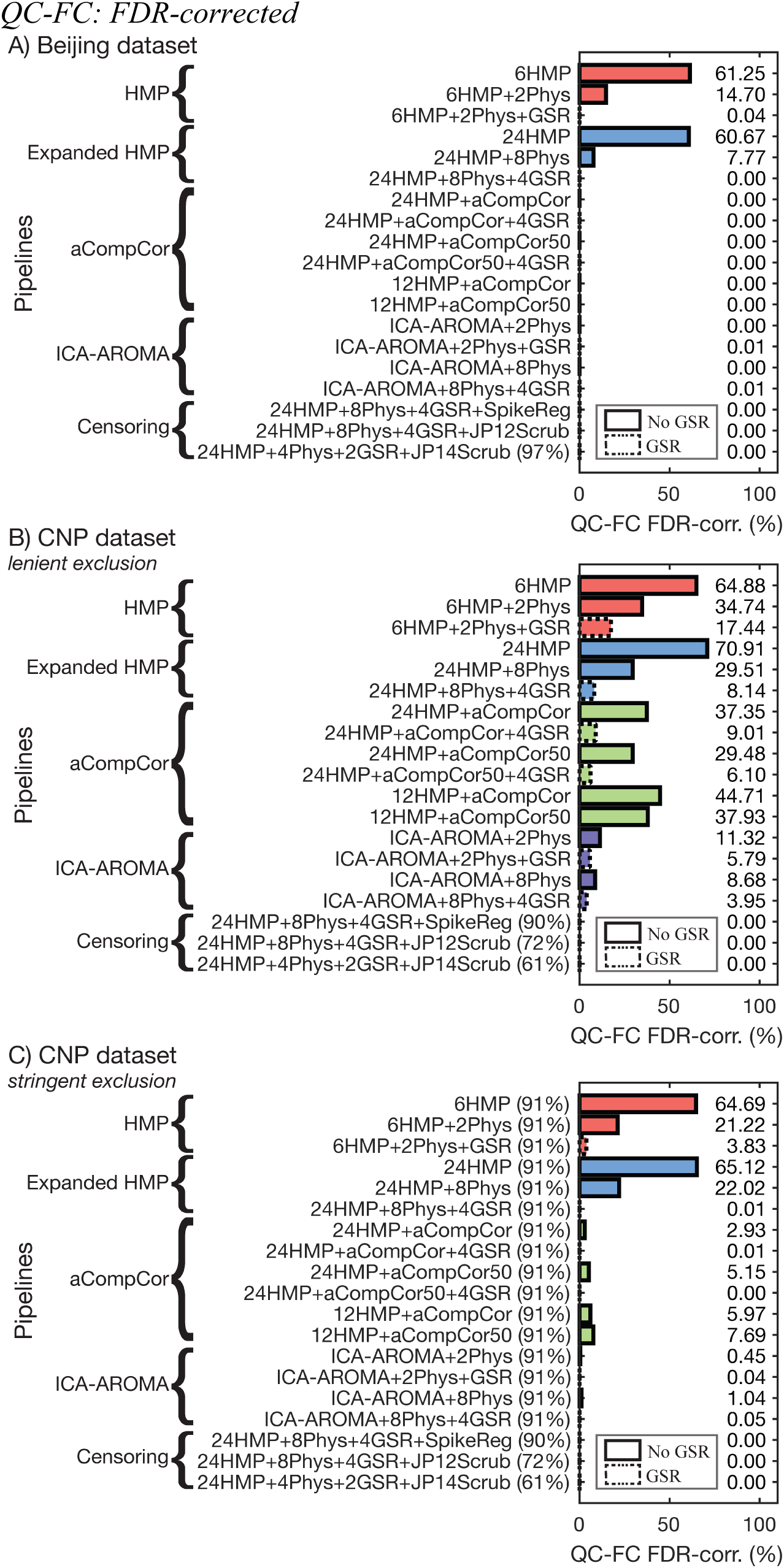
The residual effect of in-scanner motion on functional connectivity after noise correction with one of nineteen different rfMRI pre-processing pipelines for the Beijing (*A*) and CNP dataset under lenient (*B*) and stringent (*C*) exclusion criteria, corrected for multiple comparisons with FDR *p*<0.05. Percentages in parentheses next to pipeline labels are the proportion of HCs retained in analysis. Absent parentheses indicate that 100% of HCs were retained. Dotted lines around the horizontal bars denote pipelines that incorporated global signal regression (GSR).

**Figure S3.**
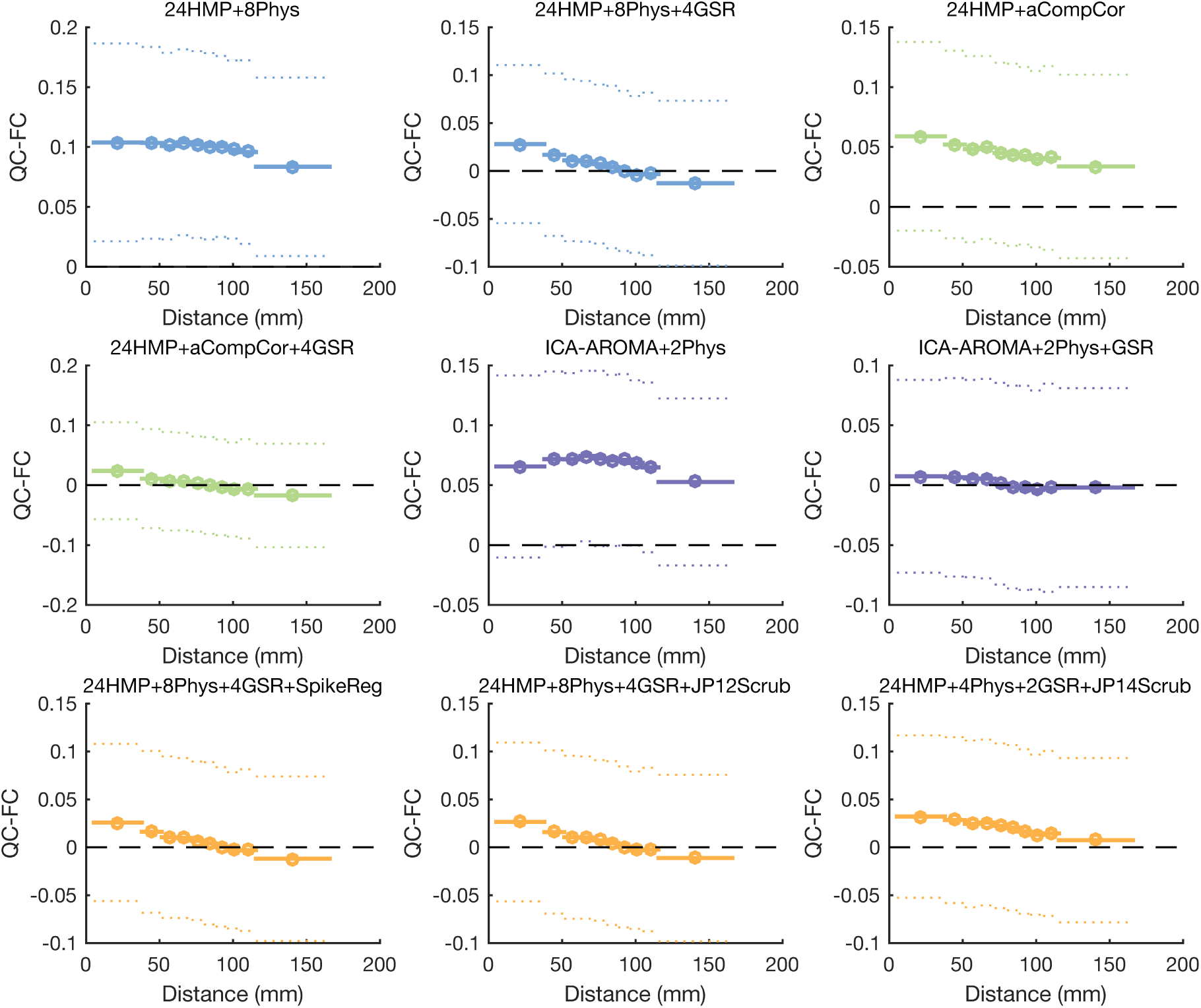
QC-FC distance-dependence for a selection of pipelines applied to the *Beijing* dataset under *lenient* exclusion criteria.

**Figure S4.**
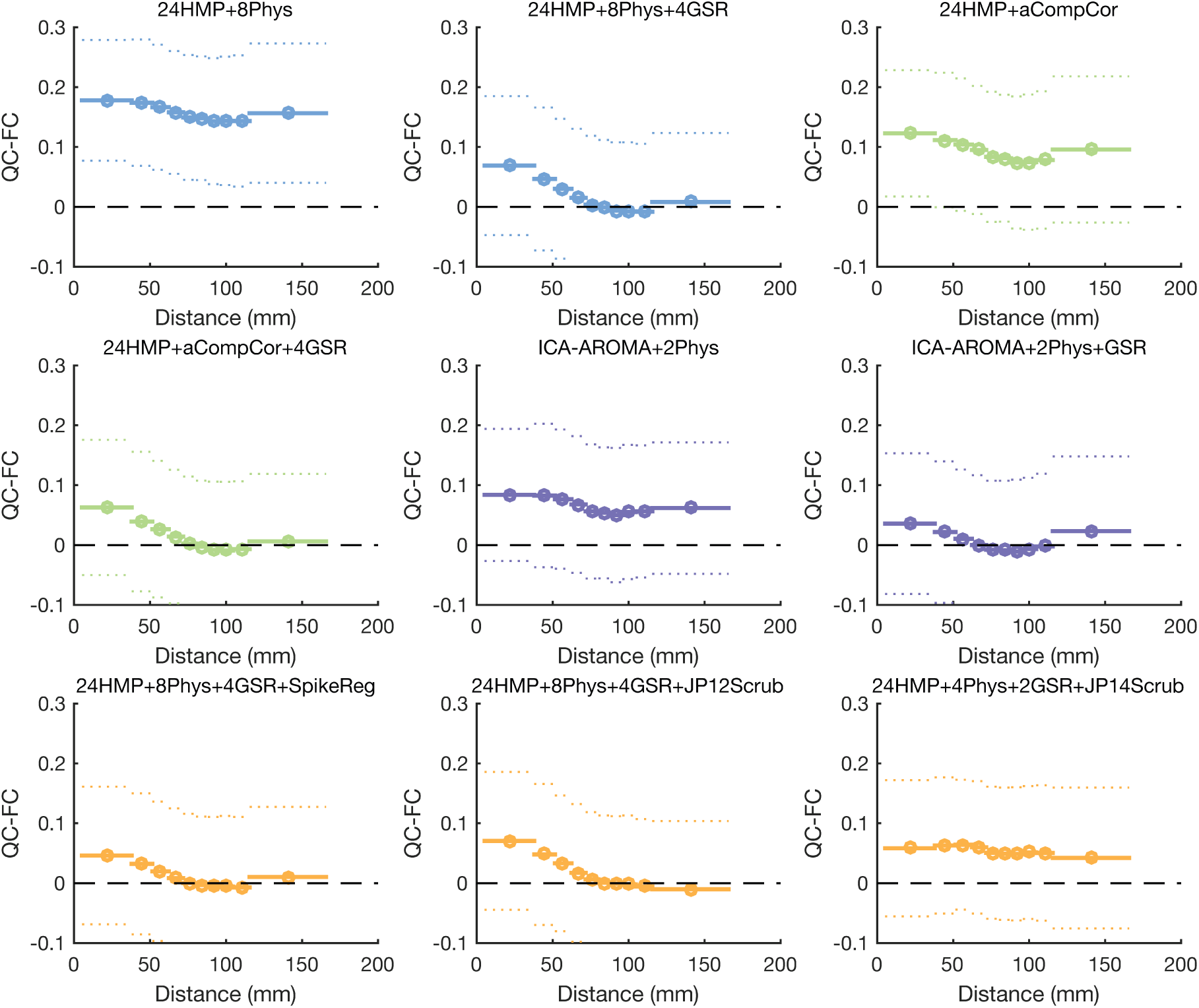
QC-FC distance-dependence for a selection of pipelines applied to the *CNP* dataset under *stringent* exclusion criteria.

**Figure S5.**
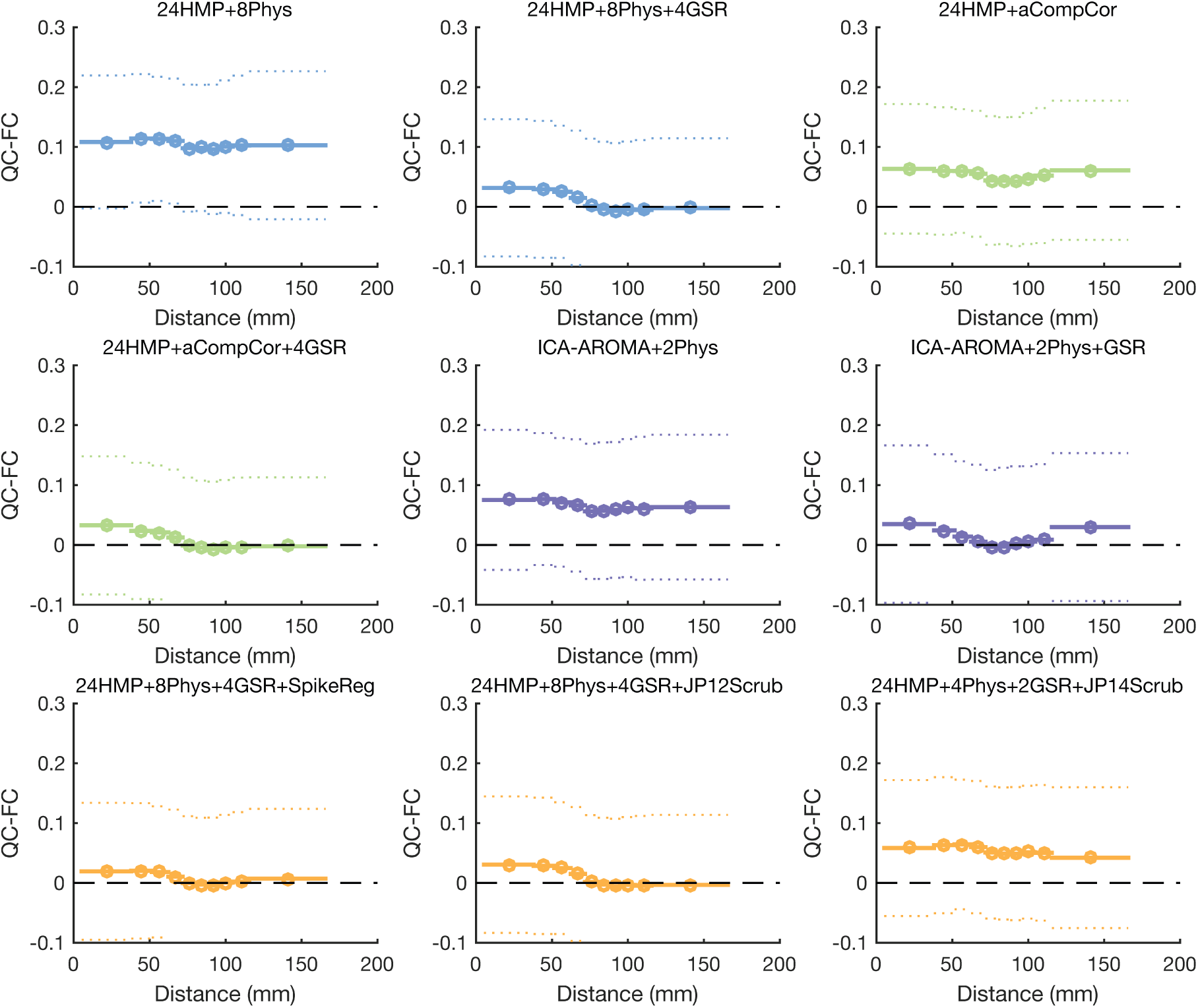
QC-FC distance-dependence for a selection of pipelines applied to the *CNP* dataset under *optimized scrubbing* exclusion criteria. *QC-FC: BMH Dataset*

**Figure S6.**
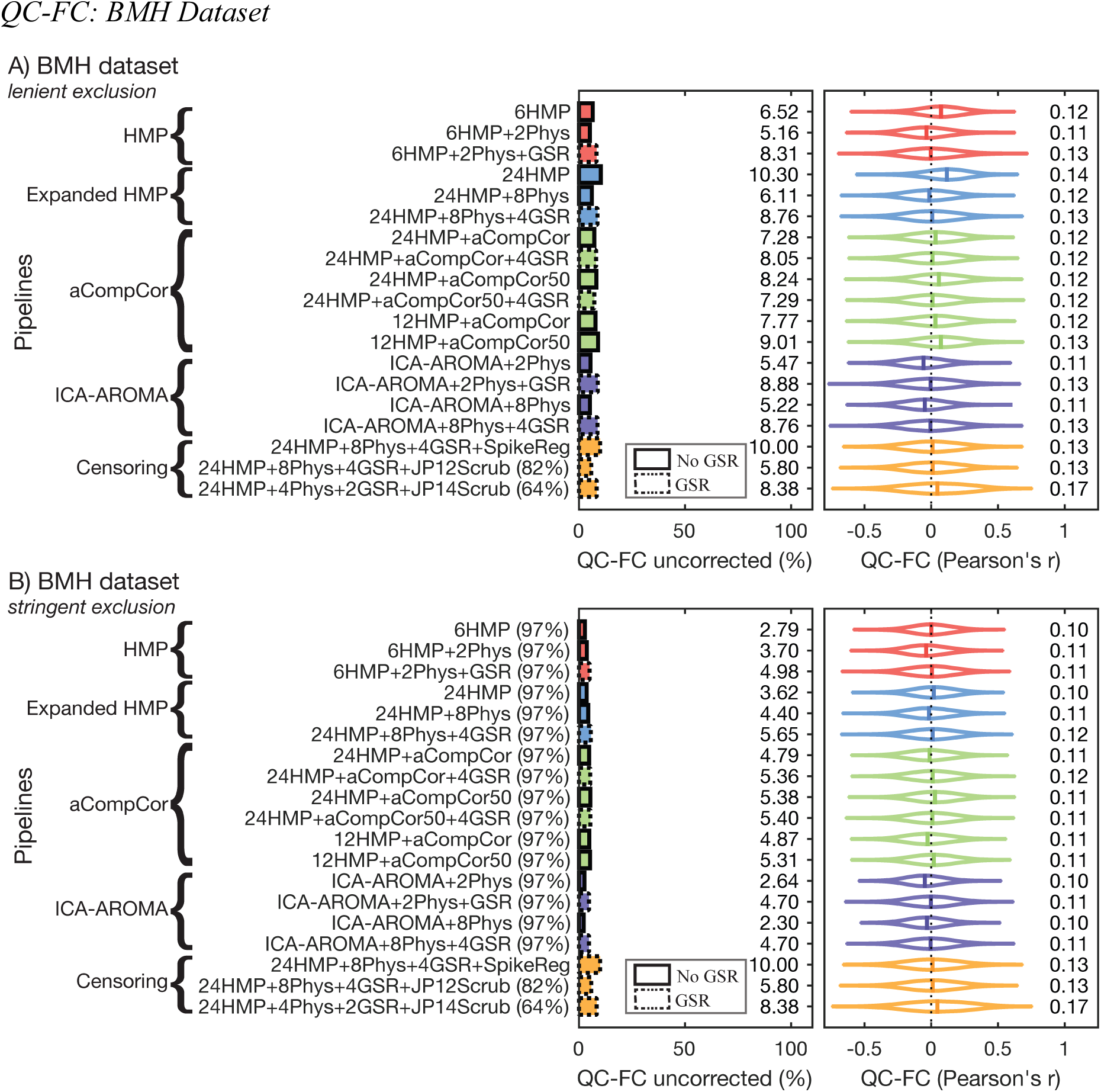
The residual effect of in-scanner motion on functional connectivity after noise correction with one of nineteen different rfMRI pre-processing pipelines for the *BMH* dataset under (A) lenient and (B) stringent exclusionary criteria. Percentages in parentheses next to pipeline labels are the proportion of HCs retained in analysis. Absent parentheses indicate that 100% of HCs were retained. Dotted lines around the horizontal bars denote pipelines that incorporated global signal regression (GSR).

**Figure S7.**
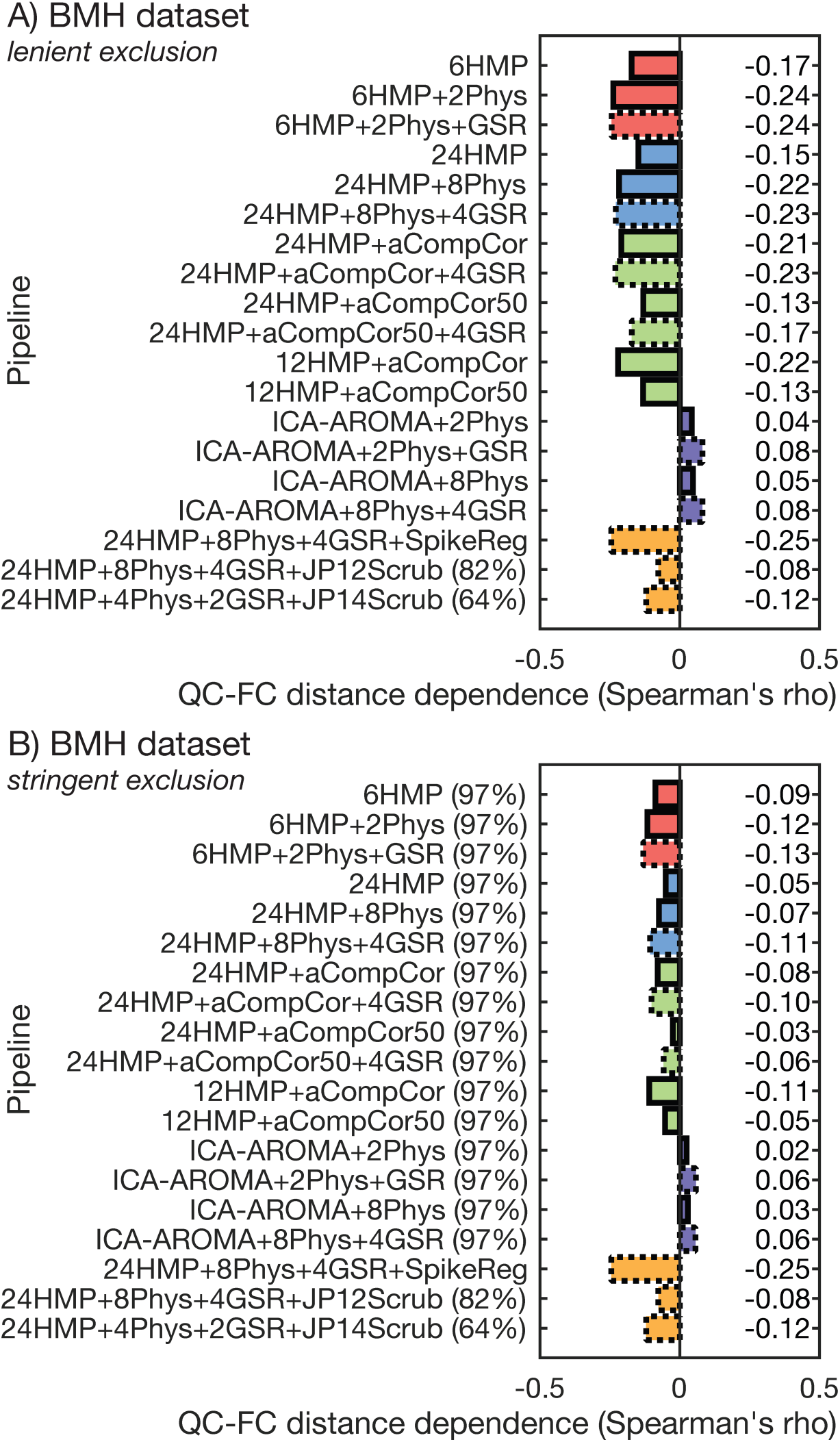
The distance-dependence of QC-FC correlations following application of each pipeline to the *BMH* dataset under (A) lenient and (B) stringent exclusionary criteria. Percentages in parentheses next to pipeline labels represent the proportion of HCs retained in analysis (absent parentheses indicate that 100% of HCs were retained).

**Figure S8.**
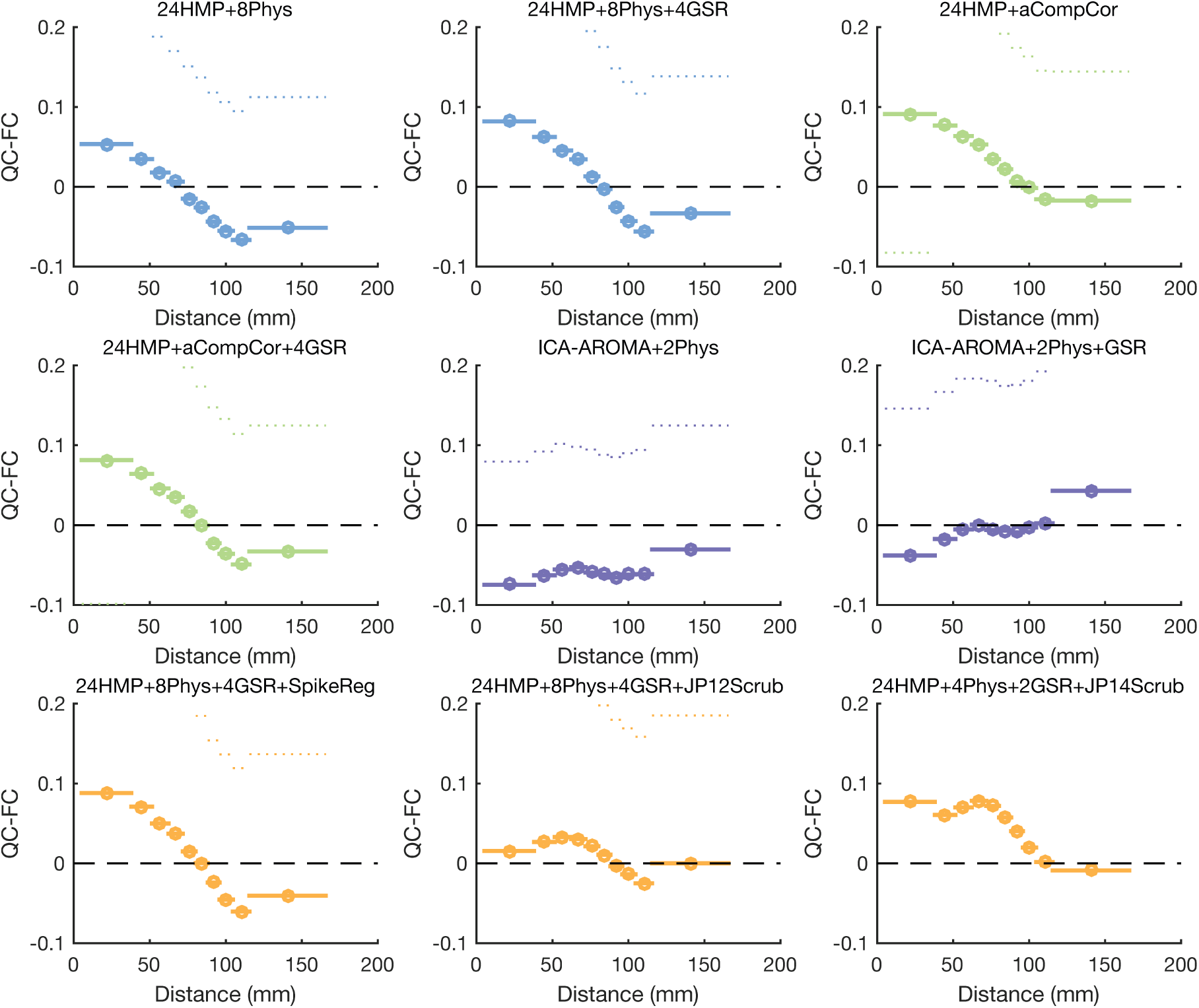
QC-FC distance-dependence for a selection of pipelines applied to the *BMH* dataset under *lenient* exclusion criteria.

**Figure S9.**
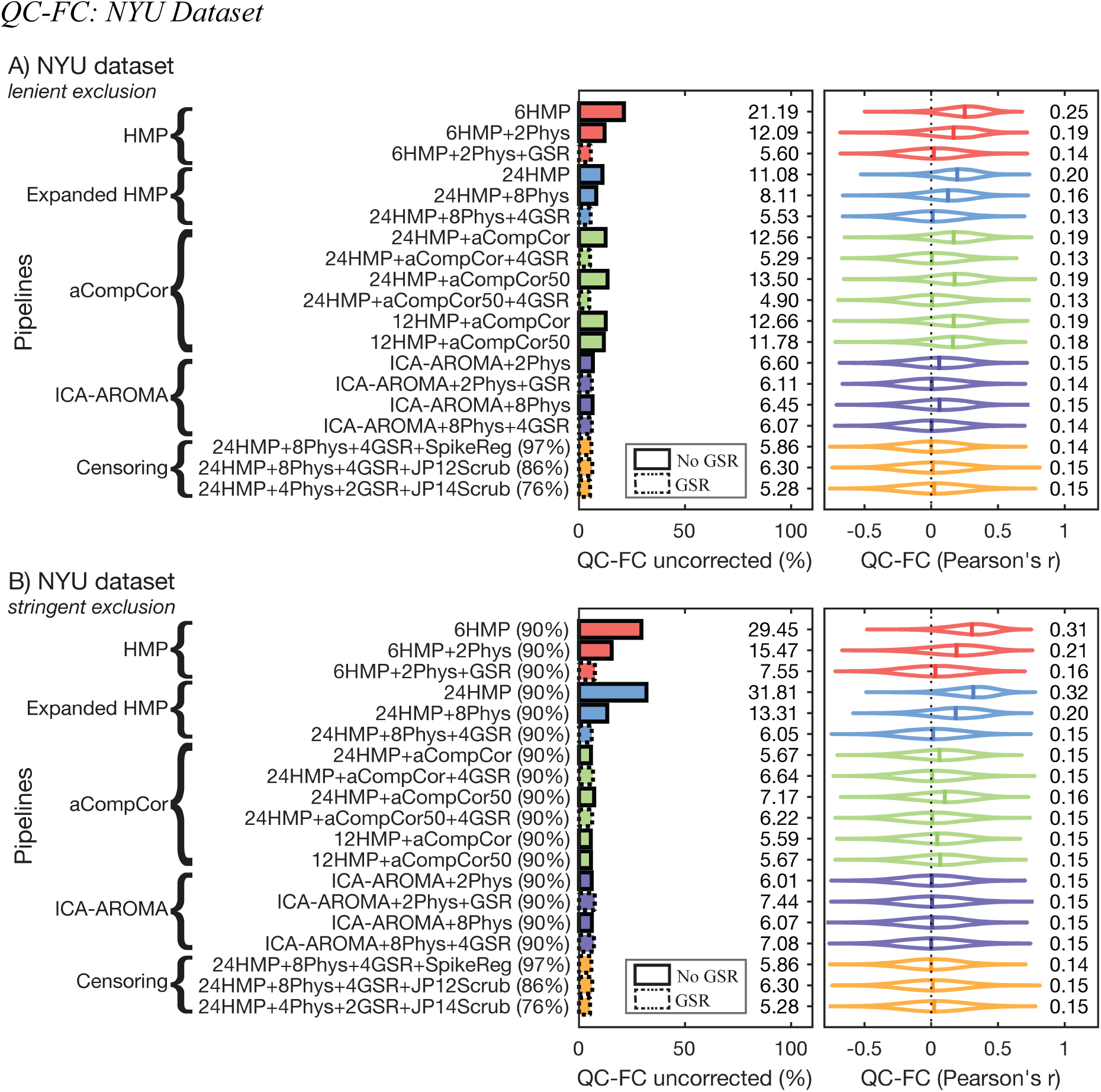
The residual effect of in-scanner motion on functional connectivity after noise correction with one of nineteen different rfMRI pre-processing pipelines for the *NYU* dataset under (A) lenient and (B) stringent exclusionary criteria. Percentages in parentheses next to pipeline labels are the proportion of HCs retained in analysis. Absent parentheses indicate that 100% of HCs were retained. Dotted lines around the horizontal bars denote pipelines that incorporated global signal regression (GSR).

**Figure S10.**
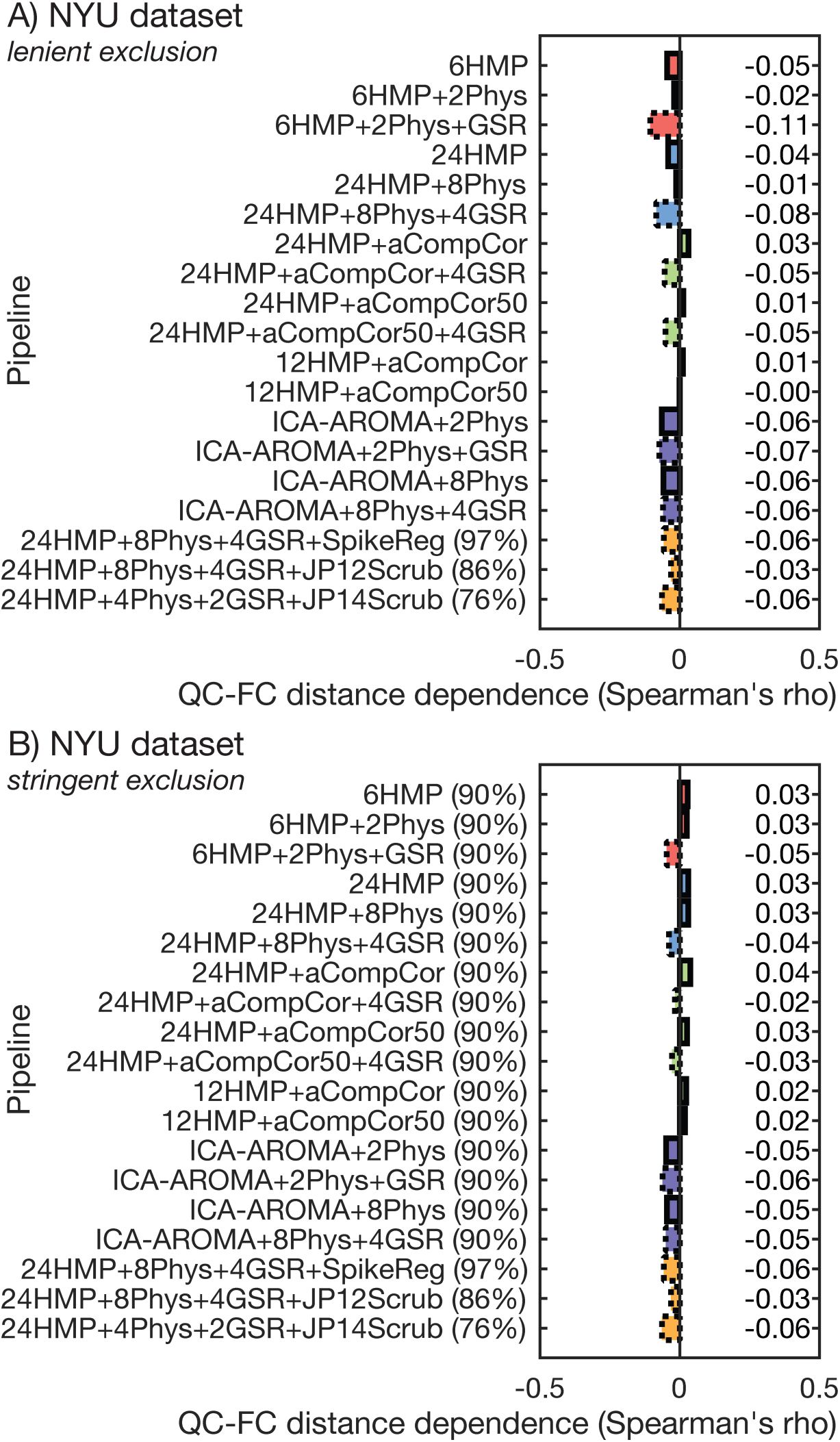
The distance-dependence of QC-FC correlations following application of each pipeline to the *NYU* dataset under (A) lenient and (B) stringent exclusionary criteria. Percentages in parentheses next to pipeline labels represent the proportion of HCs retained in analysis (absent parentheses indicate that 100% of HCs were retained).

**Figure S11.**
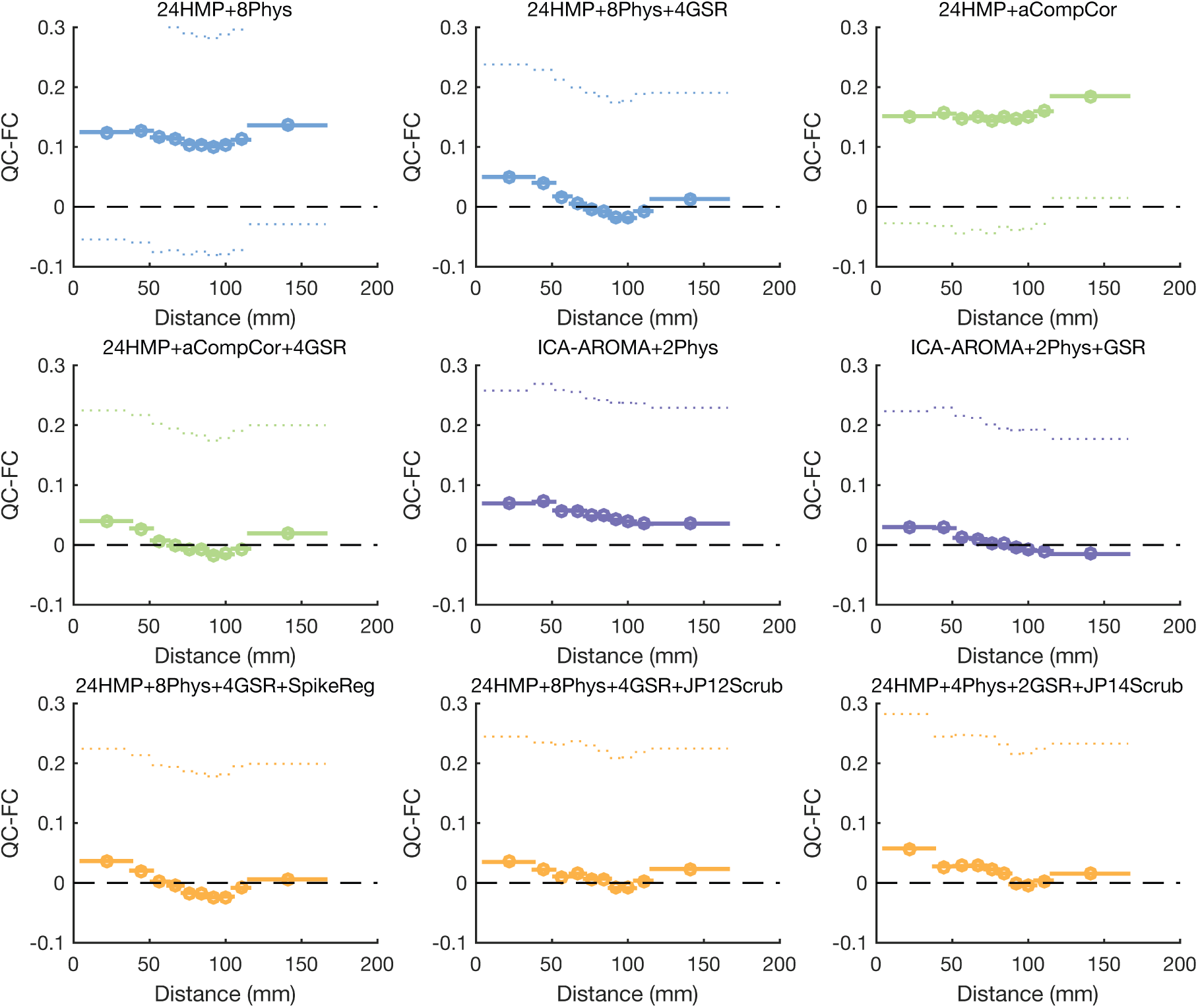
QC-FC distance-dependence for a selection of pipelines applied to the *NYU* dataset under *lenient* exclusion criteria.

**Figure S12.**
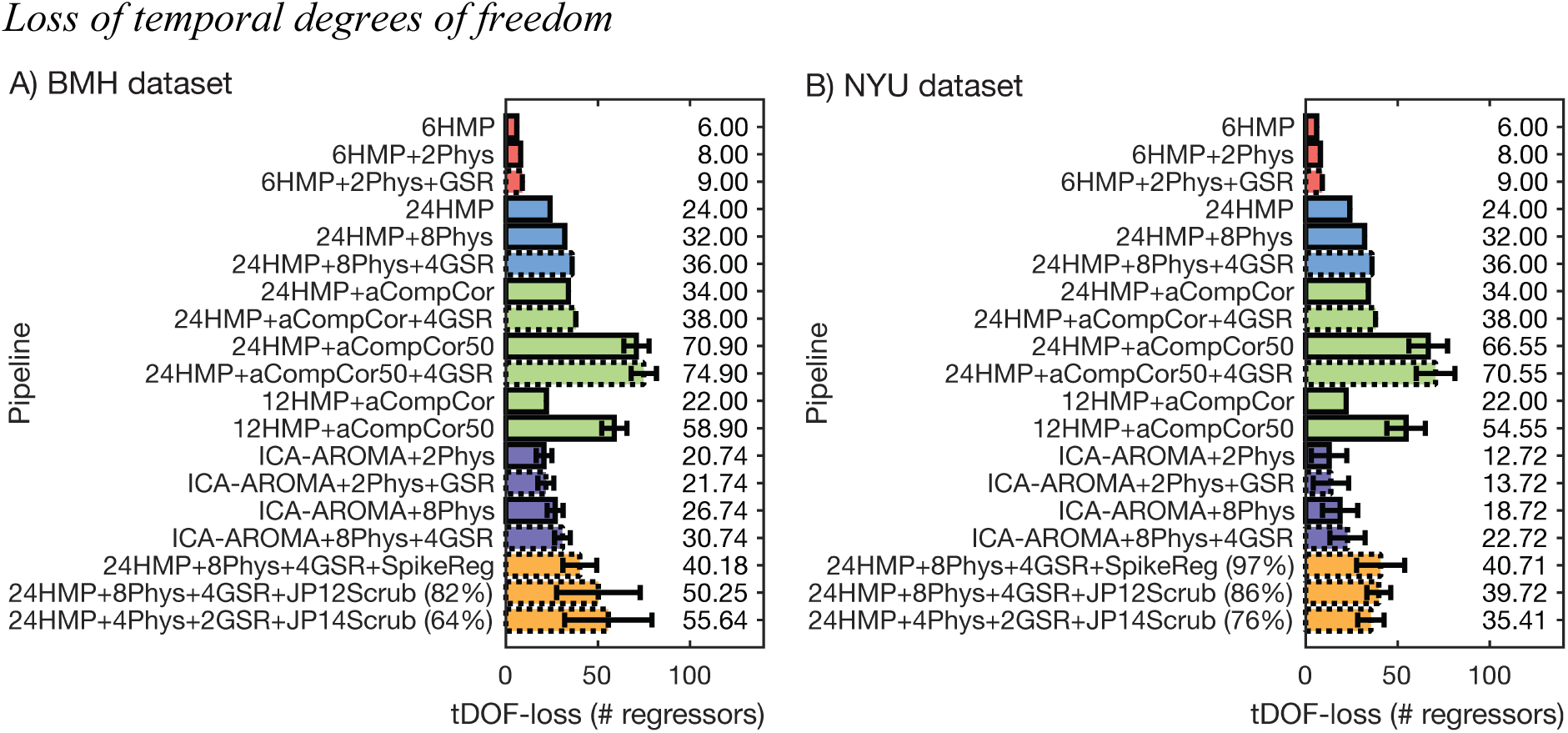
Different fMRI noise correction methods reduce tDOF by different amounts in the (A) BMH and (B) NYU data. Percentages in parentheses next to pipeline labels represent the proportion of HCs retained in analysis (absent parentheses indicate that 100% of HCs were retained).

**Figure S13.**
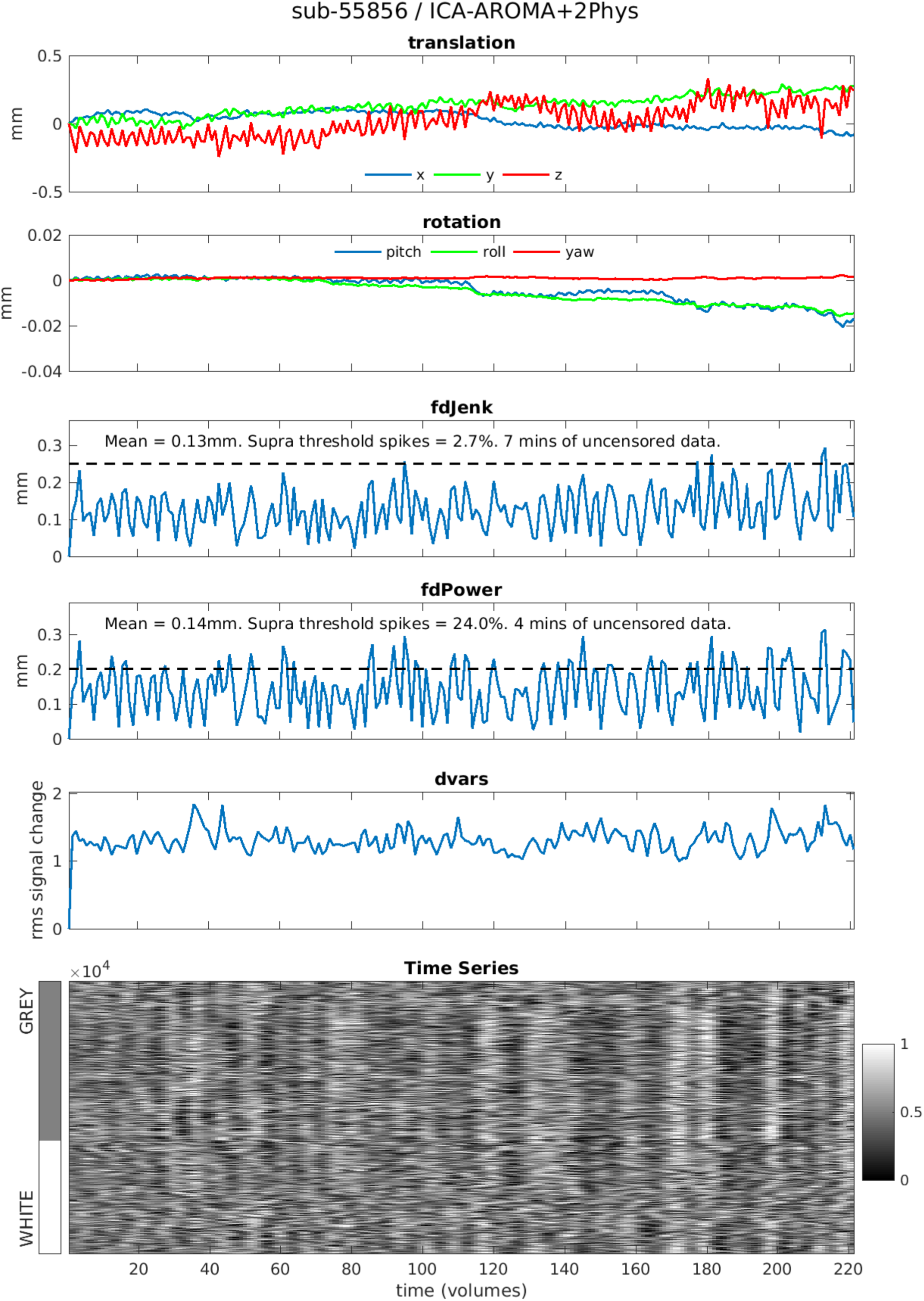
*Highest motion* participant from *Beijing* dataset processed using *ICA-AROMA+2Phys*

**Figure S14.**
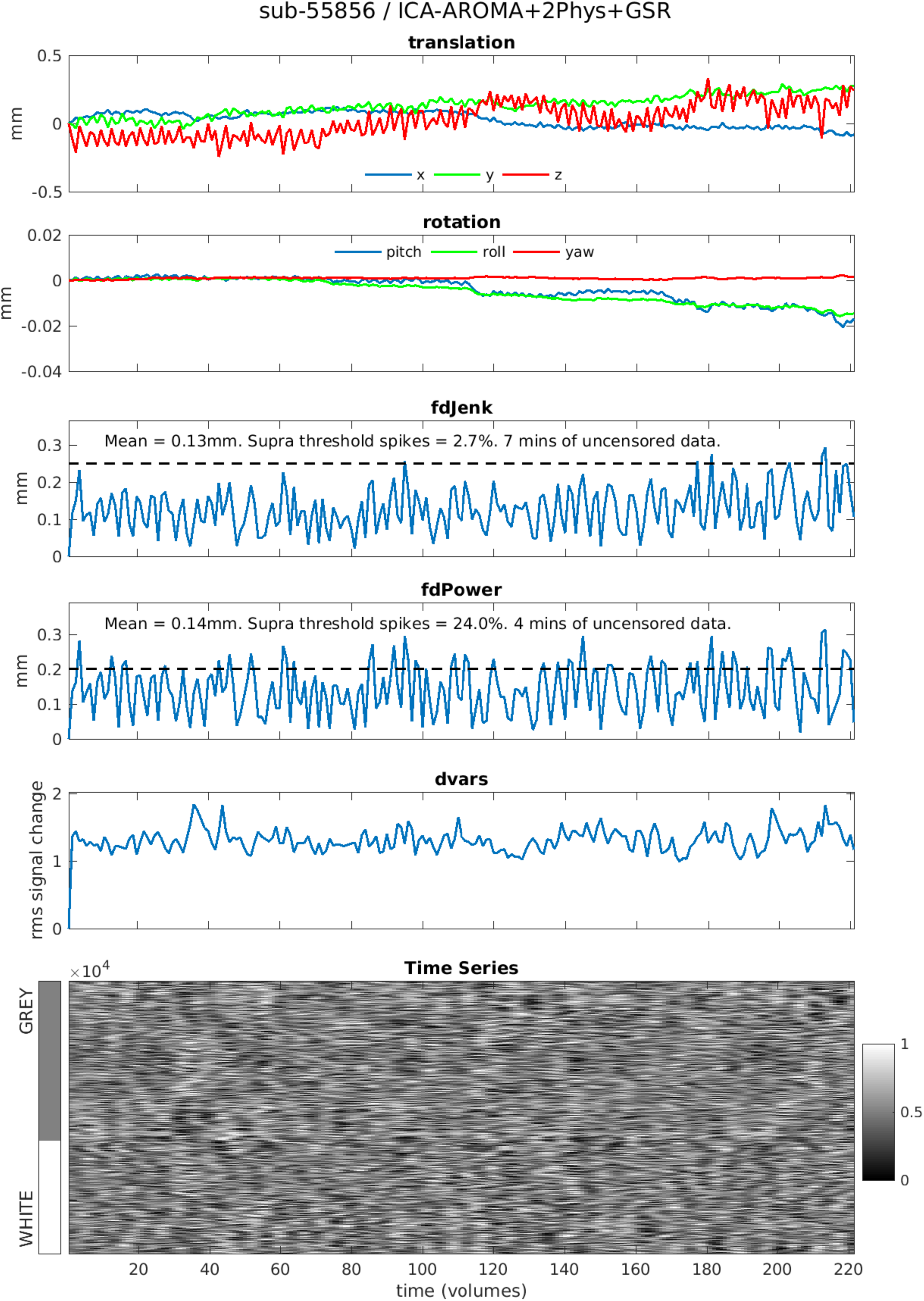
*Highest motion* participant from *Beijing* dataset processed using *ICA-AROMA+2Phys+GSR*

**Figure S15.**
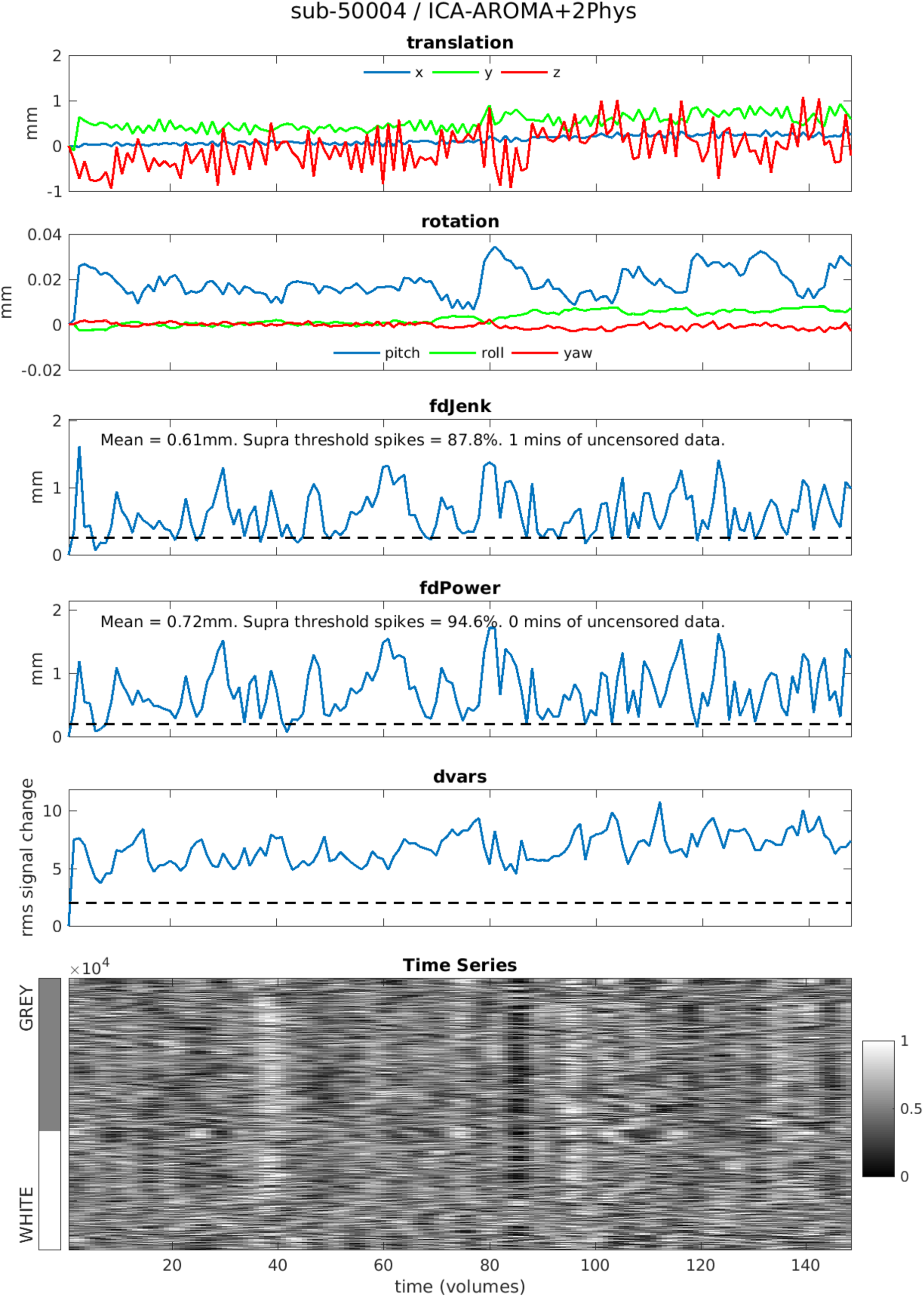
*Highest motion* participant from *CNP* dataset processed using *ICA-AROMA+2Phys*

**Figure S16.**
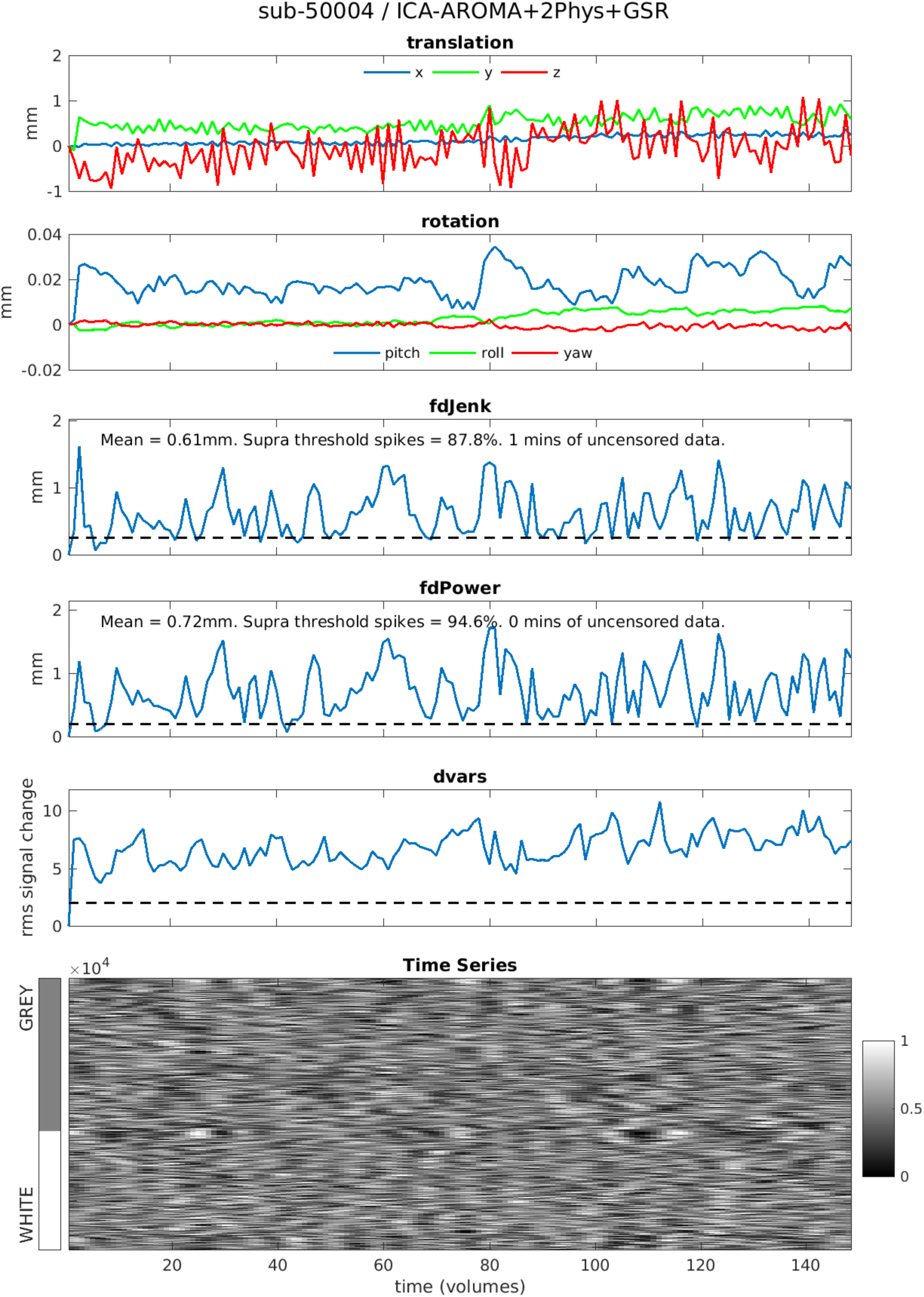
*Highest motion* participant from *CNP* dataset processed using *ICA-AROMA+2Phys+GSR*

**Figure S17.**
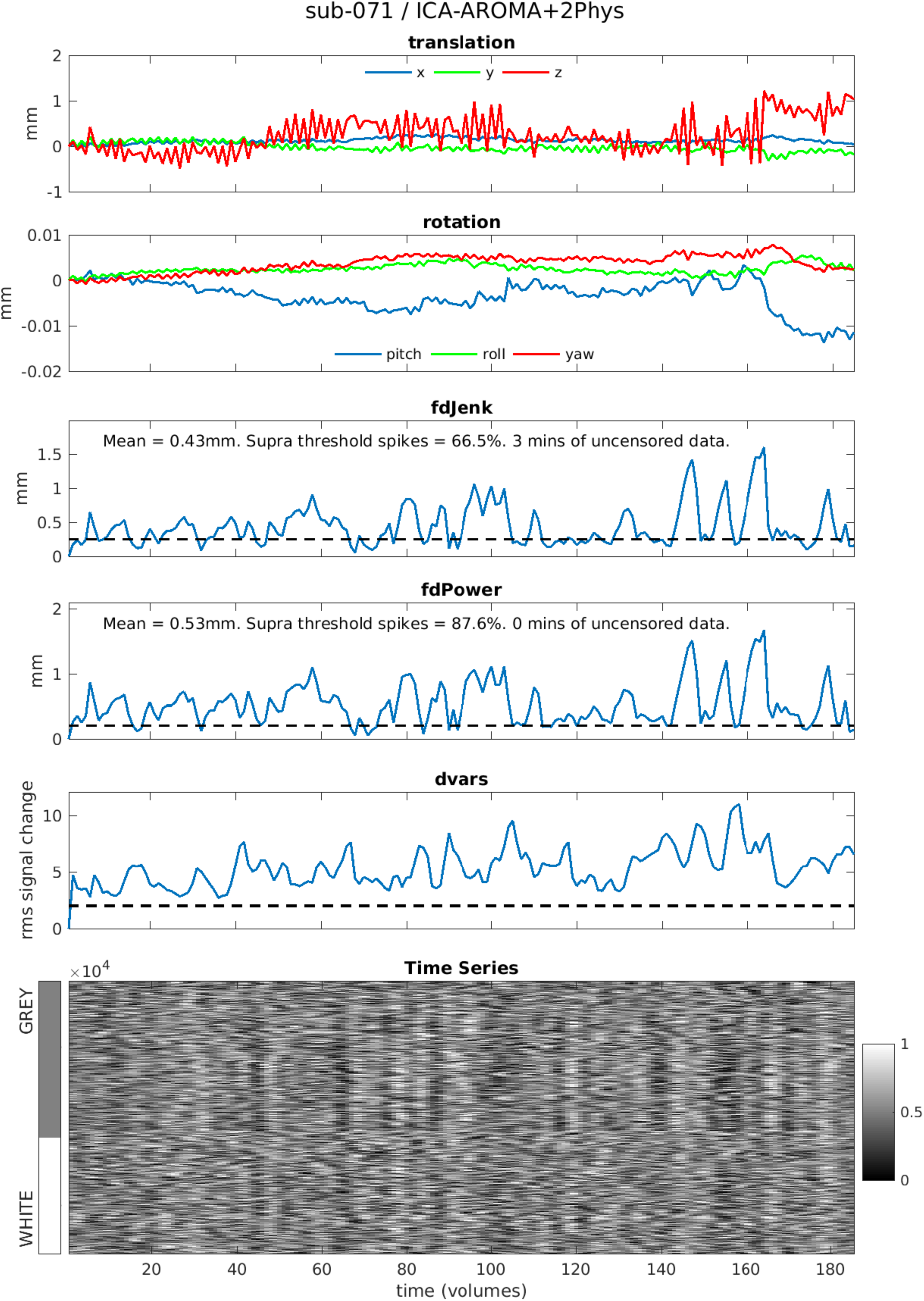
*Highest motion* participant from *BMH* dataset processed using *ICA-AROMA+2Phys*

**Figure S18.**
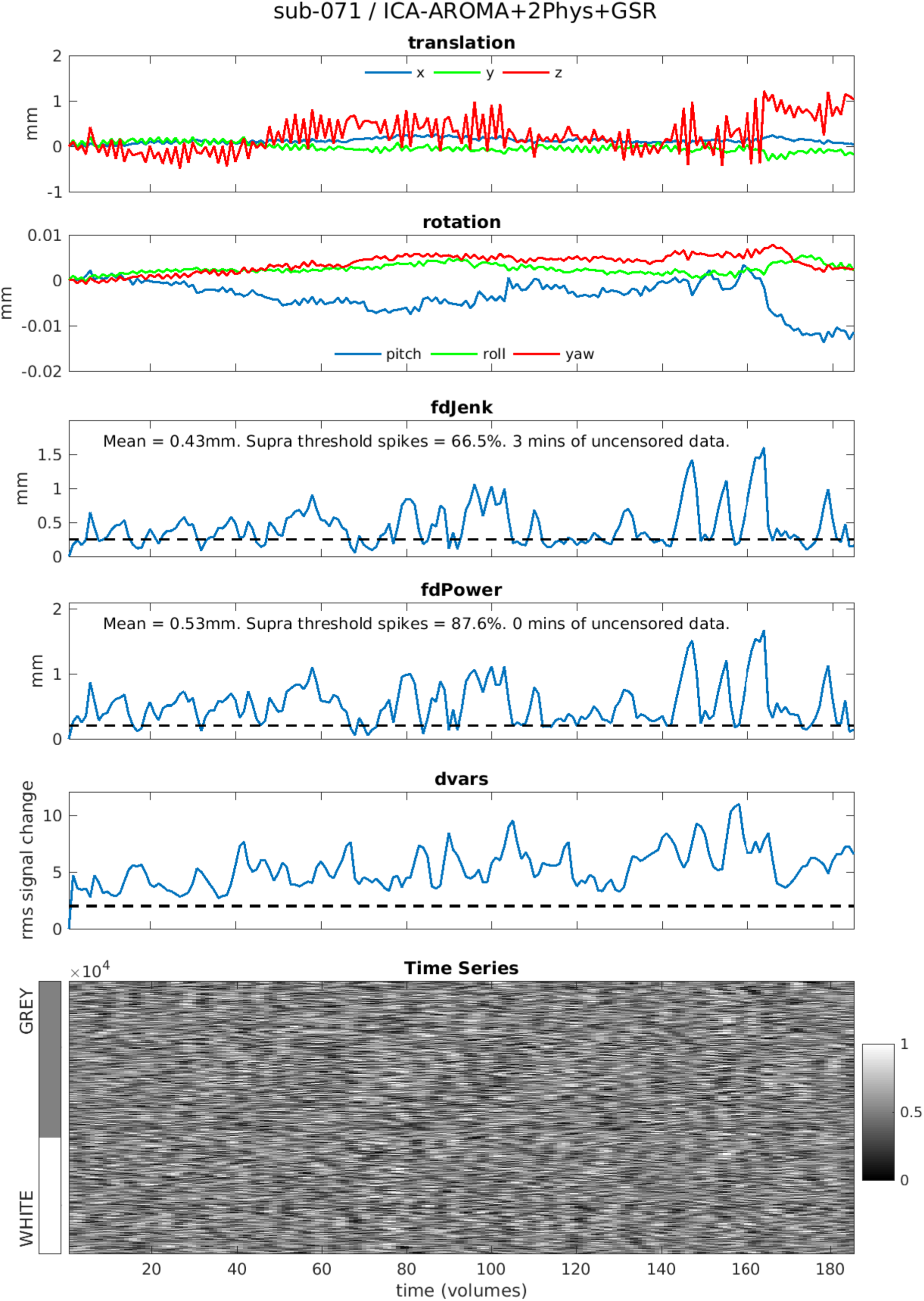
*Highest motion* participant from *BMH* dataset processed using *ICA-AROMA+2Phys+GSR*

**Figure S19.**
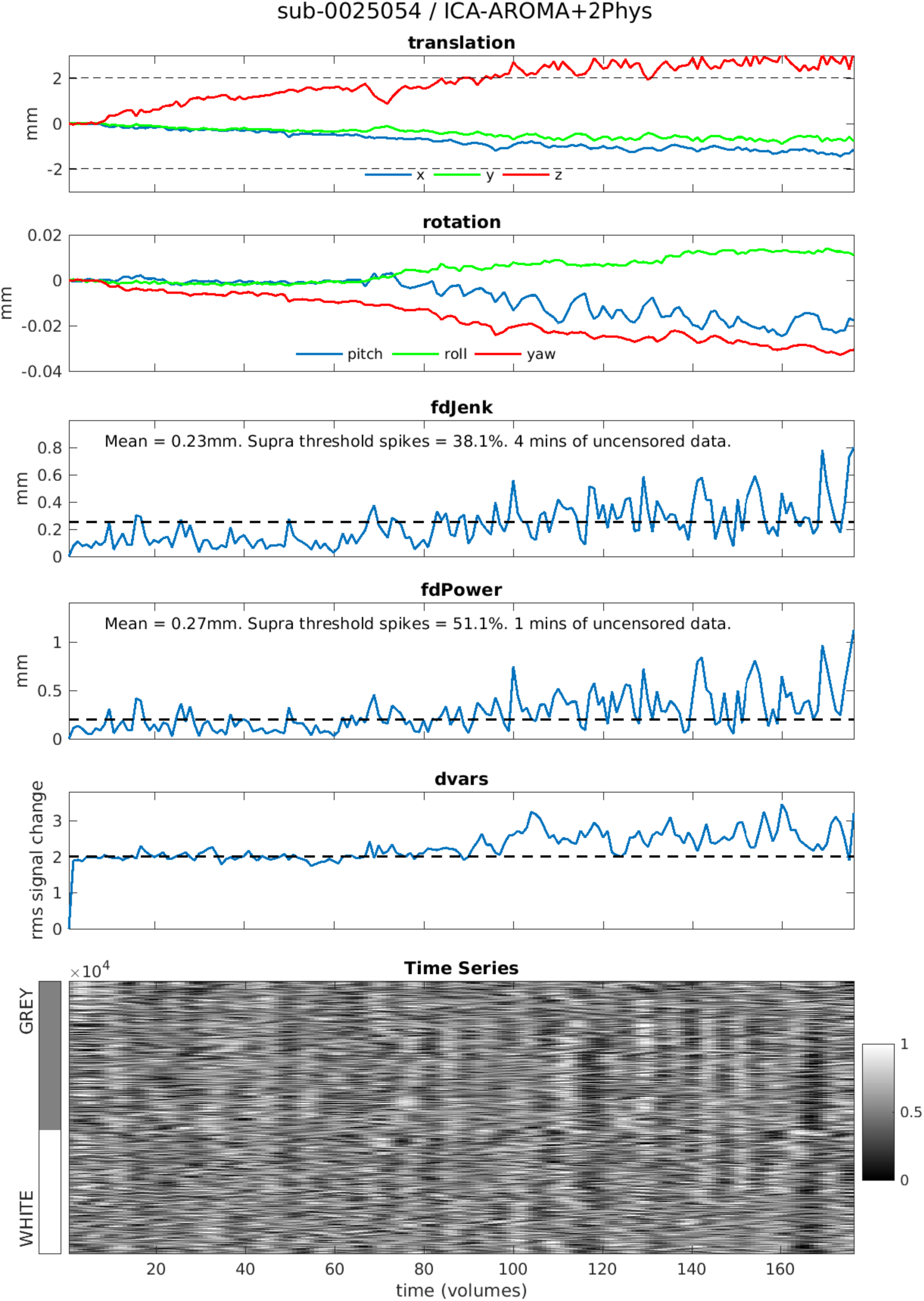
*Highest motion* participant from *NYU* dataset processed using *ICA-AROMA+2Phys*

**Figure S20.**
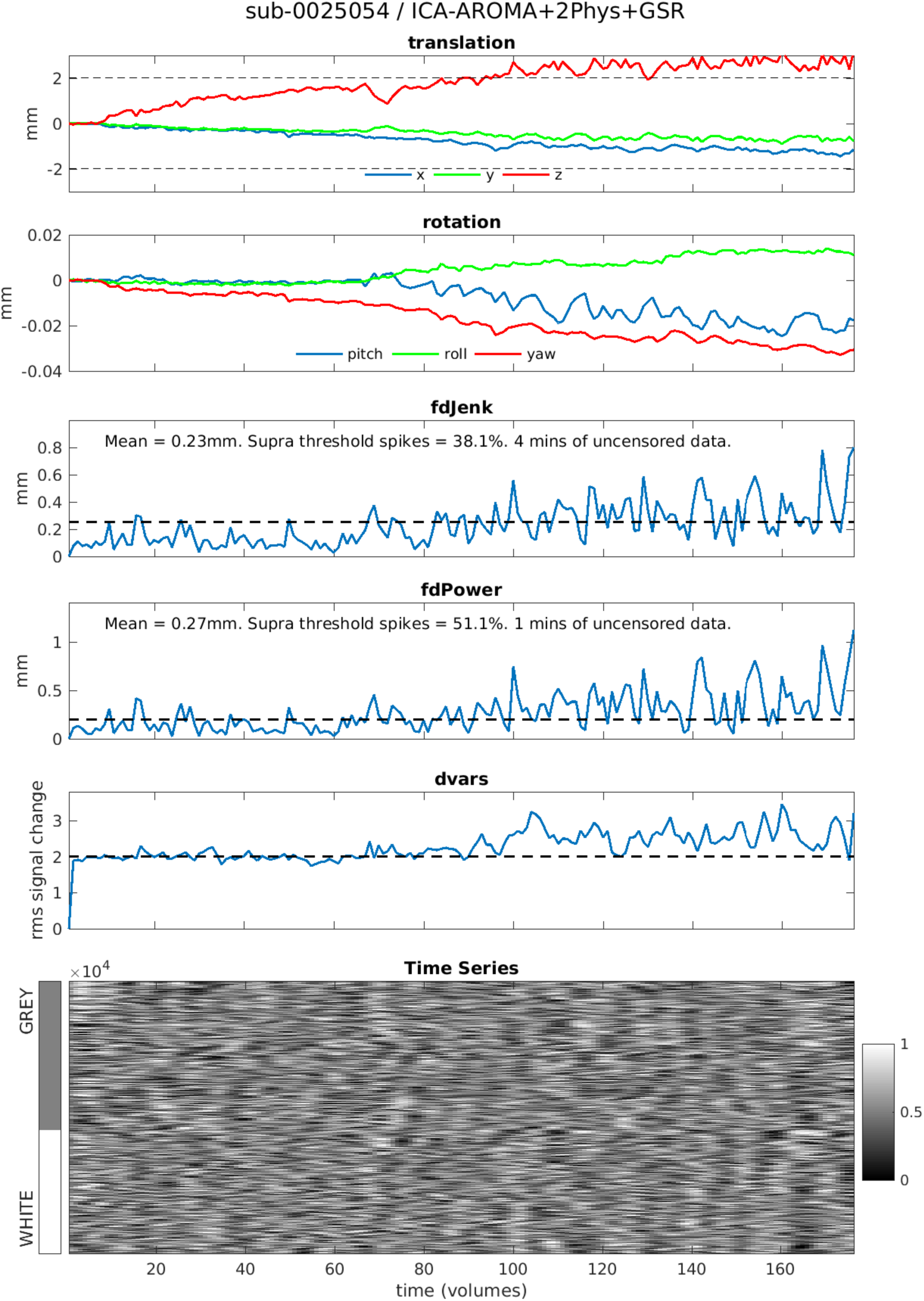
*Highest motion* participant from *NYU* dataset processed using *ICA-AROMA+2Phys+GSR*

**Figure S21.**
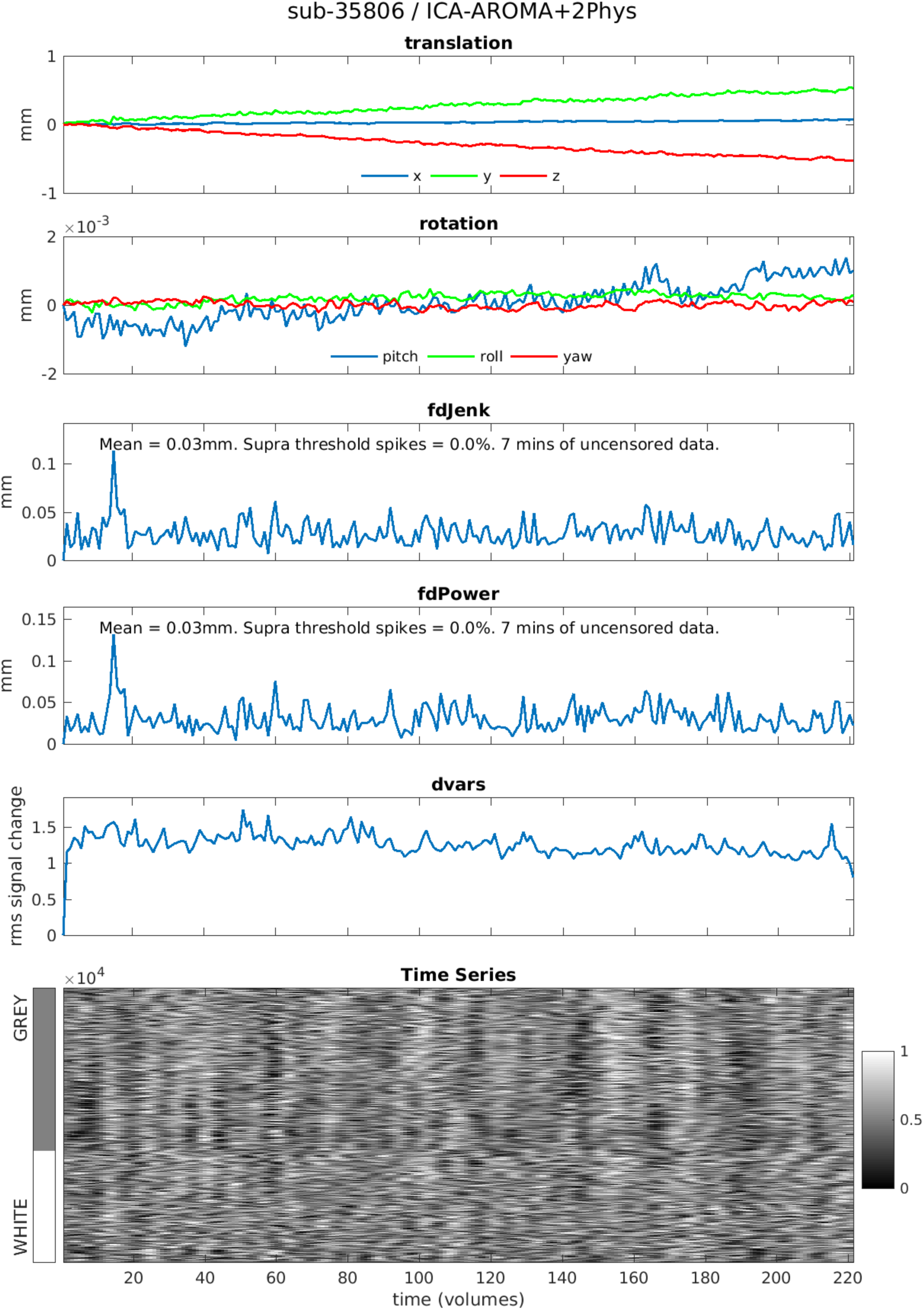
*Lowest motion* participant from *Beijing* dataset processed using *ICA-AROMA+2Phys*

**Figure S22.**
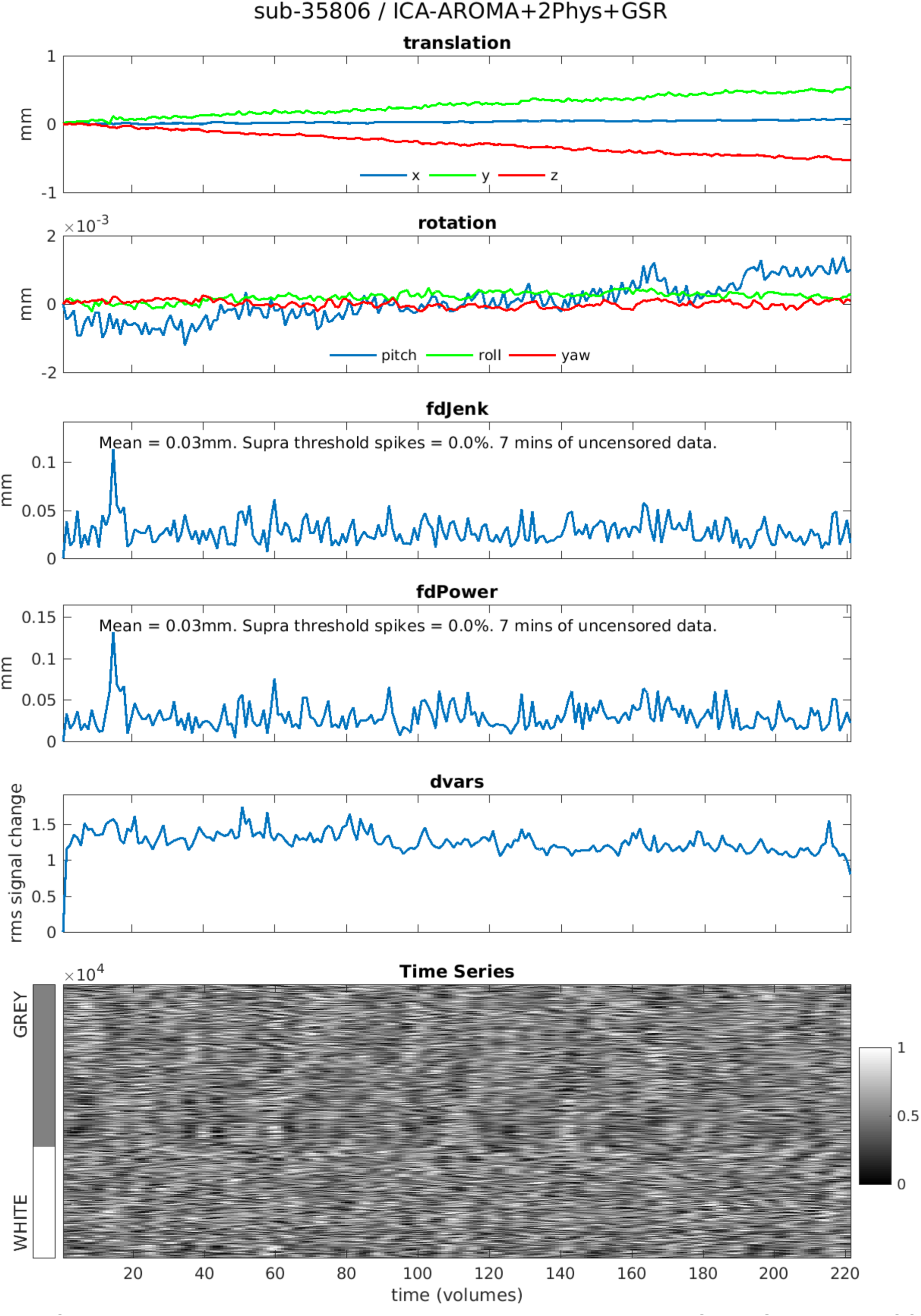
*Lowest motion* participant from *Beijing* dataset processed using *ICA-AROMA+2Phys+GSR*

**Figure S23.**
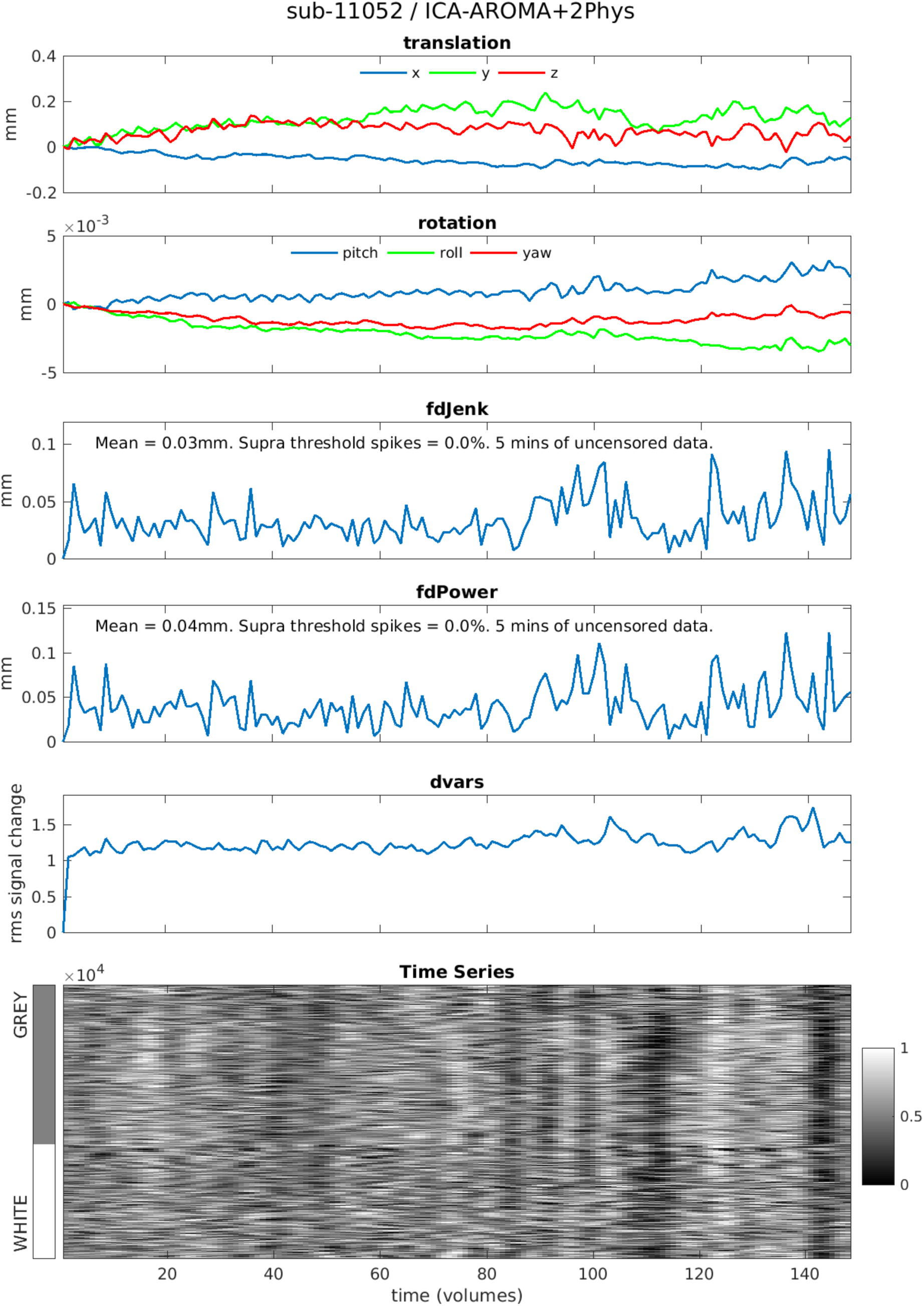
*Lowest motion* participant from *CNP* dataset processed using *ICA-AROMA+2Phys*

**Figure S24.**
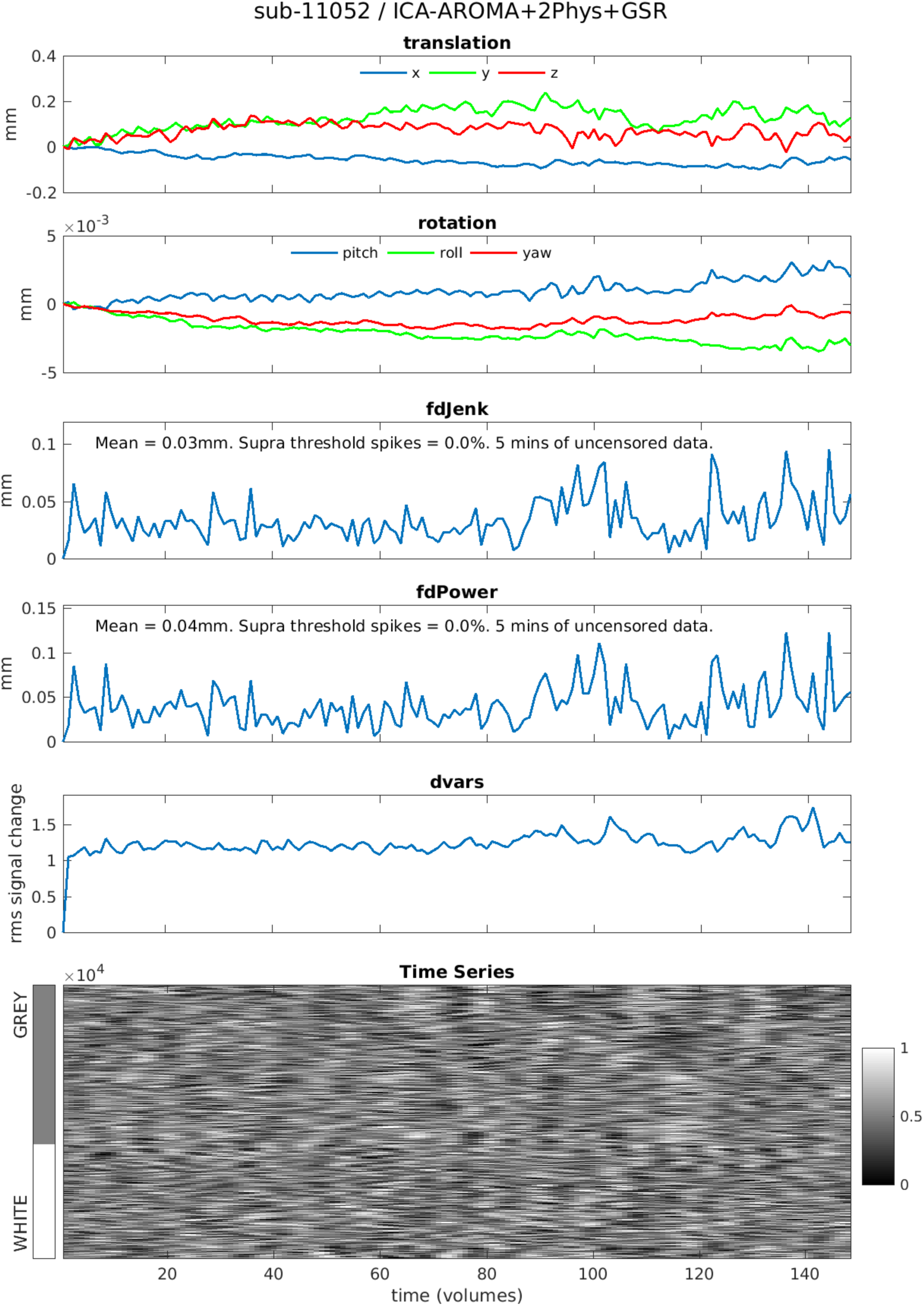
*Lowest motion* participant from *CNP* dataset processed using *ICA-AROMA+2Phys+GSR*

**Figure S25.**
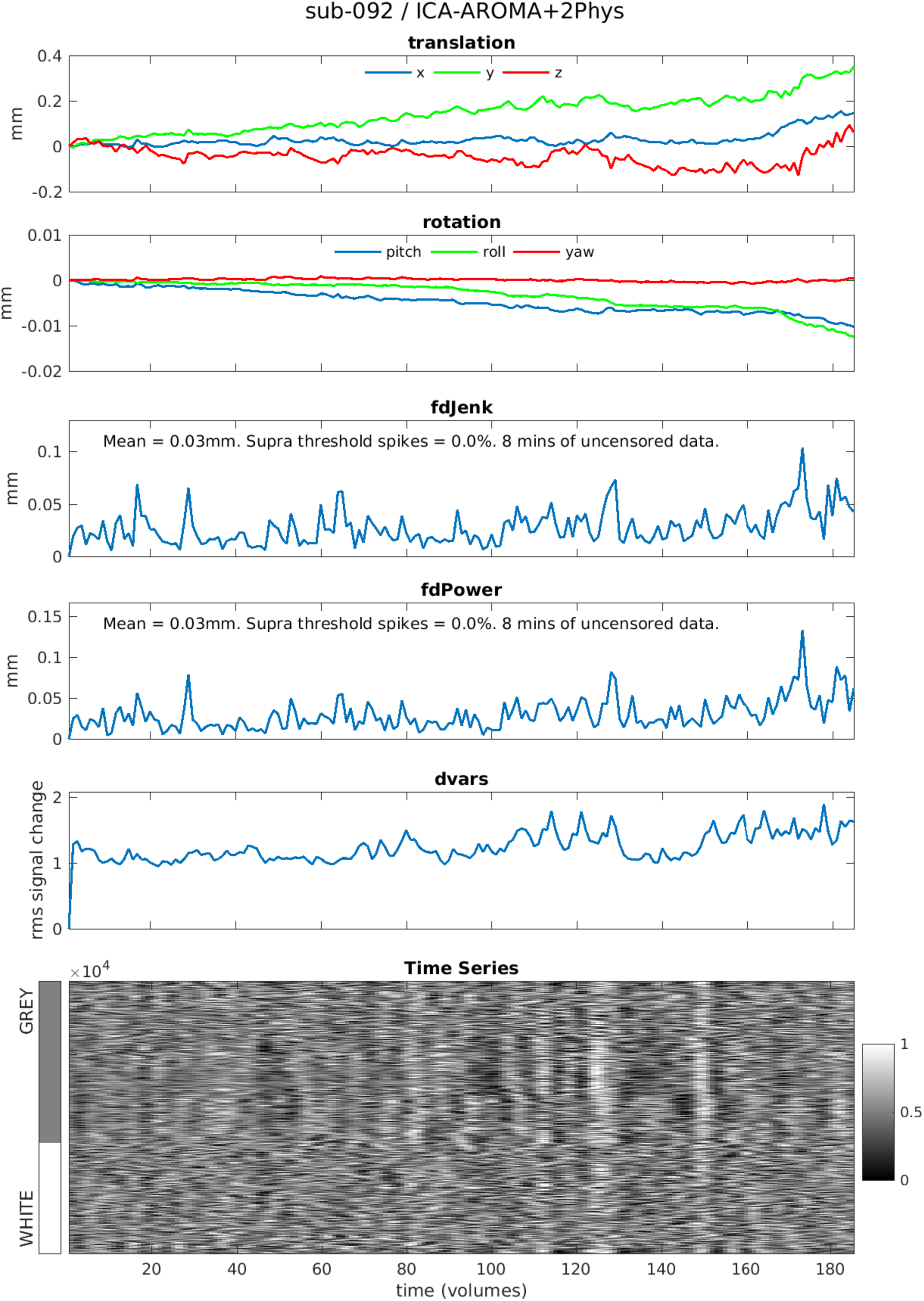
*Lowest motion* participant from *BMH* dataset processed using *ICA-AROMA+2Phys*

**Figure S26.**
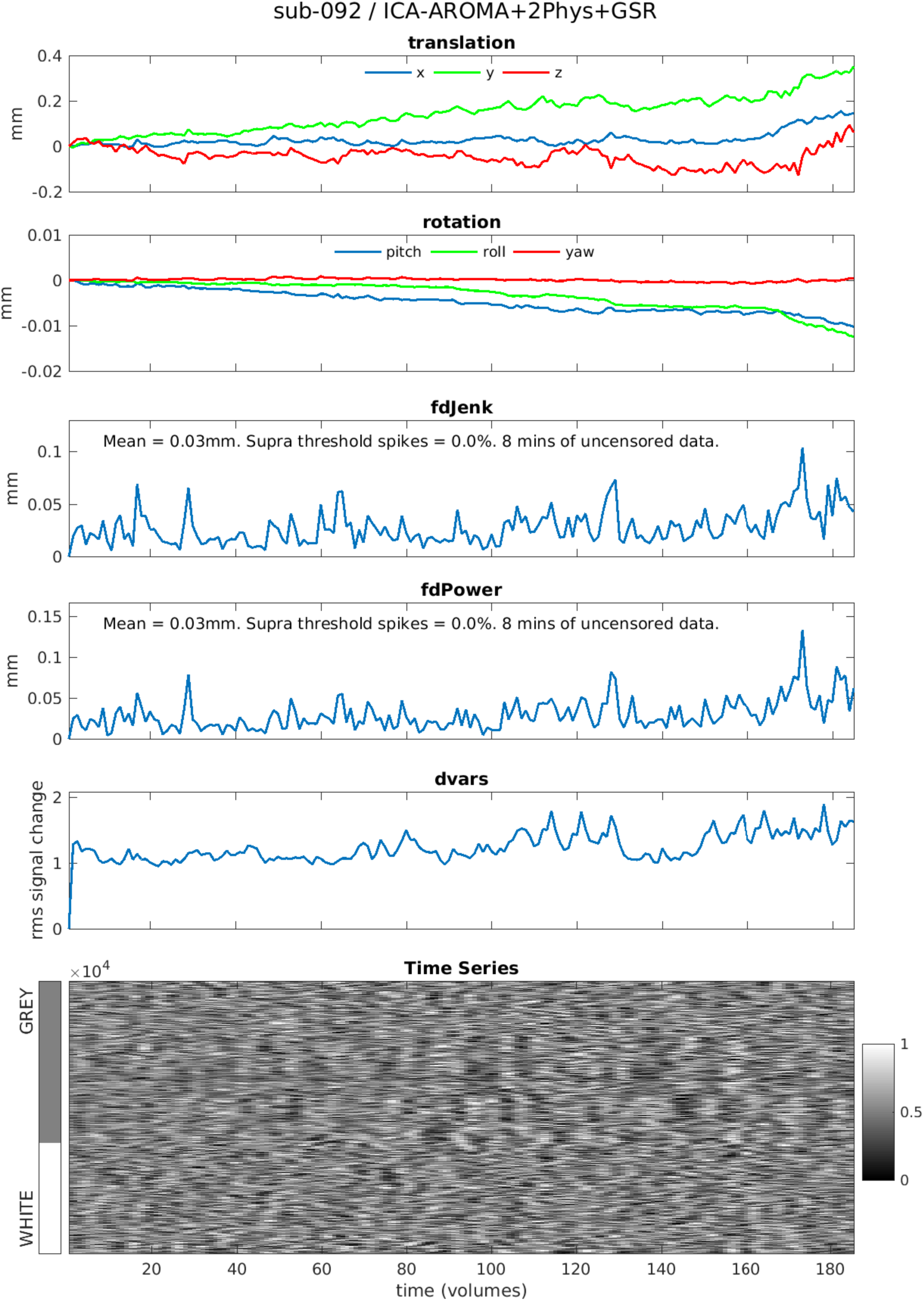
*Lowest motion* participant from *BMH* dataset processed using *ICA-AROMA+2Phys+GSR*

**Figure S27.**
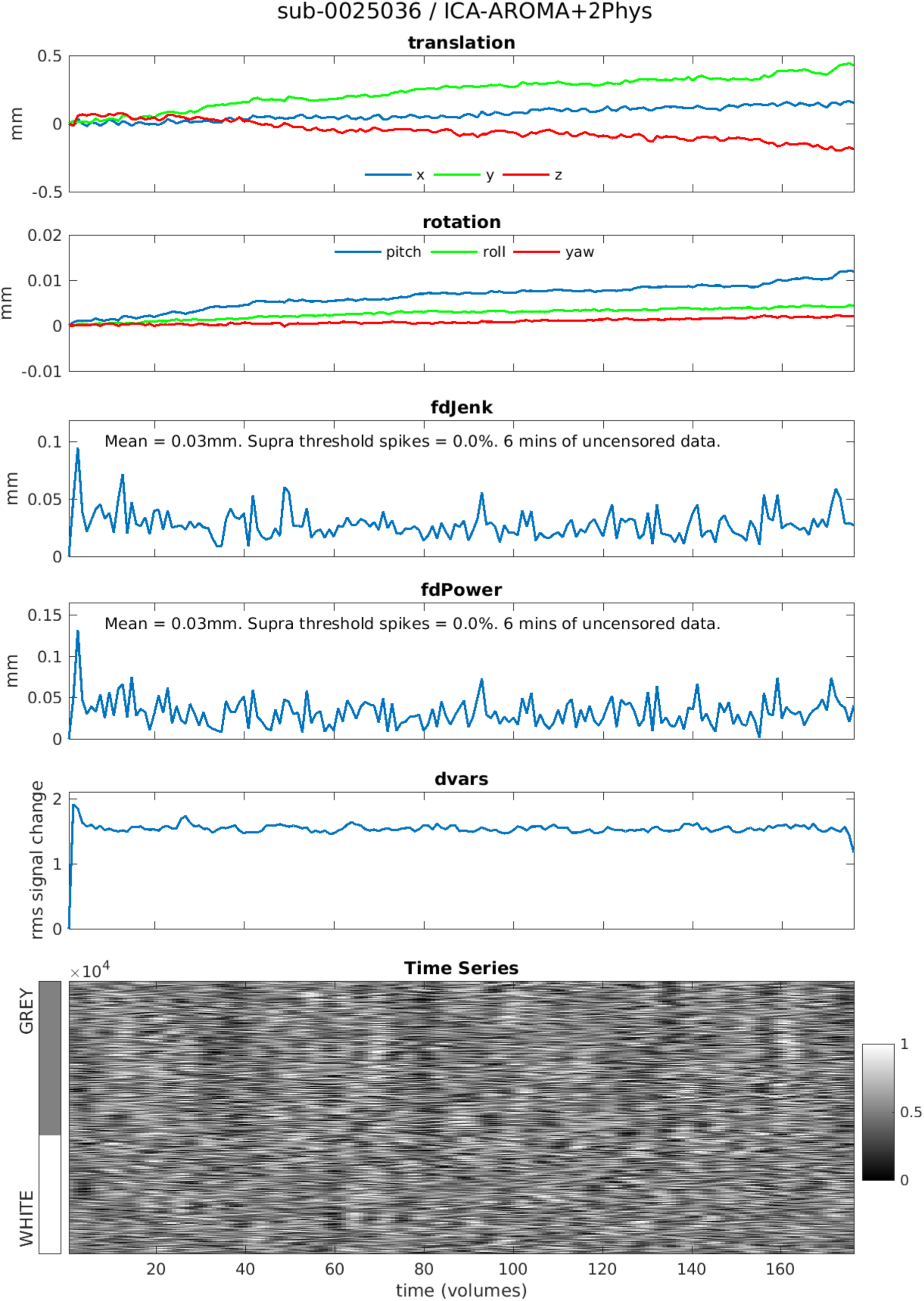
*Lowest motion* participant from *NYU* dataset processed using *ICA-AROMA+2Phys*

**Figure S28.**
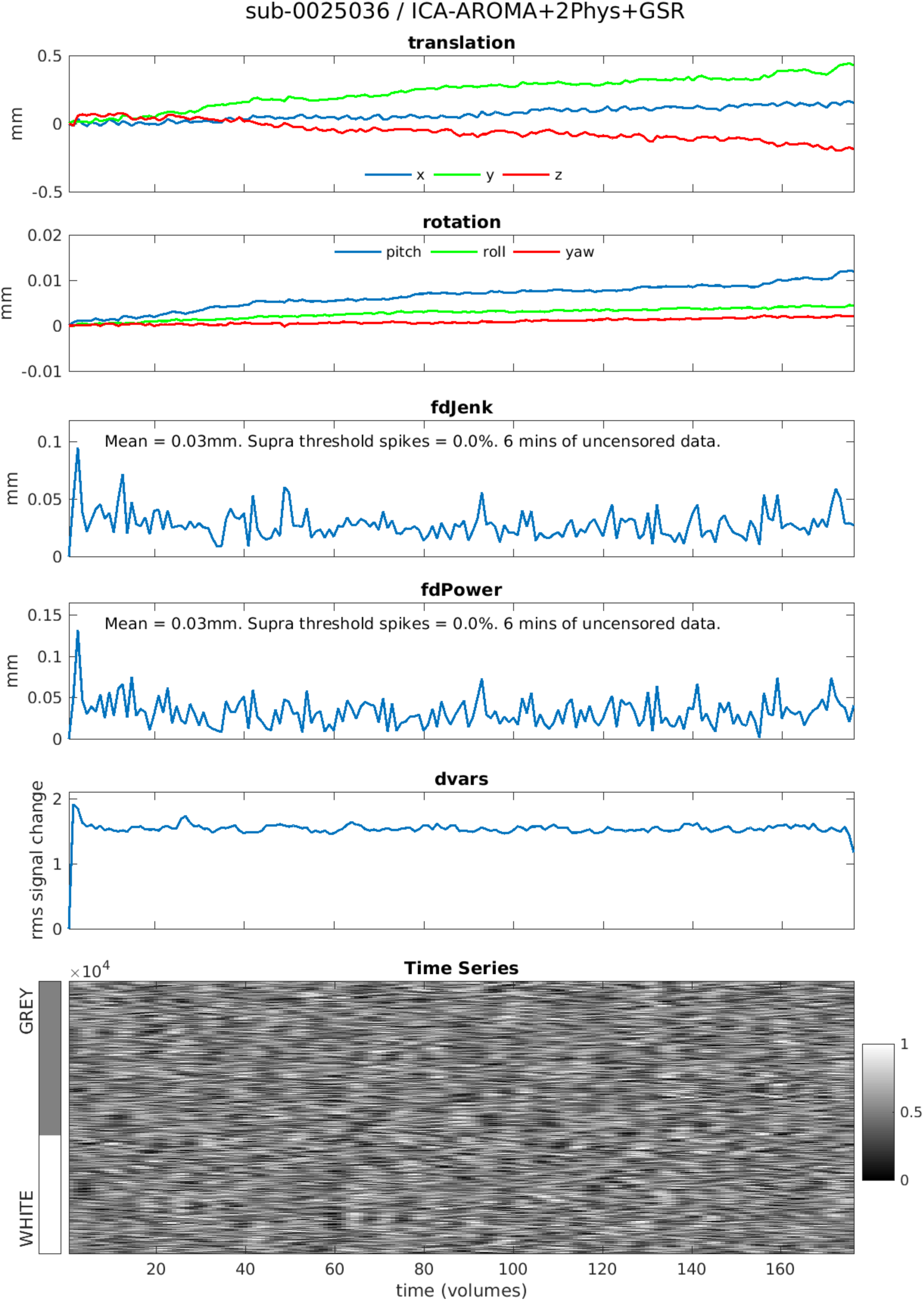
*Lowest motion* participant from *NYU* dataset processed using *ICA-AROMA+2Phys+GSR*

